# Phylogeny-driven design of broadly protective sarbecovirus receptor-binding domain nanoparticle vaccines

**DOI:** 10.1101/2025.05.11.652904

**Authors:** Amin Addetia, Alexandra Schäfer, Kaitlin Sprouse, Adian Valdez, Ashley Taylor, Mary-Jane Navarro, Jack T Brown, Elizabeth M. Leaf, Marcos C. Miranda, Alexandra C Walls, Jimin Lee, Nicholas J Catanzaro, Catherine Treichel, Isabelle Willoughby, John Powers, David R Martinez, Blue Vesari, Rashmi Ravichandran, Albert J Seo, Cameron Stewart, Benjamin Merz, Emily Beirne, Samantha Zepeda, Anthony Cook, Laurent Pessaint, Ankur Sharma, Darin Edwards, Kunse Lee, Kelly Smith, Tyler Starr, Ralph Baric, Neil P King, David Veesler

**Affiliations:** Department of Biochemistry, University of Washington, Seattle, Washington, USA; Department of Epidemiology, University of North Carolina at Chapel Hill, Chapel Hill, NC, USA; Institute for Protein Design, University of Washington, Seattle, WA 98195, USA; Department of Biochemistry, University of Utah School of Medicine, Salt Lake City, UT 84112, USA; Howard Hughes Medical Institute, Seattle, WA 98195, USA; Department of Immunobiology, Yale School of Medicine, New Haven, CT, 06511; Department of Laboratory Medicine and Pathology, Seattle, WA 98195, USA; BIOQUAL, Inc., Rockville, MD 20852, USA; Moderna Inc., Cambridge, MA 02139, USA; Department of R&D, SK Bioscience, Seongnam, Republic of Korea; Department of Microbiology and Immunology, University of North Carolina at Chapel Hill, Chapel Hill, NC, USA

## Abstract

Vaccines against emerging SARS-CoV-2 variants and sarbecoviruses with pandemic potential must elicit a robust humoral immune response in a population imprinted with the SARS-CoV-2 spike (S) protein. Here, we designed protein nanoparticle (NP) vaccines co-displaying the SARS-CoV-2 BA.5, SARS-CoV-1, and BtKY72 receptor-binding domains (RBDs) with or without the Wuhan-Hu-1 (Wu) RBD. We show that these vaccines elicit cross-reactive and broadly neutralizing plasma antibody responses against SARS-CoV-2 variants and sarbecoviruses in naive and pre-immune animals. Immunization with multivalent RBD-NPs overcomes immune imprinting and elicits neutralizing antibodies and memory B cells specific for the BA.5, SARS-CoV-1, and BtKY72 RBDs in mRNA-1273-vaccinated non-human primates. Multivalent RBD-NPs outperform a monovalent Wu RBD-NP vaccine by providing superior protection in mice and non-human primates challenged with the vaccine-mismatched SARS-CoV-2 XBB.1.5 or the pre-emergent RsSHC014. These data support the use of multivalent RBD-NP vaccines for SARS-CoV-2 variants and sarbecoviruses in naive and pre-immune populations.

## Introduction

Sarbecoviruses have twice spilled over into the human population, causing the 2002-2003 SARS epidemic (SARS-CoV-1)^1,2^ and COVID-19 pandemic (SARS-CoV-2)^3,4^. The continued circulation of SARS-CoV-2 in the human population has led to the emergence of viral variants with S glycoprotein mutations that evade neutralizing antibodies^5–12^.

Furthermore, several sarbecoviruses circulating in bat reservoirs can utilize human angiotensin-converting enzyme 2 (ACE2) as an entry receptor either directly or after accumulating a limited number of mutations in the viral spike (S) glycoprotein, suggesting possible zoonotic spillovers^13–18^. These observations underscore the necessity to develop vaccines that induce broadly protective humoral immune responses against pathogens of the *Sarbecovirus* subgenus.

SARS-CoV-2 vaccines encode or comprise the S glycoprotein to elicit robust neutralizing antibody responses^19–23^. The initial SARS-CoV-2 vaccines were developed based on the SARS-CoV-2 Wuhan-Hu-1 (Wu) S sequence and elicited potent vaccine-matched neutralizing antibody responses, which led to robust protection against COVID-19^24–28^. However, these vaccines do not elicit potent neutralizing antibody responses against vaccine-mismatched sarbecoviruses found in wildlife or immune-evasive SARS-CoV-2 Omicron variants^7,12,14,29–31^. Updated SARS-CoV-2 vaccines with the S glycoprotein of circulating variants^32–36^, elicit stronger neutralizing antibody responses against Omicron variants than Wu S-based vaccines^32–38^, but their effectiveness has been hampered by immune imprinting.

Immune imprinting, sometimes called original antigenic sin, describes a phenomenon in which a pre-existing immune response against an antigen prevents the development of a *de novo* humoral immune response towards related but distinct antigens upon subsequent exposure^39^. Most individuals who received several doses of SARS-CoV-2 Wu S-based vaccines did not develop *de novo* humoral immune responses after receiving an Omicron S-based vaccine or experiencing an Omicron infection^30,37,38,40–45^. Instead, these updated vaccines recalled pre-existing, cross-reactive memory B cells induced by Wu S and led to a modest increase of plasma neutralizing activity against Omicron variants^37,38,41,44^. Broadly protective sarbecovirus vaccines must overcome immune imprinting and elicit potent neutralizing antibody responses in Wu S-imprinted individuals.

We previously designed a vaccine displaying the receptor binding domain (RBD) of the SARS-CoV-2 Wu S glycoprotein on the icosahedral I53-50 nanoparticle (NP)^29,46–48^. Wu RBD-NP is safe and highly immunogenic in humans, leading to its licensure under the brand name SKYCovione^49,50^, which is the world’s first computationally designed medicine. This nanoparticle is de-risked for use in humans and highly versatile, enabling the display of other sarbecovirus or MERS-CoV RBDs^29,51^, making it an ideal vaccine platform for pandemic preparedness. Previously described vaccines induce neutralizing activity in animal models with varying levels of breadth across sarbecoviruses^29,52–66^.

However, few of them elicit neutralizing antibody responses against mismatched viruses and none of them outperform already existing vaccines against SARS-CoV-2 variants.

Here, we used a phylogeny driven approach to design next-generation sarbecovirus RBD-NP vaccines co-displaying multiple antigenically distinct RBDs on the I53-50 nanoparticle to elicit broad neutralizing activity and confer protection against SARS-CoV-2 variants and sarbecovirus in immunologically naive and pre-immune animal models.

## Results

### Design of multivalent RBD NP vaccines

To design a multivalent RBD-NP vaccine addressing the continued evolution of SARS-CoV-2 and sarbecoviruses with spillover potential, we selected one RBD antigen from each sarbecovirus clade with demonstrated ability to infect humans cells: SARS-CoV-1 from clade 1a, SARS-CoV-2 Wu from clade 1b, and BtKY72 from clade 3^14,15,18,67,68^. We also included in a tetravalent vaccine the SARS-CoV-2 Omicron BA.5 RBD, as it was a part of the formulation for the approved Spikevax and Cominarty bivalent Wu/BA.5 mRNA vaccines^69,70^. The SARS-CoV-2 Wu RBD was excluded from a trivalent RBD-NP to assess the impact on neutralization breadth and imprinting. We genetically fused the SARS-CoV-2 Wu, SARS-CoV-2 BA.5, SARS-CoV-1 and BtKY72 RBDs to the trimeric I53-50A nanoparticle component and recombinantly produced and purified these proteins from Expi293F cells. In vitro assembly of these RBD-I53-50A fusions with the pentameric I53-50B generated the two-component I53-50 icosahedral nanoparticle (NP), which displays sixty1 RBDs in a highly immunogenic array at its surface^29,47,51,71–73^ and was previously used for the monovalent SKYCovione^49,50^ Wu RBD vaccine.

We assembled trivalent mosaic RBD-NPs (mRBD-NPs) co-displaying the BA.5, SARS-CoV-1, and BtKY72 RBDs and tetravalent mRBD-NPs co-displaying the Wu, BA.5, SARS-CoV-1, and BtKY72 RBDs by adding I53-50B pentamers to an equimolar mixture of the appropriate RBD-I53-50A trimers (**Figure 1A**). We opted for mosaic formulations that display multiple antigens on a single NP because mosaic NPs have been hypothesized to preferentially elicit cross-reactive antibodies by favoring activation of B cells that interact with two divergent, neighboring antigens co-displayed on a NP^52,53,74,75^. We also prepared monovalent Wu and BA.5 RBD-NPs as benchmark immunogens (**Figure 1A**). After assembly, size exclusion chromatography (SEC), dynamic light scattering (DLS), and electron microscopy of negatively stained samples (nsEM) revealed that each RBD-NP homogeneously assembled to the target icosahedral architecture (**Figure 1B-D**). We confirmed the antigenic integrity of each assembled RBD-NP by demonstrating binding to a panel of monoclonal antibodies, including the SARS-CoV-2 Wu-specific LY-CoV555^76^, SARS-CoV-1-specific 80R^77^, and BtKY72-specific SK26, and ACE2 orthologs, including human and mouse ACE2s (**Figure 1E**). We demonstrated that the trivalent and tetravalent mRBD-NPs co-displayed RBDs from multiple sarbecoviruses using sandwich BLI in which the RBD-NPs were first loaded on SK26-coated biosensors tips and then dipped into 80R (**Figure 1F**). Together, these data confirm the production of monodisperse mosaic RBD-NP immunogens displaying clade 1a, clade 1b, and clade 3 sarbecovirus RBDs.

**Figure 1.**
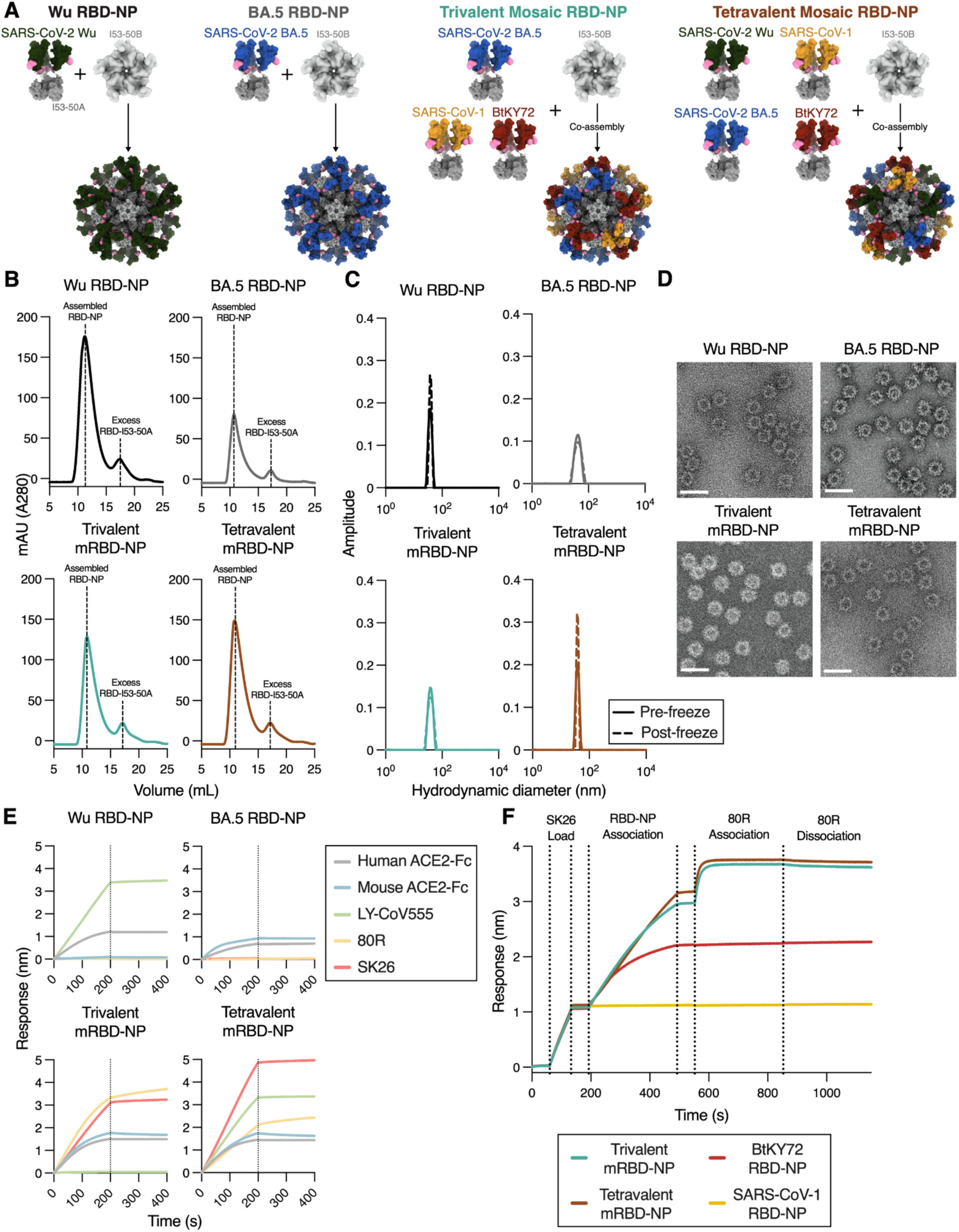
Computationally designed nanoparticle vaccines co-displaying multiple sarbecovirus RBDs. **A)** Schematic of the nanoparticles (NPs) designed in this study. **B)** Size-exclusion chromatograms for the assembled Wu, trivalent, and tetravalent mRBD-NPs. **C)** Dynamic light scattering of assembled NPs pre-freeze (solid line) and post-freeze (dashed line). **D)** Representative electron micrograph of negatively stained assembled NPs. Scale bar: 50 nm. **E)** Binding of the assembled NPs at a concentration of 100 nM to ACE2-Fc or IgGs immobilized on the surface of Protein A biolayer interferometry biosensors. **F)** Sandwich biolayer interferometry evaluating RBD-NP binding to the BtKY72-specific SK26 IgG-coated biosensor tips and subsequent binding to the SARS-CoV-1-specific 80R IgG to confirm mosaic co-display of distinct RBDs.

### Multivalent RBD-NPs elicit potent neutralizing antibody responses against vaccine-matched and -mismatched sarbecoviruses in naive mice

To assess the immunogenicity of the designed multivalent RBD-NPs, we vaccinated immunologically naive mice (n = 20 per group) with two 1 µg (RBD antigen) doses, spaced three weeks apart, of monovalent Wu RBD-NP, monovalent BA.5 RBD-NP, trivalent mRBD-NP, or tetravalent mRBD-NP adjuvanted with AddaVax (**Figure 2A**). We collected sera from these mice two weeks after the second dose and assessed neutralization potency against VSV pseudotyped with a panel of vaccine-matched and -mismatched SARS-CoV-2 variant or sarbecovirus S glycoproteins (**Figure 2B; Figure S1)**. Immunization with Wu RBD-NP elicited potent neutralizing antibody titers against the vaccine-matched SARS-CoV-2 Wu-G614 (geometric mean titer [GMT]: 6526), but not against vaccine-mismatched SARS-CoV-2 Omicron BA.5, BQ.1.1, SARS-CoV-1, BtKY72, and Khosta1 (GMT: 48, 33, 35, 34, and 33, respectively), consistent with previous data^29^. Naive mice vaccinated with monovalent BA.5 RBD-NP had potent serum neutralizing activity against BA.5 (GMT: 1852) and appreciable neutraliation of BQ.1.1 (GMT: 397), but not Wu-G614, SARS-CoV-1, or BtKY72 (GMT: 33, 37, and 33, respectively) (**Figure 2B**). In contrast, sera from mice immunized with the trivalent mRBD-NP exhibited potent neutralizing activity against the vaccine-matched BA.5, SARS-CoV-1, and BtKY72 (GMT: 624, 3108, and 1018, respectively), appreciable neutralization of the vaccine-mismatched BQ.1.1 and Khosta1 (GMT: 166 and 136, respectively) and no neutralizing activity against Wu-G614 (GMT: 36). Tetravalent mRBD-NP vaccination elicited potent neutralizing antibody titers against all four vaccine-matched viruses, Wu-G614, BA.5, SARS-CoV-1, and BtKY72 (GMT: 1165, 549, 2261, and 846, respectively), and appreciable neutralization of BQ.1.1 and Khosta1 (GMT: 239 and 194, respectively). None of the 20 Wu RBD-NP-immunized mice exhibited any detectable neutralization against the divergent SARS-CoV-2 XBB.1.5, a virus they were subsequently challenged with. In contrast, 16 of the 20 BA.5 RBD-NP-immunized mice, 11 of the 20 trivalent mRBD-NP-immunized mice, and 9 of the 20 tetravalent mRBD-NP-immunized mice had detectable neutralization of SARS-CoV-2 XBB.1.5 **(Figure S1)**. These data demonstrate that inclusion of multiple, antigenically distinct RBDs in NP vaccines elicits robust and broad polyclonal serum antibody responses against SARS-CoV-2 variants and sarbecoviruses in immunologically naive mice.

**Figure 2.**
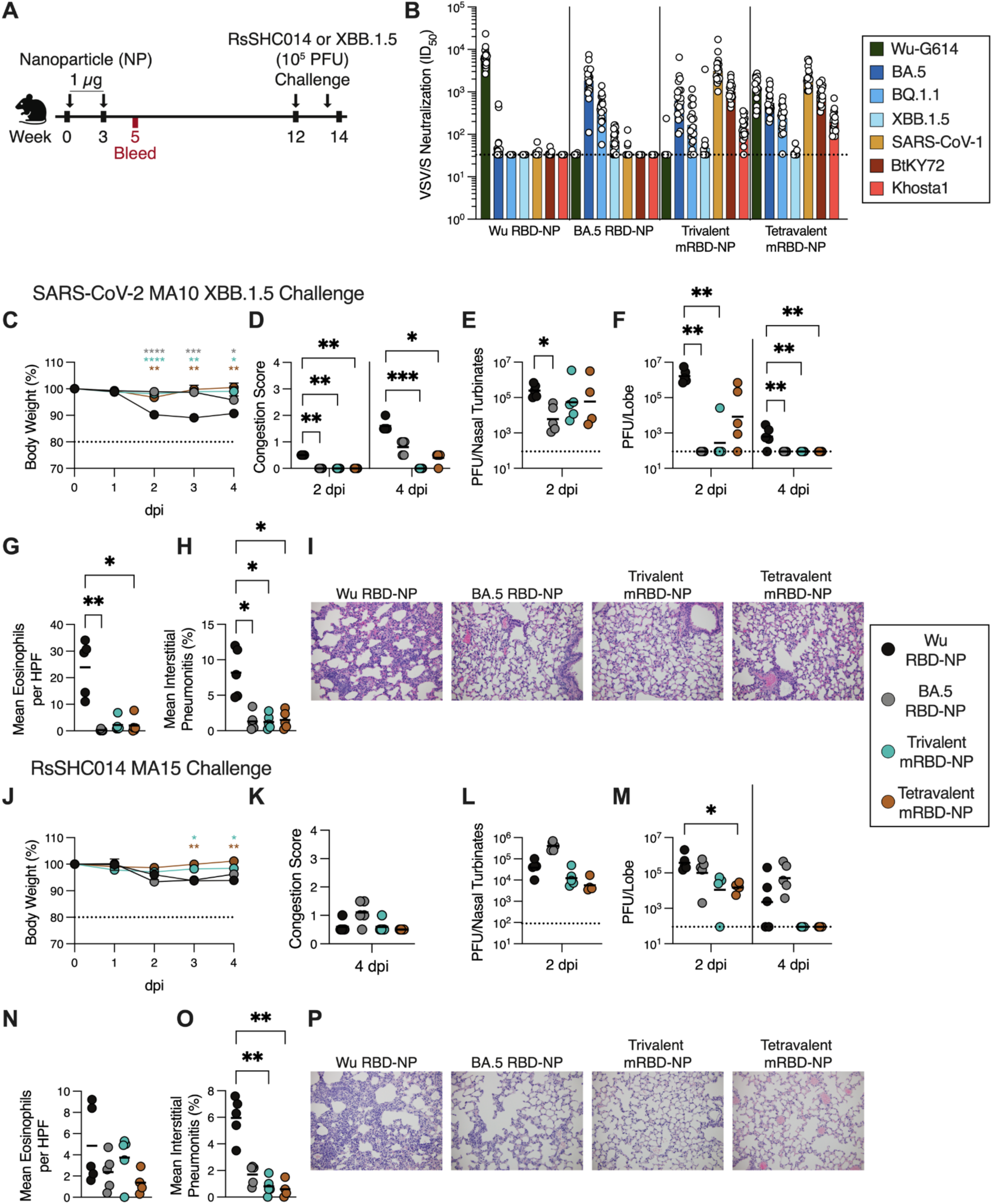
Mosaic RBD-NP vaccination elicits potent, broad and protective antibody responses in naive mice. **A)** Schematic of the immunization and challenge study design for naive mice. **B)** Serum neutralizing antibody titers measured against vaccine-matched and - mismatched SARS-CoV-2 variants and sarbecoviruses using VSV pseudotyped with the indicated S glycoprotein. Data points are presented as average ID50 values obtained from at least two biological replicates corresponding to independently produced batches of pseudovirus. The limit of detection (ID50: 1/33) is represented with a dashed line and the bars correspond to geometric mean titers (GMTs). **C)** Change in weight loss measured up to 4 days post infection (dpi) with SARS-CoV-2 MA10 XBB.1.5. Data points represent the mean body weight and error bars represent the standard error (n = 10 mice per group). **D)** Lung congestion scores measured 2 and 4 dpi with SARS-CoV-2 MA10 XBB.1.5. A score of 0 indicates that no color change of the lungs was observed and a score of 4 indicates that darkened and diseased lungs were observed. **E-F)** SARS-CoV-2 MA10 XBB.1.5 viral loads expressed as plaque forming units (PFU) per nasal turbinate 2 dpi (E) and per lung lobe 2 and 4 dpi (F). **G)** Mean eosinophils per high power field (HPF) per animal at 4 dpi for SARS-CoV-2 MA10 XBB.1.5 challenge. **H)** Mean interstitial pneumonitis per animal at 4 dpi for SARS-CoV-2 MA10 XBB.1.5 challenge. **I)** Representative images of stained lung sections from SARS-CoV-2 MA10 XBB.1.5-challenged mice immunized with the indicated vaccines at 4 dpi. **J)** Change in weight loss measured 0 to 4 dpi with RsSHC014 MA15. Points represent the mean body weight and error bars represent the standard error (n = 8-10 per group). **K)** Lung congestion scores measured 4 dpi for RsSHC014 MA15 challenge. **L-M)** RsSHC014 MA15 viral titers expressed as plaque forming units (PFU) per nasal turbinate 2 dpi (L) and per lung lobe 2 and 4 dpi (M). **N)** Mean eosinophils per high power field (HPF) per animal at 4 dpi for RsSHC014 MA15 challenge. **O)** Mean interstitial pneumonitis per animal at 4 dpi for RsSHC014 MA15 challenge. **P)** Representative images of stained lung sections from RsSHC014 MA15-challenged mice immunized with the indicated vaccines at 4 dpi. The limit of detection for viral loads (9 x 10^1^ PFU) is represented by a dotted line. The mean congestion score, geometric mean viral load, group mean of mean eosinophils per HPF, or group mean of mean interstitial pneumonitis is represented by a black bar.

### Multivalent RBD-NP vaccination protects against heterotypic challenge in naive mice

To evaluate vaccine-elicited *in vivo* protection against mismatched sarbecovirus challenge, 8-10 mice from each group were administered 10^5^ pfu of SARS-CoV-2 MA10 XBB.1.5 or of RsSHC014 MA15^16,78,79^ intranasally 10.5 or 9 weeks after the last NP dose, respectively (**Figure 2A**). Following SARS-CoV-2 MA10 XBB.1.5 challenge, mice vaccinated with the BA.5 RBD-NP, trivalent mRBD-NP, or tetravalent mRBD-NP experienced significantly less weight loss and had lower lung congestion scores 2 and 4 days post infection (dpi) compared to mice vaccinated with the Wu RBD-NP (**Figure 2C-D**). The BA.5 RBD-NP-immunized mice displayed significantly reduced viral titers in the nasal turbinates 2 dpi compared to the Wu RBD-NP (**Figure 2E**). Mice immunized with the BA.5 RBD-NP or trivalent mRBD-NP had no replicating virus in their lungs (except for one animal) and the tetravalent mRBD-NP group had lung viral titers approximately two orders of magnitude lower than the Wu RBD-NP group at 2 dpi (**Figure 2F**). The BA.5 RBD-NP, trivalent mRBD-NP, and tetravalent mRBD-NP-immunized mice had significantly lower viral titers, reduced eosinophilic infiltration, and significantly less interstitial pneumonitis in the lungs 4 dpi than Wu RBD-NP-immunized mice (**Figure 2F-I**). This enhanced protection against SARS-CoV-2 MA10 XBB.1.5 challenge is consistent with the serum neutralizing activity detected against XBB.1.5 for the BA.5 RBD-NP, trivalent mRBD-NP, and tetravalent mRBD-NP-immunized mice but not for the Wu RBD-NP-vaccinated mice (**Figure 2B, Figure S1)**.

Following RsSHC014 MA15 challenge, trivalent and tetravalent mRBD-NP-immunized mice exhibited reduced weight loss 3 and 4 dpi, lung viral titers 2 dpi, and interstitial pneumonitis 4 dpi compared to mice vaccinated with Wu RBD-NP (**Figure 2J-P**).

Moreover, none of the trivalent or tetravalent mRBD NP-vaccinated mice had detectable lung viral titers 4 dpi whereas three of the five Wu RBD-NP-vaccinated mice did (**Figure 2L-M**). The BA.5 RBD-NP-immunized mice displayed overall similar disease severity to the Wu RBD-NP-immunized mice following RsSHC014 MA15 challenge (**Figure 2J-P).** Collectively, these data show that the multivalent mRBD-NPs provide superior protection to a best-in-class monovalent Wu vaccine against mismatched challenge with an immune evasive SARS-CoV-2 variant and a pre-emergent sarbecovirus found in bats.

Statistical significance for changes in weight loss was determined using a mixed-effects analysis with Dunnett’s post-test comparing the Wu RBD-NP-immunized mice to the other groups.

Statistical significance for differences in lung congestion scores, viral loads, mean eosinophils per HPF, and mean interstitial pneumonitis was determined using the Kruskal-Wallis test with Dunn’s post-test comparing the Wu RBD-NP-immunized mice to the other groups. Only statistically significant differences are displayed. *p ≤ 0.05, **p ≤ 0.01, ***p ≤ 0.001, ****p ≤ 0.0001.

### Multivalent RBD-NPs elicit potent neutralizing antibody responses against vaccine-matched and -mismatched sarbecoviruses in pre-immune mice

Immune imprinting induced by prior SARS-CoV-2 infection or vaccination hinders the generation of a robust *de novo* humoral immune response against related antigens^30,37,38,40–43^, complicating COVID-19 vaccine updates. To model this pre-existing immunity, we vaccinated mice with a SARS-CoV-2 Wu S primary series consisting of two 0.3 µg doses of Moderna mRNA-1273, before immunization with two 1 µg doses of Wu RBD-NP, BA.5 RBD-NP, trivalent mRBD-NP, or tetravalent mRBD-NP adjuvanted with AddaVax in 3 week intervals (**Figure 3A**). We collected sera two weeks after each RBD-NP immunization to evaluate serum neutralizing antibody titers against vaccine-matched and -mismatched SARS-CoV-2 variants and sarbecoviruses.

**Figure 3.**
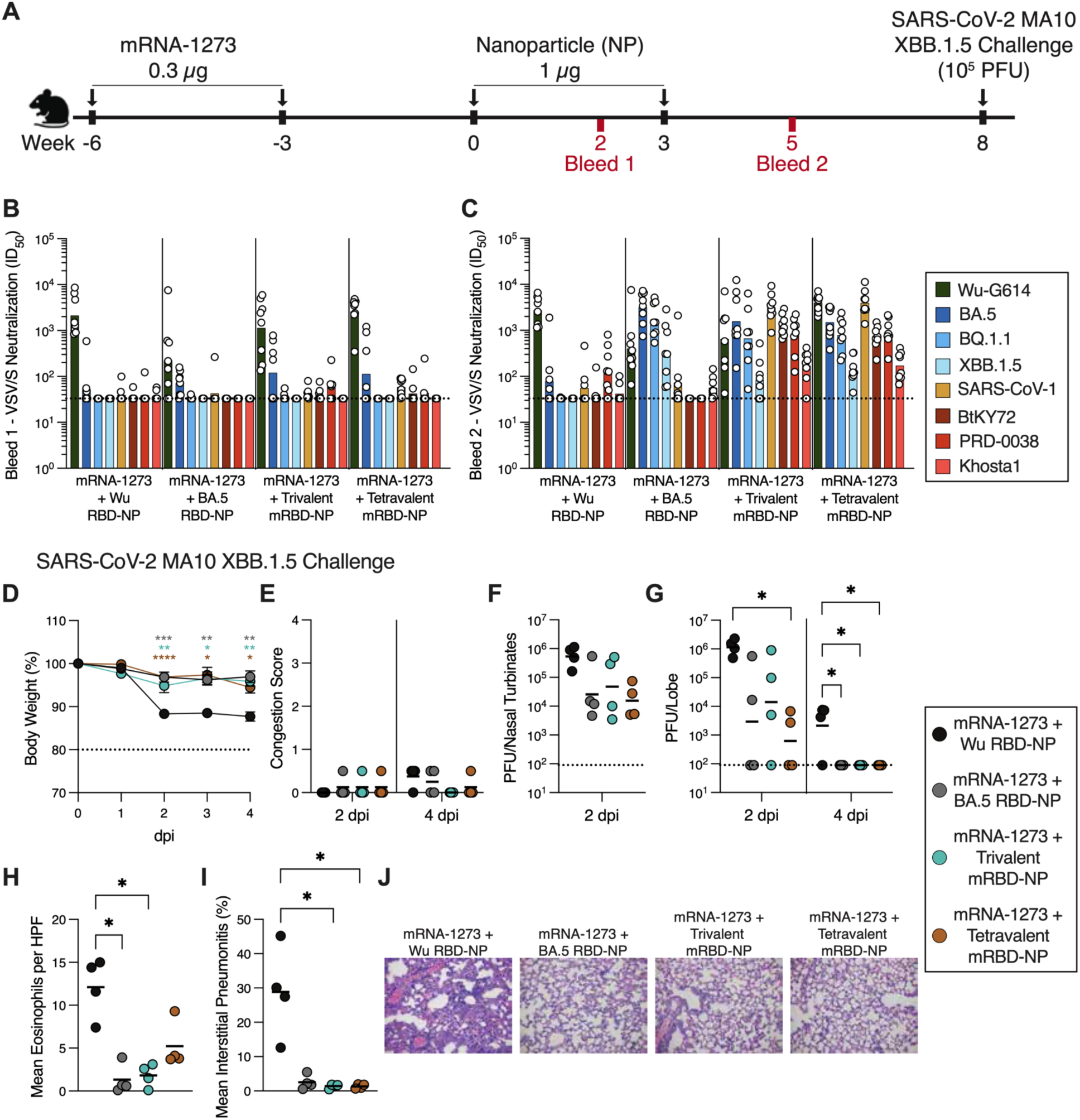
Mosaic RBD-NP vaccination elicits potent, broad and protective antibody responses in pre-immune mice. **A)** Schematic of the immunization and challenge study design for the pre-immune mice. **B-C)** Serum neutralizing antibody titers against vaccine-matched and -mismatched SARS-CoV-2 variants and sarbecoviruses using VSV pseudotyped with the indicated S glycoprotein after one dose (B) or two doses (C) of RBD-NP. Data points are presented as the average ID50 values obtained from at least two biological replicates corresponding to independently produced batches of pseudoviruses. The limit of detection (ID50: 1/33) is represented with a dashed line and the bars correspond to GMTs. **D)** Change in weight loss measured up to 4 days post infection (dpi) with SARS-CoV-2 MA10 XBB.1.5. Points represent the mean body weight and error bars represent the standard error (n = 8 per group). **E)** Lung congestion scores measured 2 and 4 dpi with SARS-CoV-2 MA10 XBB.1.5. A score of 0 indicates no color change of the lungs whereas a score of 4 indicates darkened and diseased lungs were observed. **F-G)** SARS-CoV-2 MA10 XBB.1.5 viral loads expressed as plaque forming units (PFU) per nasal turbinate 2 dpi (F) and per lung lobe 2 and 4 dpi (G). The limit of detection for viral loads (9 x 10^1^ PFU) is represented by a dotted line. **H)** Mean eosinophils per high power field (HPF) per animal at 4 dpi. **I)** Mean interstitial pneumonitis per animal at 4 dpi. **J)** Representative images of stained lung sections from mice immunized with the indicated vaccines at 4 dpi. The mean congestion score, geometric mean viral load, group mean of mean eosinophils per HPF, or group mean of mean interstitial pneumonitis is represented a black bar. Statistical significance for changes in weight loss was determined using a mixed-effects analysis with Dunnett’s post-test comparing the Wu RBD-NP-immunized mice to the other groups. Statistical significance for differences in lung congestion scores, viral loads, mean eosinophils per HPF, and mean interstitial pneumonitis was determined using the Kruskal-Wallis test with Dunn’s post-test comparing the Wu RBD-NP-immunized mice to the other groups. Only statistically significant differences are displayed. *p ≤ 0.05, **p ≤ 0.01, ****p ≤ 0.0001.

After one dose of NP, mice from all four vaccinated groups displayed modest to potent serum neutralizing activity against Wu-G614 (GMT: 2310, 222, 1139, and 2604 for Wu RBD-NP, BA.5 RBD-NP, trivalent mRBD-NP, and tetravalent mRBD-NP groups, respectively) (**Figure 3B; Figure S2)** but not against the other SARS-CoV-2 variants (BA.5, BQ.1.1, and XBB.1.5) or sarbecoviruses (SARS-CoV-1, BtKY72, PRD-0038, and Khosta1) tested. After the second NP dose, the Wu RBD-NP-immunized mice had potent serum neutralizing activity against Wu-G614 (GMT: 2734), but not against the other pseudoviruses tested (GMT: 78, 33, 33, 56, 41, 144, and 42 against BA.5, BQ.1.1, XBB.1.5, SARS-CoV-1, BtKY72, PRD-0038 and Khosta1, respectively) (**Figure 3C; Figure S2)**. BA.5-immunized mouse sera potently neutralized Wu-G614, BA.5, and BQ.1.1 (GMT: 404, 3188, and 1713 respectively) and appreciably neutralized XBB.1.5 (GMT: 330), but not SARS-CoV-1, BtKY72, PRD-0038, or Khosta1 (GMT: 73, 33, 33, and 57, respectively). Trivalent mRBD-NP vaccination elicited potent neutralizing antibody responses against Wu-G614 (GMT: 653) and all three vaccine-matched pseudoviruses (GMT: 1578, 3162, and 1310 for BA.5, SARS-CoV-1, and BtKY72, respectively). Moreover, we detected robust neutralizing activity against the vaccine-mismatched BQ.1.1 and PRD-0038 pseudoviruses (GMT: 666 and 779, respectively) and appreciable neutralization of the more divergent XBB.1.5 and Khosta1 (GMT: 83 and 172, respectively). A similar trend was observed for the tetravalent mRBD-NP-immunized mice, with potent neutralization of Wu-G614, BA.5, BQ.1.1, SARS-CoV-1, BtKY72, and PRD-0038 (GMT: 3989, 1510, 788, 3951, 706, and 894, respectively) and weaker activity against XBB.1.5 and Khosta1 (GMT: 136 and 171, respectively). The higher serum neutralizing antibody titers against Wu-G164 for the tetravalent relative to trivalent mRBD NP-immunized mice likely reflects superior boosting of Wu RBD-directed antibodies due to inclusion of this antigen in the tetravalent formulation. These results suggest that two doses of multivalent mRBD-NPs elicit robust neutralizing antibody titers against antigenically divergent SARS-CoV-2 variants and sarbecoviruses in Wu S-imprinted mice.

### Multivalent RBD-NP vaccination protects pre-immune mice against SARS-CoV-2 XBB.1.5 challenge

To assess the protective efficacy of the multivalent mRBD-NPs in mRNA-1273-imprinted mice, we challenged them with 10^5^ PFU of intranasally-administered SARS-CoV-2-MA10 XBB.1.5 five weeks after their second RBD-NP dose (**Figure 3A**). The BA.5 RBD-NP, trivalent mRBD-NP, and tetravalent mRBD-NP-immunized mice exhibited reduced weight loss 2 to 4 dpi, replicating viral titers in their lungs at 2 and 4 dpi, and reduced eosinophilic infiltration and interstitial pneumonitis in their lungs at 4 dpi compared to the Wu RBD-NP-immunized mice (**Figure 3D-J**). Three out of four Wu RBD-NP-vaccinated mice had detectable viral titers in the lungs 4 dpi, whereas none of the other RBD-NP-vaccinated mice did at this time point, underscoring the superior protection they conferred, including the monovalent BA.5 RBD-NP (**Figure 3G**).

Collectively, these results show that multivalent mRBD-NPs elicit robust and protective immune responses against an immune-evasive SARS-CoV-2 variant in Wu S-imprinted mice.

### Multivalent RBD-NPs elicit broadly neutralizing plasma antibody responses in pre-immune African green monkeys

We next assessed the immunogenicity of the multivalent mRBD-NPs in a pre-immune non-human primate model. Groups of 6 to 11 African green monkeys (AGMs) were vaccinated with two 30 µg doses of mRNA-1273 spaced one month apart to model pre-existing immunity and the resulting SARS-CoV-2 Wu S imprinting (**Figure 4A**). Eight to ten months after receiving the second dose of mRNA-1273, AGMs were vaccinated with two 40 µg (RBD antigen) doses of AddaVax-adjuvanted Wu RBD-NP, trivalent mRBD-NP, or tetravalent mRBD-NP spaced three months apart. Two weeks after the first RBD-NP immunization, AGMs from all three vaccine groups had modest to potent plasma binding titers against a wide variety of sarbecovirus prefusion S and RBD antigens (**Figure 4B-C; Figure S3; Figure S4)**. These findings were further corroborated by analyzing plasma antibody binding to a library of yeast-displayed sarbecovirus RBDs spanning all four sarbecovirus clades (**Figure 4D**). Plasma from all three vaccine groups neutralized Wu-G614 (GMTs: 497, 196, 172) whereas neutralization of SARS-CoV-2 variants (BA.5, BQ.1.1, XBB.1.5, and JN.1) and other sarbecoviruses (SARS-CoV-1, RsSHC014, BtKY72, PRD-0038, and Khosta1) was weak to undetectable (**Figure 4E; Figure S5)**, concurring with the pre-immune mouse immunogenicity data (**Figure 3B**). These data suggest that a single dose of multivalent mRBD-NPs induces humoral responses dominated by recall of Wu-imprinted antibodies, thereby limiting elicitation of potent neutralizing antibody titers against antigenically divergent SARS-CoV-2 variants or sarbecoviruses.

**Figure 4.**
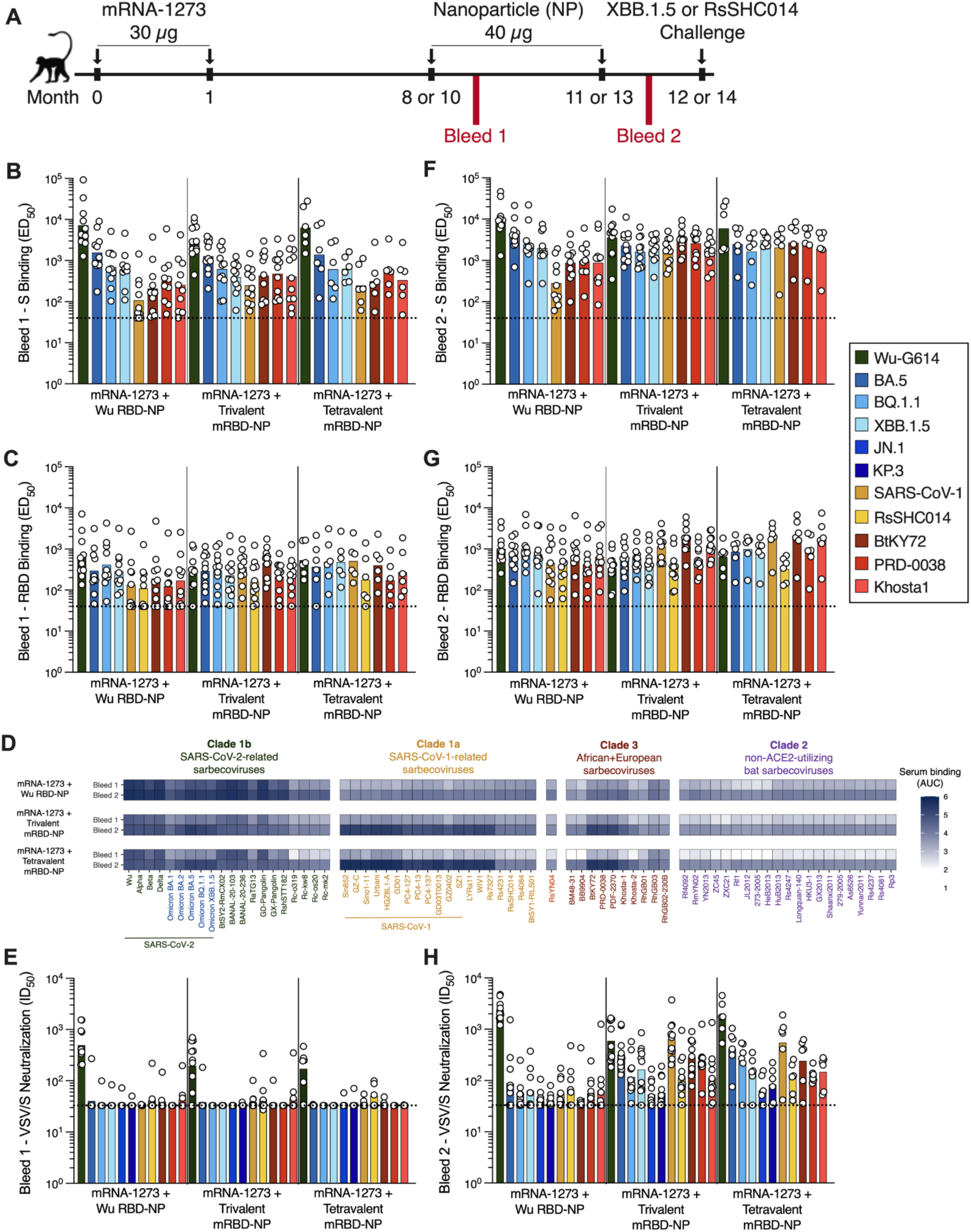
Multivalent RBD-NPs elicit cross-reactive and broadly neutralizing antibodies in mRNA-1273-imprinted African Green monkeys. **A)** Schematic of the immunization and challenge study schedule in mRNA-1273 pre-immune African Green monkeys (AGMs). **B-C)** Plasma antibody binding titers against the indicated prefusion S ectodomain trimers (B) and RBDs (C) measured by ELISA for plasma collected after one dose of RBD-NP. **D)** Binding to yeast surface-displayed sarbecovirus RBDs after one or two doses of RBD-NP. The results reflect averaged binding data from three AGMs with the highest XBB.1.5 neutralization titers per group. **E)** Neutralizing antibody titers for plasma collected two weeks after the first dose of RBD-NP against VSV pseudotyped with the indicated S glycoprotein. **F-G)** Plasma antibody binding titers against the indicated prefusion S ectodomain trimers (F) and RBDs (G) measured by ELISA for plasma collected two weeks after the second dose of RBD NP. **H)** Neutralizing antibody titers for plasma collected two weeks after the second dose of RBD-NP against VSV pseudotyped with the indicated S glycoprotein. Each data point represents the ED50 or ID50 for a given animal obtained by averaging two or more biological replicates (independently produced batches of proteins or pseudoviruses) conducted in technical duplicate. The limit of detection (ED50: 1/40 and ID50: 1/33) is represented by the dotted line. Bars indicate geometric mean titers (GMTs).

Plasma from all three vaccine groups collected after two RBD-NP doses exhibited higher binding titers against all prefusion S and RBD antigens compared to plasma collected after one RBD-NP dose (**Figure 4F-G; Figure S3; Figure S4)**. We observed greater binding to clade 1b RBDs for the Wu RBD-NP-immunized AGMs compared to trivalent or tetravalent mRBD NP-immunized AGMs using our library of yeast-displayed sarbecovirus RBDs (**Figure 4D**). In contrast, trivalent and tetravalent mRBD NP-immunized AGMs exhibited greater plasma binding for clade 1a and clade 3 RBDs compared to Wu RBD-NP-immunized AGMs. Plasma from all groups exhibited weak binding to non-ACE2-utilizing clade 2 sarbecovirus RBDs, consistent with the lack of an antigen from this clade in the vaccines. Two doses of multivalent mRBD-NPs therefore increase the magnitude and breadth of plasma binding activity towards diverse sarbecoviruses, but not for antigenically divergent SARS-CoV-2 variants, compared to the monovalent Wu RBD-NP in pre-immune AGMs.

We then examined the neutralization potency of plasma collected from AGMs that received two NP doses. Plasma from the Wu RBD-NP-immunized AGMs potently neutralized Wu-G614 (GMT: 2346) but only weakly neutralized the other pseudoviruses tested (GMT: 88, 51, 63, 44, 40, 57, 61, 46, 57, and 63 against BA.5, BQ.1.1, XBB.1.5, JN.1, KP.3, SARS-CoV-1, RsSHC014, BtKY72, PRD-0038, and Khosta1, respectively) (**Figure 4H; Figure S5)**. In contrast, the trivalent and tetravalent mRBD-NP-immunized AGMs exhibited robust plasma neutralizing titers against Wu-G614 (GMT: 593 and 1690, respectively) and the three additional vaccine-matched viruses, BA.5 (GMT: 322 and 409, respectively), SARS-CoV-1 (GMT: 639 and 549, respectively), and BtKY72 (GMT: 282 and 241, respectively). We observed dampened plasma neutralizing antibody titers against SARS-CoV-2 BQ.1.1 and XBB.1.5 for the trivalent (GMT: 117 and 165, respectively) and tetravalent (GMT: 239 and 141, respectively) mRBD-NP-immunized AGMs relative to the vaccine-matched BA.5 pseudovirus. Plasma from both groups neutralized the vaccine-mismatched RsSHC014 (GMT: 102 and 89, respectively), PRD-0038 (GMT: 237 and 126, respectively), and Khosta1 (GMT: 114 and 147, respectively) pseudoviruses and weakly neutralized the SARS-CoV-2 JN.1 (GMT: 53 and 64, respectively) and KP.3 variants (GMT: 58 and 84, respectively).

These results demonstrate that two doses of the mRBD-NPs elicit broadly neutralizing plasma antibodies against both antigenically divergent SARS-CoV-2 variants and sarbecoviruses in mRNA-1273-imprinted AGMs, outperforming the monovalent Wu RBD-NP vaccine.

### Multivalent RBD-NPs reduce SARS-CoV-2 XBB.1.5 or RsSHC014 viral titers in mRNA-1273-imprinted African Green monkeys

To evaluate vaccine protective efficacy, we challenged the pre-immune, RBD NP-immunized AGMs, along with unvaccinated AGMs, with either SARS-CoV-2 XBB.1.5 or RsSHC014 (**Figure 4A**). The RsSHC014 challenge experiment included AGMs that received the monovalent Wu RBD-NP and the trivalent mRBD-NP, while the XBB.1.5 challenge experiment additionally included animals that received the tetravalent mRBD-NP. Each animal was inoculated via the intranasal and intratracheal routes with either 10^7^ PFU (RsSHC014) or 3x10^6^ TCID_50_ (SARS-CoV-2 XBB.1.5) of virus one month after receiving their second RBD-NP dose, and viral titers were measured 1, 2, and 4 dpi in bronchio-alveolar lavages (BALs), oral swabs, and nasal swabs.

Following XBB.1.5 challenge (**Figure 5; Figure S6)**, the tetravalent mRBD-NP-immunized AGMs exhibited significantly lower viral titers in the BAL compared to the unvaccinated AGMs at 1 and 2 dpi (**Figure 5A**). Both the trivalent and tetravalent mRBD-NP-immunized AGMs displayed significantly lower viral loads in the nasal swabs than the unvaccinated AGMs at 2 dpi (**Figure 5C**). Additionally, the trivalent mRBD-NP-immunized AGMs displayed significantly lower lung consolidation than the unvaccinated animals at 4 dpi (**Figure 5D,F**). Following RsSHC014 challenge (**Figure 5; Figure S6)**, the AGMs immunized with the trivalent mRBD-NP exhibited significantly lower viral titers in the BAL and tracheal inflammation compared to the naive AGMs at 4 dpi (**Figure 5G,K,L)**. No significant differences were observed between the unvaccinated and monovalent Wu RBD-NP-immunized AGMs by any measure in these experimental conditions. These data indicate that immunization with the multivalent mRBD-NPs provides enhanced protection from challenge with heterotypic SARS-CoV-2 variants and sarbecoviruses compared to the monovalent Wu RBD-NP in pre-immune AGMs.

**Figure 5.**
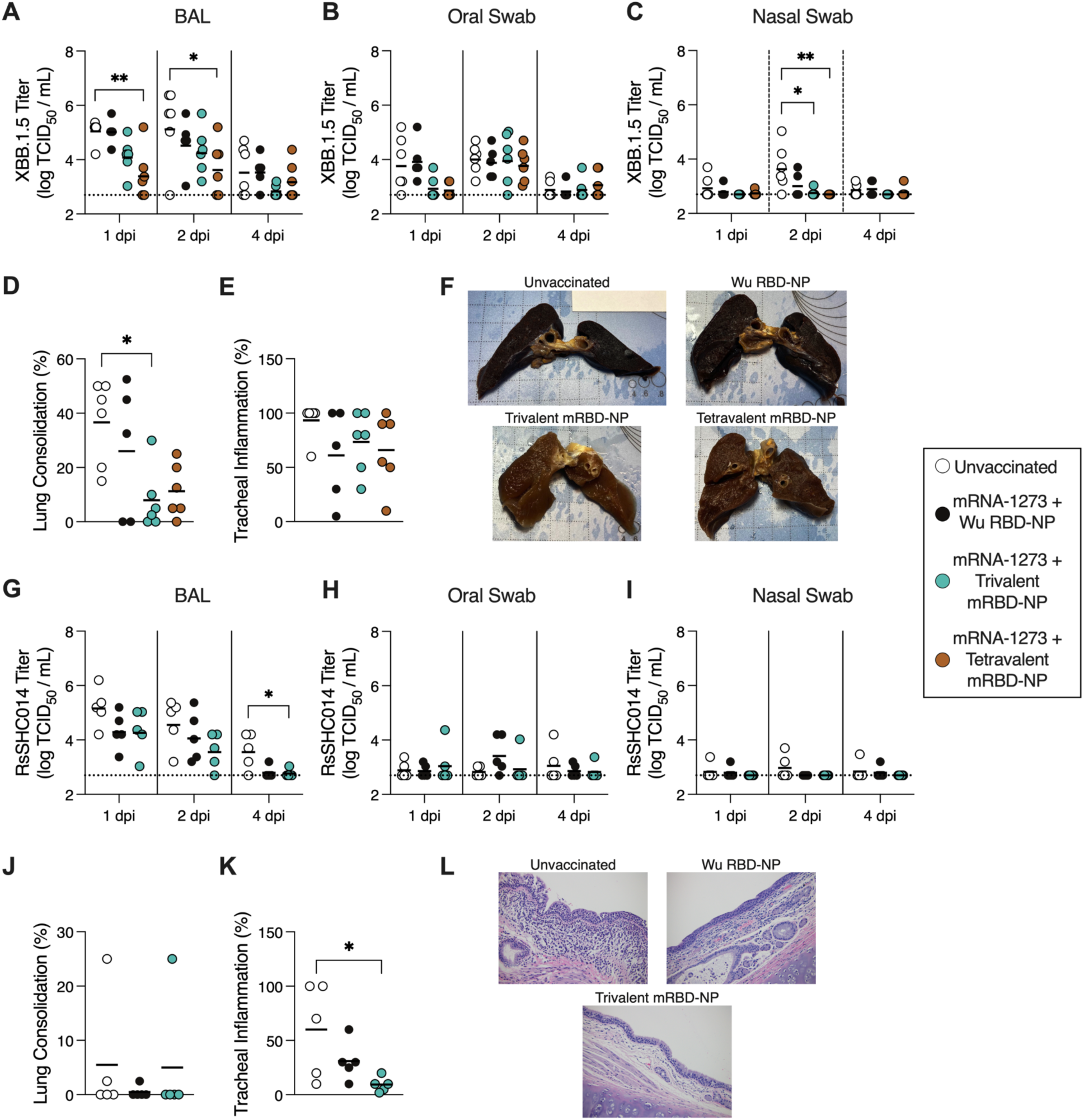
Multivalent RBD-NPs protect mRNA-1273-imprinted African Green monkeys upon challenge with the vaccine-mismatched SARS-CoV-2 XBB.1.5 and RsSHC014. A-C) Viral titers for BAL samples (A), oral swabs (B), and nasal swabs (C) collected from unvaccinated or RBD NP-immunized African Green monkeys (n = 5-6) challenged with SARS-CoV-2 XBB.1.5 at 1, 2, and 4 days post-infection (dpi). **D-E)** Lung consolidation (D) and tracheal inflammation (F) observed at 4 dpi with SARS-CoV-2 XBB.1.5. **F)** Gross lung images from AGMs challenged with SARS-CoV-2 XBB.1.5. **G-I)** Viral titers for BAL samples (G), oral swabs (H), and nasal swabs (I) collected from unvaccinated or RBD NP-immunized African Green monkeys (n = 5) challenged with RsSHC014 at 1, 2, and 4 dpi. **J-K)** Lung consolidation (J) and tracheal inflammation (K) observed at 4 dpi with RsSHC014. **L)** Representative images of stained tracheal tissue from AGMs challenged with RsSHC014. Viral titers are expressed as the tissue culture infectious dose 50 per milliliter (TCID50/mL) and the limit of detection (log[TCID50/mL]: 2.7) is represented by the dashed line. Black bars indicate the geometric mean viral titer, mean lung consolidation, or mean tracheal inflammation. Statistical significance was determined using Kruskal-Wallis test with Dunn’s post-test comparing the viral titers, lung consolidation, or tracheal inflammation of the unvaccinated AGMs to the other groups. Only statistically significant differences are displayed. *p ≤ 0.05, **p ≤ 0.01, ns: non-significant, p > 0.05.

### Multivalent RBD-NP vaccination overcomes immune imprinting in African Green monkeys

We next examined whether the mRBD-NP-elicited plasma binding and neutralizing activity against BA.5, SARS-CoV-1, and BtKY72 derived from antibodies cross-reacting with the Wu RBD or if they were specific for these three RBDs. We depleted plasma of Wu RBD-directed antibodies and examined the residual plasma binding titers against the Wu, BA.5, SARS-CoV-1, or BtKY72 RBDs along with neutralizing activity against Wu-G614, BA.5, SARS-CoV-1, and BtKY72 VSV pseudoviruses. Plasma samples from AGMs that received one dose of the Wu RBD-NP did not have detectable RBD binding or neutralizing antibody titers against any pseudovirus tested after depletion **(Figure S7A-H; Figure S8)**. After depletion of Wu RBD-directed antibodies, plasma from AGMs immunized with one dose of the trivalent or tetravalent mRBD-NP had no detectable binding to the Wu and BA.5 RBDs, modest binding to the SARS-CoV-1 (Mock depleted GMT: 330 and 587 versus Wu RBD-depleted GMT: 127 and 165 for the trivalent and tetravalent mosaic RBD NP-immunized AGMs, respectively) and BtKY72 RBDs (Mock depleted GMT: 593 and 453 versus Wu RBD-depleted GMT: 104 and 83, respectively) and weak to no plasma neutralizing activity against the four pseudoviruses tested **(Figure S7A-H; Figure S8)**. These data indicate that one dose of the mRBD-NPs is sufficient for eliciting binding antibodies specific for highly divergent sarbecoviruses, relative to the Wu S imprinting antigen, but not for more genetically closely related SARS-CoV-2 variants in mRNA-1273-vaccinated AGMs.

After two doses of NP, plasma collected from Wu RBD-NP-immunized AGMs and depleted of Wu RBD-directed antibodies did not exhibit detectable binding or neutralizing activity against any of the four viruses tested (**Figure 6A-H; Figure S8)**. The trivalent and tetravalent mRBD-NP-immunized AGMs had no detectable plasma binding activity against the Wu RBD after depletion of Wu RBD-directed antibodies, whereas potent binding titers were retained against the SARS-CoV-1 (Mock depleted GMT: 936 and 1221 versus Wu RBD-depleted GMT: 443 and 699, respectively) and BtKY72 RBDs (Mock depleted GMT: 1706 and 2431 versus Wu RBD-depleted GMT: 934 and 947, respectively) (**Figure 6A-D; Figure S8)**. Following depletion of Wu RBD-directed antibodies, BA.5 RBD binding titers were reduced (Mock depleted GMT: 492 and 729 versus Wu RBD-depleted GMT: 48 and 51, respectively) but remained detectable for both the trivalent and tetravalent mRBD NP-immunized AGMs (**Figure 6B; Figure S8)**. As expected, plasma collected from either the trivalent or tetravalent mRBD-NP-immunized AGMs did not exhibit detectable neutralizing activity against Wu-G614 following depletion of Wu RBD-directed antibodies (**Figure 6E; Figure S8)**.

**Figure 6.**
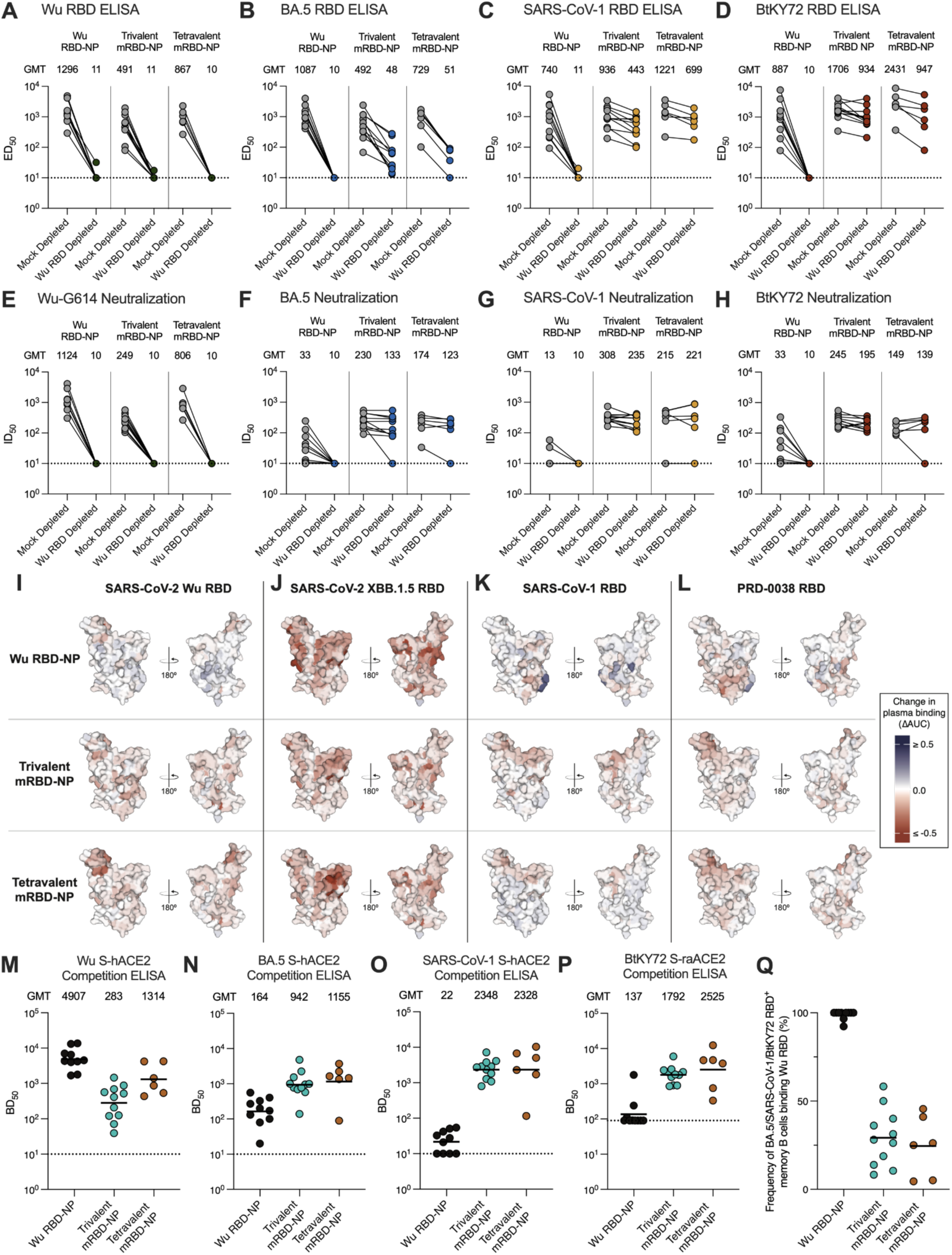
Mosaic RBD-NP vaccination overcomes Wu S-imprinting in African Green monkeys. A-D) Plasma binding titers against the (A) SARS-CoV-2 Wu, (B) SARS-CoV-2 BA.5, (C) SARS-CoV-1, or (D) BtKY72 RBDs following mock depletion or depletion of Wu RBD-directed antibodies for African Green monkeys immunized with two doses of the indicated RBD-NPs. **E-H)** Neutralizing antibody titers against VSV pseudotyped with the (E) SARS-CoV-2 Wu-G614, (F) SARS-CoV-2 BA.5, (G) SARS-CoV-1, (H) BtKY72 S following mock depletion or depletion of Wu RDD-directed antibodies for African Green monkeys immunized with two doses of the indicated RBD-NPs. One representative out of two biological replicates (each comprising technical duplicates) is shown for ELISAs and neutralization assays. The geometric mean titers (GMT) are displayed above the plots. The limit of detection (ED50: 1/10 or ID50: 1/10) is represented by a dotted line. **I-L)** Deep mutational scanning data against the SARS-CoV-2 Wu (I), SARS-CoV-2 XBB.1.5 (J), SARS-CoV-1 (K), or PRD-0038 (L) RBDs using plasma collected after two doses of RBD-NP. Data reflect averaged results from plasma collected from three AGMs with the highest ID50 values against XBB.1.5 in each group. Residue mutations that increase plasma binding are represented in blue, while those that decrease plasma binding are represented in red. **M-P)** Plasma ACE2 blocking titers for the SARS-CoV-2 Wu (M), SARS-CoV-2 BA.5 (N), SARS-CoV-1 (O), or BtKY72 (P) prefusion S trimer. Each data point represents the BD50 for a given animal obtained by averaging two biological replicates (independently produced batches of proteins) each conducted in technical duplicates. The limit of detection (BD50: 1/10 for Wu, BA.5, and SARS-CoV-1 and 1/90 for BtKY72) is represented by a dotted line. The line indicates geometric mean titers (GMTs). **Q)** Frequency of memory B cells binding to the BA.5/SARS-CoV-1/BtKY72 RBD pool that additionally recognize the Wu RBD in peripheral blood collected after two doses of the indicated RBD-NP as enumerated by flow cytometry.

Strikingly, depletion of Wu RBD-directed antibodies for the trivalent and tetravalent mosaic RBD NP-immunized AGMs did not affect neutralizing antibody titers against BA.5 (Mock depleted GMT: 230 and 174 versus Wu RBD-depleted GMT: 133 and 123, respectively), SARS-CoV-1 (Mock depleted GMT: 308 and 215 versus Wu RBD-depleted GMT: 235 and 221, respectively), and BtKY72 (Mock depleted GMT: 245 and 149 versus Wu RBD-depleted GMT: 195 and 139, respectively) (**Figure 6F-H; Figure S8)**. These data suggest two doses of the multivalent RBD-NPs are sufficient to overcome immune imprinting induced by the mRNA-1273 vaccines and elicit both binding and neutralizing antibodies specific to the antigenically distinct RBDs included in the mosaic RBD-NP vaccines.

### Plasma antibodies elicited by the multivalent RBD-NPs are directed towards epitopes recognized by potent neutralizing antibodies

To further understand how the multivalent RBD-NPs influenced the plasma antibody repertoires of mRNA-1273-imprinted AGMs, we mapped the epitopes targeted by plasma antibodies on the SARS-CoV-2 Wu, SARS-CoV-2 XBB.1.5, SARS-CoV-1, and PRD-0038 RBDs using yeast-displayed deep mutational scanning (DMS) libraries^14,80,81^. After a single RBD-NP dose, escape mutations primarily mapped to residues 484-490 within the receptor-binding motif (RBM) of the SARS-CoV-2 Wu RBD for all three vaccine groups **(Figure S7I-L; Figure S9)**, consistent with prior observations of immunodominant antibody responses to this epitope in early SARS-CoV-2 vaccination and infection cohorts^81^. Escape mutations were broadly distributed across the SARS-CoV-2 XBB.1.5 RBD for plasma from all three groups. In contrast, escape mutations were localized to the core residues on the SARS-CoV-1 RBD for plasma from Wu RBD-NP-immunized AGMs, whereas they primarily mapped to the RBM of the SARS-CoV-1 RBD for trivalent and tetravalent mRBD-NP-immunized AGMs. This result is consistent with the failure of the Wu RBD to deplete SARS-CoV-1 RBD-specific antibodies elicited by a single immunization of the mRBD-NPs **(Figure S7C)**, indicating that SARS-CoV-2 imprinting did not prevent strain-specific responses against this divergent RBD. For the PRD-0038 RBD, escape mutations localized to the core for plasma from Wu RBD-NP-immunized AGMs, whereas they were dispersed across both the RBM and core for plasma from trivalent and tetravalent mRBD-NP-immunized AGMs **(Figure S7I-L; Figure S9)**. Plasma collected from all three groups strongly competed with ACE2 binding to prefusion Wu S, moderately competed with ACE2 binding to the BA.5 S, and minimally competed with ACE2 binding to the BtKY72 S **(Figure S7M-P; Figure S10)**. Plasma from trivalent and tetravalent mRBD-NP-immunized AGMs exhibited moderate competition with ACE2 binding to SARS-CoV-1 S, while plasma from Wu RBD-NP-immunized AGMs displayed minimal competition with ACE2 binding to SARS-CoV-1 S **(Figure S7M-P; Figure S10)**.

Escape mutations from plasma collected from AGMs that received two doses of the Wu RBD-NP mapped to residues 484-490 on the SARS-CoV-2 Wu RBD, whereas they were more distributed across the RBD surface for the trivalent and tetravalent mRBD-NP groups with a hotspot on the RBM for the tetravalent RBD-NP group (**Figure 6I; Figure S9)**. Escape mutations mapped primarily to the RBM of the SARS-CoV-2 XBB.1.5 RBD for all three groups with additional enhanced relative targeting of antigenic site V (corresponding to the S2H97 cryptic epitope^84^) for Wu RBD-NP-immunized AGMs (**Figure 6J; Figure S9)**. SARS-CoV-1 and PRD-0038 RBD DMS revealed that escape mutations localized to the core residues for the Wu RBD-NP-immunized AGMs, whereas they mapped to the RBM for the trivalent and tetravalent mRBD-NP groups (**Figure 6K-L; Figure S9)**. Plasma obtained from AGMs vaccinated twice with the Wu RBD-NP strongly competed with ACE2 binding to Wu S, weakly competed with ACE2 binding to BA.5 S, and hardly competed with ACE2 binding to SARS-CoV-1 S and BtKY72 S (**Figure 6M; Figure S10)**. Conversely, plasma from the trivalent and tetravalent mRBD-NP-immunized AGMs exhibited strong competition with ACE2 binding to all four vaccine-matched prefusion S trimers (**Figure 6N-P; Figure S10)**, indicating a shift of antibody responses towards the RBM of SARS-CoV-2 variants and sarbecoviruses, which is the target of most potent neutralizing antibodies.

### Elicitation of BA.5, SARS-CoV-1, or BtKY72 RBD-specific memory B cells by multivalent RBD NPs in mRNA-1273-vaccinated AGMs

To assess how multivalent RBD-NP vaccination shapes the memory B cell repertoire, we enumerated the frequency of peripheral blood memory B cells (Single/Live/CD20^+^/CD3^-^/IgG^+^/IgD^-^/Bait^-^) recognizing the BA.5/SARS-CoV-1/BtKY72 RBD pool that also cross-reacted with the Wu RBD using flow cytometry. We analyzed memory B cells 2 weeks after both one and two NP doses for all three vaccine groups. After one and two doses, nearly all of the BA.5/SARS-CoV-1/BtKY72 RBD^+^ memory B cells identified from the Wu RBD NP-immunized AGMs cross-reacted with the Wu RBD (Mean: 96.4% and 98.9% after one and two doses, respectively) (**Figure 6Q; Figure S7Q; Figure S11)**. In contrast, the trivalent and tetravalent mRBD-NP-immunized AGMs had low frequencies of BA.5/SARS-CoV-1/BtKY72 RBD^+^ memory B cells that cross-reacted with the Wu RBD after one dose (mean: 30.5% and 25.4%, respectively) **(Figure S7Q; Figure S11)** and after two doses (mean: 29.3% and 24.6%, respectively) (**Figure 6Q; Figure S11)**. These data demonstrate that immunization with the mRBD-NPs elicits memory B cells specific to the RBDs included in the mosaic vaccines.

## Discussion

Here, we designed next-generation mosaic RBD-NP vaccines that co-display the RBDs of multiple SARS-CoV-2 variants and sarbecoviruses on the surface of the I53-50 nanoparticle^29,47^. The multivalent mRBD-NPs elicited broad sarbecovirus neutralizing activity in immunologically naive and pre-immune animals and conferred protection from vaccine-mismatched viral challenge, outperforming monovalent Wu RBD-NP and BA.5 RBD-NP vaccines. We hypothesized that mosaic co-display of the distinct sarbecovirus RBDs on NPs would preferentially activate broadly reactive B cells recognizing conserved antigenic sites^82–86^ and that the resulting antibodies would mediate broad and potent sarbecovirus neutralizing activity based on prior work on multivalent mosaic influenza hemagglutinin vaccines^74,75^. However, our depletion data suggest that most plasma neutralizing activity against SARS-CoV-2 Omicron BA.5, SARS-CoV-1, and BtKY72 derived from virus-specific antibodies rather than broadly neutralizing antibodies. These data suggest that multivalent sarbecovirus vaccines in development should include RBDs from different clades to elicit virus-and clade-specific neutralizing antibodies rather than aiming to elicit broadly neutralizing sarbecovirus antibodies.

The lack of ‘forward’ neutralization of SARS-CoV-2 Omicron variants upon Wu RBD-NP vaccination and of ‘back’ neutralization of SARS-CoV-2 Wu upon BA.5 RBD-NP vaccination in naive mice concurs with recent data on newborns infected and vaccinated with XBB.1.5^87^. Collectively, these results point to the elicitation of clade-specific immunity and suggest that SARS-CoV-2 Wu and Omicron variants constitute distinct antigenic clades^59^. The low frequency and limited potency of broadly neutralizing sarbecovirus antibodies, with few exceptions^88,89^, has thus far limited the suitability of targeting the most conserved antigenic sites to elicit high titers of polyclonal plasma neutralizing antibodies through vaccination. Hence, our data strongly support inclusion of multiple SARS-CoV-2 RBDs in vaccines developed for naive individuals to confer broad protection from circulating and emerging SARS-CoV-2 variants as well as historical isolates, particularly given to the discovery of several viruses in bats with RBD sequences similar to that of SARS-CoV-2 Wu^4,17,90^. The multivalent mRBD-NPs described here may be ideal for eliciting potent humoral immune responses against SARS-CoV-2 variants and other sarbecoviruses in newborns and infants without prior infection or vaccination.

Immune imprinting induced by prior SARS-CoV-2 Wu S exposures hinders mounting potent neutralizing antibody responses against divergent SARS-CoV-2 variants following either infection with an Omicron variant or immunization with a variant-specific booster^30,37,38,40–43^, complicating vaccine design. As most of the human population has pre-existing immunity against Wu S through infection and vaccination, we evaluated the multivalent RBD-NPs in pre-immune animals that received two doses of the widely used Moderna mRNA-1273 vaccine^19^. mRNA-1273 was selected to model pre-existing immunity against Wu S as the vaccine platform utilized can influence the severity of immune imprinting^45^. Furthermore, we focused on evaluating the impact of multivalent mRBD-NPs immunizations on immune imprinting in African Green monkeys, as mice are a less suitable model for studying imprinting due to murine germinal centers restricting re-entry of reactivated memory B cells more than human germinal centers^91,92^. In our stringent model of Wu S-imprinting, we show that depleting Wu RBD-directed plasma antibodies after one multivalent mRBD-NP immunization abrogated BA.5 plasma binding and neutralizing activity. These results point to recall of imprinted, cross-reactive antibodies, similar to observations of Wu S-imprinting in humans^38,41,44^. In contrast, depleting Wu RBD-directed antibodies from plasma collected from NHPs vaccinated twice with multivalent RBD-NPs did not appreciably impact neutralizing activity against BA.5, SARS-CoV-1 or BtKY72. This indicates that the mRBD-NPs overcame Wu S-imprinting and elicited *de novo* neutralizing antibodies specific to these RBDs. Given that the most potent neutralizing antibodies recognize the RBM^93–96^ and that our epitope mapping data show that plasma antibodies elicited by mRBD-NPs target the divergent RBMs of the antigens present in these vaccines, these results explain the mechanism underlying overcoming of immune imprinting and elicitation of robust binding and neutralizing antibody responses specific for SARS-CoV-2 Omicron BA.5, SARS-CoV-1, and BtKY72. These data support the use of the multivalent mRBD-NP for both SARS-CoV-2 variants and sarbecoviruses with spillover potential in pre-immune individuals.

Vaccines for emerging SARS-CoV-2 variants and sarbecoviruses must confer protection against vaccine-mismatched SARS-CoV-2 variants and sarbecoviruses due to the rapid rate of SARS-CoV-2 evolution and unpredictability of zoonotic spillover^9,10,15^. Our multivalent mRBD-NPs vaccines fulfill this requirement by eliciting broad plasma neutralizing activity and conferring *in vivo* protection against mismatched, immune-evasive SARS-CoV-2 variants and sarbecoviruses. The upcoming clinical advancement of these multivalent RBD-NPs could herald a new generation of “supra-seasonal” vaccines that confer protection from multiple future SARS-CoV-2 variants.

Additionally, as RBD-NPs can be delivered as mRNA vaccines^97^, our mRBD-NPs could be rapidly reformulated as needed to keep pace with the antigenic evolution of sarbecoviruses. We propose that phylogeny-driven design of multivalent vaccines for other genera and subgenera, such as the highly diverse merbecovirus subgenus^98–101^, will enable the development of broadly protective coronavirus vaccines.

## Acknowledgements

This work was funded by the National Institute of Allergy and Infectious Disease (1P01AI167966 to T.N.S., R.B., N.P.K and D.V., DP1AI158186 and 75N93022C00036 to D.V.), the Coalition for Epidemic Preparedness Innovations, FastGrants, the Bill & Melinda Gates Foundation (INV-010680 to N.P.K. and D.V.), an Investigators in the Pathogenesis of Infectious Disease Awards from the Burroughs Wellcome Fund, the National Institutes of Health (S10OD032290), a Searle Scholars Award (T.N.S.), the Audacious Project at the Institute for Protein Design, and a Hanna H. Gray Fellowship from the Howard Hughes Medical Institute (D.R.M). D.V. is an Investigator of the Howard Hughes Medical Institute and the Hans Neurath Endowed Chair in Biochemistry at the University of Washington. Research reported in this publication was generated using the DLMP Flow Cytometry Core at the University of Washington.

## Contributions

AA, MM, ACW, NPK, and DV conceived the project and designed the experiments. AV, BV, and RR expressed, assembled, and characterized the RBD nanoparticles. EML, CT, and IW performed the immunogenicity studies in mice. KRS and MJN performed the pseudovirus neutralization assays. AS, NJC, JP, DRM, and RB carried out the viral challenge studies in mice. EB and KS performed the histological analyses. AC, LP, and AS2 carried out the immunogenicity and viral challenge studies in African Green monkeys. JTB, CS, and SZ recombinantly expressed glycoproteins. KRS and AJS performed the ELISAs. AT and TS performed the deep mutational scanning and pan-sarbecovirus RBD binding assays. AA and KRS carried out the depletion assays. BM performed the competition ELISAs. AA carried out the flow cytometry analysis. DE provided unique reagents. AA, AS, TS, JL, NPK, and DV analyzed the data. AA and DV wrote the manuscript with input from all authors.

## Methods

### Cell Culture

Expi293 cells were grown in Expi293 media at 37°C and 8% CO_2_ rotating at 130 RPM or 150 RPM. Expi293 cells stably expressing the Wu-RBD-I53-50A, BA.5-Rpk9-RBD-I53-50A, SARS-CoV-1-RBD-I53-50A, or BtKY72-RBD-I53-50A were grown in Expi293 media at 37°C and 8% CO_2_ rotating at 150 RPM. HEK-293T cells and HEK-293T cells stably expressing human ACE2 (HEK-ACE2)^102^ were grown in DMEM supplemented with 10% FBS and 1% PenStrep at 37°C and 5% CO_2_. Vero cells stably expressing human TMPRSS2 (Vero-TMPRSS2)^103^ were grown in DMEM supplemented with 10% FBS, 1% PenStrep, and 8 µg/mL puromycin at 37°C and 5% CO_2_.

### Mice

For immunogenicity studies, female BALB/cAnNHsd were purchased from Envigo (order code 047) at 7 weeks of age. Mice were housed in a specific-pathogen free facility within the Department of Comparative Medicine at the University of Washington, Seattle, accredited by the Association for Assessment and Accreditation of Laboratory Animal Care (AAALAC). Animal studies were conducted in accordance with the University of Washington’s Institutional Animal Care and Use Committee (IACUC) under protocol 4470-01.

### African Green Monkeys

Thirty-eight (20 female) adult African Green Monkeys (AGMs) were sourced from PrimeGen (Lehigh Acres, FL) and housed at BIOQUAL, Inc, accredited by the AAALAC. Prior to enrollment in the studies, AGMs were determined to be free of obvious abnormalities indicative of health problems by a veterinarian and pre-screened for the absence of antibodies to SARS-CoV-2 by plaque reduction neutralization test and enzyme-linked immunosorbent assay. All studies were conducted in accordance with BIOQUAL’s IACUC under protocol 22-046.

### Constructs

Constructs encoding the SARS-CoV-2 Wu RBD fused to I53-50A and SARS-CoV-1 RBD fused to i53-50A were previously described^29,47,104^. The SARS-CoV-2 BA.5 RBD (residues 323-526) stabilized using the Rpk9 mutations^105^ and the BtKY72 RBD (residues 318-520) genetically fused using a 16-residue glycine-serine linker to I53-50A with a C-terminal octahistidine tag and an N-terminal µ-phosphatase signal peptide (MGILPSPGMPALLSLVSLLSVLLMGCVAETGT) were codon optimized, synthesized, and inserted into pCMVR by Genscript. The construct encoding I53-50B.4PT1 was previously described^104^. Constructs encoding full-length SARS-CoV-2 Wu-G614, BA.5, BQ.1.1, XBB.1.5, JN.1, SARS-CoV-1, RsSHC014, BtKY72, PRD-0038, and Khosta1 S were previously described^7,14,15,29,37,38,68,102^. The full-length SARS-CoV-2 KP.3 S lacking the final C-terminal 21 amino acids was codon optimized, synthesized, and inserted into pHDM by Genscript. Constructs encoding the SARS-CoV-2 Wu-G614, XBB.1.5, SARS-CoV-1, and PRD-0038 S ectodomain containing the HexaPro stabilizing mutations^106^ were previously described^14,38,79^. The SARS-CoV-2 BA.5 and BQ.1.1 ectodomains (residues 1-1203) containing the HexaPro stabilizing mutations^106^ and the furin cleavage site mutations (_677_RRAR_680_ to _677_GSAS_680_) along with a C-terminal T4 foldon followed by an Avi tag and octahistidine tag were codon optimized, synthesized, and inserted into pcDNA3.1(+) by Genscript. The BtKY72 S ectodomain (residues 16-1196) containing our previously described prefusion-stabilizing S_2_ subunit mutations^79^ along with an N-terminal secretion signal peptide (MGILPSPGMPALLSLVSLLSVLLMGCVA) and a C-terminal T4 foldon followed by an octahistidine tag and Avi tag was codon optimized, synthesized, and inserted into pCMVR by Genscript. The Khosta1 S ectodomain (residues 16-1189) containing the VFLIP stabilizing mutations^107^ along with an N-terminal secretion signal peptide (MGILPSPGMPALLSLVSLLSVLLMGCVA) and a C-terminal T4 foldon followed by an octahistidine tag and Avi tag was codon optimized, synthesized, and inserted into pCMVR by Genscript. Constructs encoding the SARS-CoV-2 Wu, BA.5, BQ.1.1, XBB.1.5, SARS-CoV-1, BtKY72, and PRD-0038 RBDs were previously described^7,14,15,37,68^. The RsSHC014 RBD (residues 316-518) with an N-terminal secretion signal (MGILPSPGMPALLSLVSLLSVLLMGCVA) and a C-terminal octahistidine tag followed by an Avi tag was codon optimized, synthesized, and inserted into pCMVR by Genscript. Khosta1 RBD (residues 331-531) with an N-terminal CD5 leader sequence (MPMGSLQPLATLYLLGMLVASVLA) and a C-terminal octahistidine tag followed by an Avi tag was codon optimized, synthesized, and inserted into pCMVR by Genscript. The human ACE2 ectodomain with the native dimerization domain (residues 18-740) and containing an N-terminal secretion signal (MGILPSPGMPALLSLVSLLSVLLMGCVA) and a C-terminal octahistidine tag followed by an Avi tag was codon optimized, synthesized, and inserted into pCMV by Genscript. The *Rhinolophus alcyone* ACE2 ectodomain with the native dimerization domain (residues 1-740) and a C-terminal octahistidine tag followed by an Avi tag was codon optimized, synthesized, and inserted into pCMVR by Genscript. The mouse ACE2 ectodomain (residues 19-615) fused to an N-terminal CD5 leader sequence (MPMGSLQPLATLYLLGMLVASVLA) and a C-terminal human Fc sequence was codon optimized, synthesized, and inserted into pCMV by Genscript. The heavy chain Fab sequence of SK26, a BtKY72-specific monoclonal antibody, fused to an N-terminal murine heavy chain signal peptide sequence (MGWSCIILFLVATATGVHS) and human Fc sequence was codon optimized, synthesized, and inserted into pcDNA3.1(+) by Genscript. The light chain of SK26 fused to an N-terminal murine heavy chain signal peptide sequence (MGWSCIILFLVATATGVHS) was codon optimized, synthesized, and inserted into pcDNA3.1(+) by Genscript. The human ACE2 ectodomain (residues 19-615) fused to a C-terminal human Fc (hACE2-Fc) and N-terminal BM40 signal peptide was codon optimized, synthesized, and inserted into pCMV by Genscript. The heavy and light chains of LY-CoV555 containing an N-terminal BM40 signal peptide, were codon optimized, synthesized, and inserted into pCMV by Genscript. The heavy and light chains of 80R containing an N-terminal CD5 signal peptide were codon optimized, synthesized, and inserted into pcDNA3.1(+) by Genscript.

### Recombinant expression and purification of sarbecovirus RBDs and S ectodomains and dimeric ACE2 ectodomains

Recombinant sarbecovirus RBDs and S ectodomains and dimeric ACE2 ectodomains were expressed as previously described^7,37,108^. Briefly, Expi293 cells were grown to a density of 3 x 10^6^ cells/mL and transfected using the Expifectamine293 transfection kit following the manufacturer’s recommendations. Three to five days after transfection, the supernatants were collected, clarified by centrifugation, and flowed over HisTrap FF (Cytiva) or HP (Cytiva) columns or incubated with Ni Sepharose excel resin (Cytiva).

The columns or resin were then washed with at least ten column volumes of wash buffer containing either 100 mM Tris pH 8.0, 300 mM NaCl, and 10-20 mM imidazole or 25 mM sodium phosphate pH 8.0, 300 mM NaCl, and 20 mM imidazole. Proteins were then eluted using a gradient up to 500 mM imidazole after which the the proteins were further purified by size-exclusion chromatography using a Superdex 200 Increase 10/300 GL or Superose 6 Increase 10/300 GL column into either 100 mM Tris pH 8.0 and 300 mM NaCl or 25 mM sodium phosphate pH 8.0 and 300 mM NaCl. A subset of the proteins were further biotinylated after purification using the BirA biotin-protein ligase reaction kit (Avidity) following the manufacturer’s recommendations. The biotinylated proteins were then re-purified using a HisTrap FF or HP column and size-exclusion chromatography. Both biotinylated and non-biotinylated proteins were concentrated, flash frozen, and stored at -80°C until use.

### Recombinant expression and purification of ACE2-Fc and monoclonal antibodies

Expi293 cells were transfected with hACE-Fc or paired heavy and light chain constructs supplied at a 1:1 ratio for LY-CoV555 using 4.5% transfection-grade polyethyleneimine (PEI) in Opti-MEM Reduced Serum Medium. The transfected cells were grown at 33°C for 3 days at 70% humidity, 8% CO2, and rotating at 150 RPM. The supernatant was collected, clarified by centrifugation after which Poly(diallyldimethylammonium chloride) was added to a final concentration of 0.0375%. The supernatant was again clarified by centrifugation and filtered using a 0.22 µM filter. The clarified supernatant was flowed over a MabSelect PrismA 2.6x5 cm column using an AKTA Avant150 FPLC system.

The column was washed with five column volumes of 20 mM Na_3_PO_4_, 150mM NaCl pH 7.2. Proteins were eluted using five column volumes of 100mM glycine pH 3.0, and neutralized with one-eighth volumes of 2 M Tris pH 8.0.

To produce mACE2-Fc, 80R, and SK26, Expi293 cells were grown to a density of 3 x 10^6^ cells/mL and transfected with the Expifectamine293 transfection kit following the manufacturer’s recommendations. A 1:1 ratio of heavy to light chain construct was used for producing 80R and SK26. The culture was cultivated for 4 to 5 days after which the supernatant was clarified by centrifugation and flowed over a 1-ml HiTrap Protein A HP affinity column (Cytiva). The column was washed with at least ten column volumes of 20 mM Na_2_HPO_4_, pH 8.0 or 50 mM Tris pH 7.4 150mM NaCl and the protein was eluted using 5 column volumes of 100 mM citric acid pH 3.0 or on a linear gradient from 50mM Tris pH 7.4 150 mM NaCl to 100 mM Citric Acid pH 3.0 100 mM NaCl, which was immediately neutralized with 1 M Tris-HCl pH 9.0. The protein was exchanged into 1x PBS, pH 7.4 (10 mM Na_2_HPO_4_, 1.8 mM KH_2_PO_4_, 2.7 mM KCl, 137 mM NaCl) or 50 mM Tris pH 7.4 150 mM NaCl concentrated, flash frozen, and stored at -80°C until use.

### I53-50 Nanoparticle Expression, Assembly, and Purification

To produce the individual RBD-I53-50A components, Expi293F cells were transfected using PEI MAX and cultured for six days at 37°C, 8% CO2, rotating at 150 RPM. The supernatant was clarified via centrifugation and filtered with a 0.22 µM filter. The clarified supernatant was collected and flowed over a Ni-Excel column. The column was then washed with 10 column volumes of 20 mM Tris pH 8.0, 300 mM NaCl or 25 mM sodium phosphate pH 8.0, 300 mM NaCl, 40 mM imidazole, after which the proteins were eluted in a two-stage period, first eluting with 1 column volume of elution buffer (20 mM Tris pH 8.0 or 25 mM sodium phosphate pH 8.0, 300 mM NaCl, 300mM imidazole) followed by 1.5 column volumes of elution buffer. The proteins were concentrated using Amicon Ultra-15 10,000 or 30,000 MWCO centrifugal concentrators and further purified via size-exclusion chromatography using a Superdex 200 Increase or Superose 6 increase 10/300 column on an AKTA Avant using a running buffer consisting of 50 mM Tris pH 7.4, 185 mM NaCl, 100 mM Arginine, 4.5% v/v Glycerol, 0.75% w/v CHAPS. The proteins were flash-frozen and stored at -80°C until use.

I53-50B.4PT1 was expressed in Lemo21(DE3) cells that were cultured in LB media (1% tryptone, 0.5% yeast extract, 1% NaCl) and aerated via a dissolved-oxygen cascade aerator. At an OD of ∼12, 100 mL of 100% glycerol and 1 mL of 1 M IPTG was added to the culture to induce expression. The culture was harvested via centrifugation and the resulting pellet was resuspended in PBS, homogenized, and lysed via a Microfluidics M100P Microfluidizer at 18,000 psi. The lysate was centrifuged and the resulting supernatant was discarded. The pellet was washed with 1x PBS, 0.1% Triton X-100, pH 8.0 before being resuspended in 1x PBS (137 mM NaCl, 2.7 mM KCl, 10 mM Sodium Phosphate, 1.8 mM Potassium Dihydrogen Phosphate), 2 M urea, 0.75% CHAPS pH

8.0 and applied to a DEAE Sepharose QFF Column on an AKTA Avant 150 FPLC system. The DEAE Sepharose column was washed with 1x PBS, 0.1% Triton X-100 and then washed five times more with 1x PBS, 0.75% CHAPS. The protein was then eluted with 1x PBS, 500 mM NaCl, concentrated with Amicon Ultra-15 10,000 MWCO centrifugal concentrators, filtered with a 0.22 µM syringe filter, and assessed for endotoxin levels prior to being used for nanoparticle assembly. The proteins were flash-frozen and stored at -80°C until use.

To assemble the monovalent Wu and BA.5 I53-50 RBD-NPs, the RBD-I53-50A and I53-50B components were mixed at a 1.1:1 molar ratio and incubated for 1 hour at 20°C. To assemble multivalent mosaic I53-50 RBD-NPs, three to four unique RBD-I53-50A components were combined at an equimolar ratio before this RBD-I53-50A mixture was mixed I53-50B at a 1.1:1 molar ratio and incubated for 1 hour at 20°C. To assemble the monovalent BtKY72 and SARS-CoV-1 RBD-NPs, the RBD-I53-50A and I53-50B components were mixed at a 1.1:1 molar ratio and incubated for 30 minutes at room temperature in 50 mM Tris pH 7.4, 100 mM Arginine, 4.5% glycerol, 0.75% CHAPS and were used immediately or eluted on a superose 6 increase 16/40 increase equilibrated in 50mM Tris pH 7.4, 100mM Arginine, 4.5% glycerol, 0.75% CHAPS. The assembled NPs were then concentrated using Amicon Ultra-15 10,000 MWCO centrifugal concentrators and purified via size-exclusion chromatography using a Superose 6 Increase 10/300 gel filtration column on an AKTA Avant. The assembled NPs were then flash-frozen and stored at -80°C until use.

### Dynamic Light Scattering

Dynamic light scattering (DLS) was performed on the RBD-NPs using an Uncle NanoDSF (Unchained laboratories). The RBD-NPs were loaded into an 8.8 µL quartz capillary cuvette and measured with ten 5 s acquisitions per sample at 20°C. Increased viscosity due to glycerol in the buffer was accounted for by the Uncle analysis software.

### Negative Stain Electron Microscopy

RBD-NPs diluted to 0.1 mg/mL in 50mM Tris pH 8.0, 150mM NaCl, 100mM Arginine, 4.5% v/v Glycerol, 0.75% w/v CHAPS and were added to freshly glow-discarded lacey carbon grids and stained with 2% uranyl formate. Data were acquired using a Talos L120C G2 Transmission Electron Microscope at a nominal magnification of 57,000x and -1.5 µm defocus.

### Bio-Layer interferometry (BLI)

To confirm the antigenicity of the assembled RBD-NPs, monoclonal antibodies or Fc-tagged ACE2 proteins diluted to 10 µg/mL in Octet Buffer (1x HBS-EP, 0.05% non-fat milk, 0.02% sodium azide) were first loaded onto ProteinA biosensors for 100 seconds. Next, the IgG- or ACE2-coated biosensors were equilibrated for 60s into Octet Buffer. The biosensors were dipped into the RBD-NPs diluted to 100 nM in Octet buffer for 200 seconds after which the biosensors were then dipped into Octet buffer for an additional 200 seconds. The resulting data were plotted in GraphPad Prism 10.

### Sandwich BLI

To biotinylate the SK26 IgG, the protein was exchanged into PBS (137 mM NaCl, 2.7 mM KCl, 10 mM Na_2_HPO_4_, 1.8 mM KH_2_PO_4_, pH 7.4) and concentrated to 2 mg/mL using a 100-kDa Amicon filter. The IgG was then biotinylated using the EZ-Link Sulfo-NHS-SS-Biotinylation Kit (ThermoFisher) using a 20-fold molar excess of Sulfo-NHS-SS-biotin and incubating the reaction at room temperature for 30 minutes. Following incubation, 1M Tris pH 8.0 was added to the IgG solution to a final concentration of 25 mM to quench the reaction. The biotinylated SK26 IgG was then purified into 50mM Tris pH 7.4 150mM NaCl by size-exclusion chromatography using a Superdex 200 Increase 10/300 GL column.

Multivalent incorporation of RBD components in mosaic nanoparticles was assessed with BLI using the Octet RED96 instrument (Sartorius) and the Octet Data acquisition software. BLI measurements were performed at 30°C and shaking at 1,000 rpm. After equilibrating streptavidin biosensors in 10x kinetics buffer (Sartorius) for 60s, biotinylated SK26 diluted to 2.5 µg/mL in 10x kinetics buffer was loaded onto biosensors to a 1 nm shift. The biosensors were then equilibrated in 10x kinetics buffer for 60 seconds before being dipped into tetravalent mosaic, trivalent mosaic, SARS-CoV-1, or BtKY72 RBD-NP diluted in 10x kinetics buffer to a final concentration of 10 µg/mL. The biosensors were equilibrated in 10x kinetics buffer for 60s prior to a second association with 80R diluted in 10x kinetics buffer to a final concentration of 10 µg/mL for 300s.

Dissociation was observed by dipping the biosensors in 10x kinetics buffer for 300s. Sensorgrams were plotted in GraphPad Prism10.

### Mouse immunizations and viral challenge

For the mosaic RBD-NP alone study, nanoparticles were diluted in sterile formulation buffer and mixed with 1:1 v/v AddaVax to obtain a final dose of 1 μg of antigen per animal, per injection. At 8 weeks of age, 20 mice per group were injected subcutaneously in the inguinal region with 100 μL total volume at weeks 0 and 3.

Animals were bled using the submental route at week 5. Mice were then shipped to UNC for challenge.

For the pre-immune study, mRNA-1273 was diluted in sterile PBS according to manufacturer’s instructions to obtain a final dose of 0.3 μg of mRNA-1273 per animal, per injection. Nanoparticles were diluted in sterile formulation buffer and mixed with 1:1 v/v AddaVax to obtain a final dose of 1 μg of antigen per animal, per injection. At 8 weeks of age, 10 mice per group were anesthetized via isoflurane and injected intramuscularly in the quadriceps unilaterally with 50 μL of mRNA-1273 at -6 and -3 weeks followed by bilateral intramuscular injections of the nanoparticles in the quadriceps with 50 μL per leg at 0 and 3 weeks. Animals were bled using the submental route at weeks 2 and 5. Mice were then shipped to UNC for challenge. Mice were provided with 0.05 mg/kg buprenorphine HCl preemptively prior to all IM injections to reduce pain. For all bleeds, whole blood was collected in serum separator tubes (BD #365967) and rested for 30 min at room temperature for coagulation. Tubes were then centrifuged for 10 min at 2,000 x *g* and serum was collected and stored at -80°C until use.

After receipt, mice were moved into the Animal Biosafety Level 3 (ABSL3) Laboratory at the University of North Carolina at Chapel Hill and acclimated for 7 days. For infection, mice were anesthetized with a mixture of Ketamine/Xylazine and inoculated intranasally with either 10^5^ pfu of RsSHC014 MA15^109^ or 10^5^ pfu SARS-CoV-2 MA10 XBB.1.5^110^, respectively. Following infection, all mice were monitored for weight loss and on day 2 and 4 post infection groups of mice were euthanized, and the lung congestion score was assessed (score 0 to 4, with 0=no discoloration, 4=discoloration of the entire lung tissue). On each harvest day, the caudal lung lobe and nasal turbinates were collected for determination of the viral load by plaque assay.

For plaque assays, collected tissue samples were homogenized in 1ml PBS, clarified by centrifugation, serially diluted, and inoculated onto confluent monolayers of Vero E6 cells (ATCC CCL-81), followed by an overlay with agarose. Plaques were visualized at 2 (RsSHC014 MA15) or at 3 (SARS-CoV-2 MA10 XBB.1.5) days post infection by staining with neutral red dye.

Statistical differences in weight loss were determined using a mixed-effects analysis with Dunnett’s post-test using GraphPad Prism 10. All other statistical differences were assessed with the Kruskal-Wallis test with Dunn’s post-test using GraphPad Prism 10.

### African Green monkey immunizations and viral challenge

AGMs were distributed into treatment groups on the basis of sex and weight. Animals were housed and immunized under ABSL-2 containment. The *in vivo* challenge and post-challenge phases of the study were performed under ABSL-3 containment. mRNA-1273 was diluted in sterile PBS according to the manufacturer’s instructions to obtain a final dose of 30 µg per animal. Nanoparticles were diluted in sterile formulation buffer and mixed with 1:1 v/v AddaVax to obtain a final dose of 40 µg per animal. Animals were sedated and immunized intramuscularly in the quadriceps with mRNA-1273 at 0 and 1 month and with nanoparticles at 8 or 10 and 11 or 13 months. Two weeks after each nanoparticle dose, the animals were anesthetized by IM injection of 10 mg/kg ketamine HCl and blood was collected into EDTA tubes via femoral venipuncture. Blood samples were processed to isolate plasma and PBMCs.

Viral stocks of SARS-CoV-2 XBB.1.5 were obtained from BEI Resources (NR-59105) and stocks of RsSHC014 were obtained from Ralph Baric’s laboratory at the University of North Carolina at Chapel Hill. The stocks were diluted in PBS to a concentration of 5 x 10^5^ PFU/mL for SARS-CoV-2 XBB.1.5 and 5 x 10^6^ PFU/mL for RsSHC014. Animals were sedated and administered the challenge virus via IN and IT routes. 0.5 mL of the challenge virus was administered dropwise to each nostril, for a total of 1 mL via the IN route. A second 1 mL of the challenge virus was administered via the IT route. A total dose of 10^6^ PFU for SARS-CoV-2 XBB.1.5 and 10^7^ PFU for RsSHC014 was administered per animal. At 1, 2, and 4 dpi, the animals were sedated and nasal swabs, oral swabs, and bronchoalveolar lavage (BAL) were collected from each animal. At 4 dpi, animals were euthanized via sodium pentobarbital overdose and lung, nare, trachea, and tonsil tissues were collected and stored in 10% neutral buffered formalin at 2-8°C for histological analyses.

For TCID50 assays, BAL, nasal swabs, and oral swabs were centrifuged at 450 x g for 10 minutes at 4°C after collection and frozen at ≤ -70°C until use. Vero-TMPRSS2 cells were split into clear, flat-bottom 96-well plates and incubated at 37°C and 5% CO_2_ until a confluent monolayer formed. The existing media was removed from the cells and 180 µL of fresh media was added to the cells. Next, 20 µL of sample was added to the top row and mixed by pipetting. The sample was serially diluted 10-fold down the plate, such that the serial dilutions ranged from 1:10 to 1:10^8^. The plates were incubated for 4 days at 37°C and 5% CO_2_ after which the cell monolayers were visually inspected for cytopathic effects and scored. The TCID_50_ was calculated using the Reed-Muench formula. TCID_50_ measurements were performed in quadruplicates for each sample.

Statistical differences were assessed with the Kruskal-Wallis test with Dunn’s post-test using GraphPad Prism 10.

### Histology

Mouse infections: For each mouse the left lung was incubated in formalin for at least 7 days to fix tissue and inactivate virus. The fixed tissue was processed and embedded in paraffin. 5 μm sections were cut and stained with either Congo red or hematoxylin and eosin (H&E). Eosinophils were enumerated by counting the Congo red positive leukocytes in 10 high power fields (400X final magnification) per mouse lung. Airway pathology was assessed in H&E stained sections to assess interstitial pneumonitis (score = percentage of pulmonary alveolar parenchyma with septae expanded by leukocyte; scored for 10 100X fields).

AGM Infections: For each AGM, nares, trachea and bilateral lungs were incubated in formalin for at least 7 days to fix tissue and inactivate virus. The nares were decalcified with EDTA prior to processing. The fixed lungs were examined grossly to determine the fraction of each lobe that was consolidated (i.e. firm due to filling of the airspaces), and representative sections from each lobe were sampled for histology to confirm gross observations. Sampled tissues were processed and embedded in paraffin. 5 µm sections were stained with H&E and assessed for tracheal inflammation (scored as the percentage of the mucosa involved by leukocytic infiltration of the submucosa and epithelium).

### Enzyme-linked immunosorbent assay (ELISA)

For ELISAs using the recombinantly expressed, biotinylated sarbecovirus RBDs, the RBDs were diluted to 0.003 mg/mL in PBS and added to NeutrAvidin coated 384-well plates (ThermoFisher) and incubated overnight at room temperature. The next day, the plates were slapped dry and blocked with Blocker Casein for 1 hour at 37°C. The plates were slapped dry again and plasma samples diluted to a starting dilution of 1:40 in TBS with 0.1% Tween 20 (TBST) and serially diluted 1:4 in TBST thereafter were added to the plates. The plates were incubated for 1 hour at 37°C, slapped dry, and washed four times with TBST. Next, a goat anti-human IgG (H+L) HRP conjugated antibody (ThermoFisher) diluted 1:5,000 in TBST was added to each well and the plates were incubated for 1 hour at 37°C. The plates were again slapped dry and washed four times with TBST. SureBlue Reserve TMB 1-Component Microwell Peroxidase Substrate (SeraCare) was added to each well and the reaction was developed for 90 seconds after which an equal volume of 1N HCl was added to stop the reaction. The absorbance at 450 nm was measured immediately after stopping the reaction using a BioTek Synergy Neo2 plate reader. The resulting data were analyzed using GraphPad Prism 10 using a four parameter logistic curve to determine the ED_50_ for each sample. Two biological replicates, each with two technical replicates, were performed using two distinct batches of protein.

The ELISAs against the recombinantly expressed sarbecovirus S ectodomains were performed similarly to ELISAs against the sarbecovirus RBDs with one modification. The diluted S proteins were added to 384-well Maxisorp plates (ThermoFisher) and incubated at room temperature overnight.

### Pseudotyped virus production

Vesicular stomatitis virus (VSV) pseudotyped with the full-length sarbecovirus S proteins were produced as previously described^7,37,108,111^. In brief, HEK-293T cells were split into 10 cm poly-lysine coated plates and grown overnight at 37°C and 5% CO_2_ until they reached approximately 90% confluency. The cells were then washed three times with Opti-MEM, transfected with the sarbecovirus S constructs using Lipofectamine 2000 and incubated for 5 hours at 37°C and 5% CO_2_ after which an equal volume of DMEM containing 20% FBS and 2% PenStrep was added to the cells. Twenty to 24 hours after transfection, the cells were washed three times with DMEM, infected with VSVΔG/luc, and incubated for 2 hours at 37°C and 5% CO_2_. Next, the cells were washed five times with DMEM supplemented with an anti-VSV-G antibody (I1-mouse hybridoma supernatant diluted 1:25 for CRL-2700, ATCC) and incubated for 20 to 24 hours at 37°C and 5% CO_2_. Following this incubation, supernatant was harvested, clarified by centrifugation, filtered using an 0.45 µM filter, and concentrated with a 100 kDa centrifugal filter (Amicon). The resulting pseudovirus was frozen at -80°C until use.

### Pseudovirus neutralization assay

Neutralization assays were performed as previously described^7,37,108,111^ using Vero-TMPRSS2 cells as the target cell line for SARS-CoV-2 variants and SARS-CoV-1 against mice sera, HEK-ACE2 cells as the target cell line for SARS-CoV-1 against African green monkey plasma and RsSHC014, and HEK-293T cells transiently transfected with *Rhinolophus alcyone* ACE2 for the clade 3 sarbecoviruses. For Vero-TMPRSS2 cells, the cells were split into white-walled, clear bottom 96-well plates and grown overnight at 37°C and 5% CO_2_ until they reached 80-90% confluency. HEK-ACE2 cells were split into poly-lysine coated, white-walled, clear bottom 96-well plates and incubated overnight at 37°C and 5% CO_2_ until they reached 80-90% confluency. To transfect HEK-293T cells with *R. alcyone* ACE2, HEK-293T cells were split into 10 cm, poly-lysine coated plates and grown overnight at 37°C and 5% CO_2_ until they reached approximately 90% confluency. The cells were then transfected with a construct encoding *R. alcyone* ACE2 using Lipofectamine 2000 and incubated for 5 hours at 37°C and 5% CO_2_ after which the cells were trypsinized and plated into poly-lysine coated, white-walled, clear bottom 96-well plates. The cells were again grown overnight at 37°C and 5% CO_2_ until they reached 80-90% confluency. Plasma samples were diluted in DMEM at a starting concentration between 1:10 and 1:33 and serially diluted 1:3 thereafter. An equal volume of pseudotyped VSV diluted 1:25 in DMEM was added to diluted plasma and the virus-plasma mixture was incubated at room temperature for 30 minutes. The growth media was removed from the target cell line and replaced with the virus-plasma mixture and the cells were incubated for 2 hours at 37°C and 5% CO_2_ after which an equal volume of DMEM supplemented with 20% FBS and 2% PenStrep was added to each well. The cells were incubated for an additional 16 to 20 hours at 37°C and 5% CO_2_. Next, ONE-Glo EX (Promega) was added directly to each well. The plates were then incubated at 37°C for 5 minutes with constant shaking and the luminescence values were recorded using a BioTek Synergy Neo2 plate reader.

Data were normalized using GraphPad Prism 10 with the relative light unit (RLU) values obtained from uninfected cells to define 0% infectivity and the RLU values obtained from cells infected with pseudovirus only to define 100% infectivity. ID_50_ values for plasma samples were determined using the normalized data using an [inhibitor] vs. normalized response - variable slope model. At least two biological replicates using distinct batches of pseudoviruses were performed for each sample.

### Plasma antibody depletion assays

PureCube Ni-INDIGO MagBeads (Cube Biotech) were collected using a magnet and washed once with TBS (20 mM Tris, 100 mM NaCl, pH 7.4). An excess of His-tagged SARS-CoV-2 Wu RBD was added to the beads and the Wu RBD-bead mixture was incubated at room temperature for 30 minutes with constant rotation. The beads were collected again with a magnet, washed three times with TBS containing 0.01% Tween 20, and resuspended in TBS containing 0.01% Tween 20. Additional uncoupled beads were treated similar to above without the addition of a His-tagged protein and used as a mock depletion control.

Plasma samples were mixed with the Wu RBD-beads or uncoupled beads at a 1:4 volume ratio and incubated for 1 hour at 37°C. The beads were then collected with a magnet and the supernatant was transferred to a new tube containing an additional 4 volumes of beads. The plasma-bead mixture was incubated for an additional 1 hour at 37°C after which the beads were collected again with a magnet and the supernatant was transferred to a new tube. ELISAs against the SARS-CoV-2 Wu, SARS-CoV-2 BA.5, SARS-CoV-1, and BtKY72 RBDs were performed and analyzed as described above using either the Wu RBD-depleted or mock depleted plasma as input.

Neutralization assays against VSV pseudotyped with SARS-CoV-2 Wu-G614, SARS-CoV-2 BA.5, SARS-CoV-1, or BtKY72 S were performed and analyzed as described above using the Wu RBD-depleted or mock depleted plasma as input. Both the ELISAs and neutralization assays were performed in technical duplicate. Two biological replicates of the depletion assay were performed using distinct batches of Wu RBD for the depletion as well as distinct batches of biotinylated RBDs and pseudoviruses for the ELISAs and neutralization assays, respectively.

### DMS and pan-sarbeco RBD binding

Plasma from the three AGMs per group that exhibited the highest neutralizing antibody titers against XBB.1.5 were included in the pan-sarbecovirus RBD binding and deep mutational scanning analysis. Yeast-display RBD deep mutational scanning libraries for SARS-CoV-2 Wuhan-Hu-1^112^, SARS-CoV-2 XBB.1.5^113^, PRD-0038, and SARS-CoV-1 Urbani^14^ and a pan-sarbecovirus library^15,89^ were previously described. Binding of serum samples was evaluated against the pooled libraries via FACS coupled to high-throughput sequencing. Serum was first depleted of non-specific yeast-reactive antibodies as previously described^114^. Pooled yeast-display libraries were induced for yeast surface expression, and labeled with serum at 1:100 and 1:10,000 dilutions for one hour at room temperature. Yeast were washed with PBS-BSA and labeled with 1:100 FITC-conjugated chicken anti-Myc (Immunology Consultants CMYC-45F) to label for RBD surface expression and 1:200 PE-conjugated goat anti-human-IgG (Jackson ImmunoResearch 109-115-098) to label for bound serum antibody. Libraries were then partitioned into four bins of serum binding on a BD FACSAria, collecting approximately 5 million RBD^+^ cells per sample concentration across the four bins. Cells were grown up post-sort, plasmid purified, N16 barcode amplified via PCR, and sequenced on an Illumina NextSeq. Raw Illumina sequencing data are uploaded to the NCBI Sequence Read Archive, BioProject PRJNA714677, Biosample SAMN46431257.

Our complete pipeline for processing sequencing reads is available from GitHub: https://github.com/tstarrlab/SARSr-CoV_MAP_NHP-imprinting/blob/main/results/summary/summary.md. Barcode reads were mapped to library barcodes, and for each library variant an area under the curve (AUC) metric was derived from its distribution of sequence reads across sort bins. First, the strength of serum binding to each barcode at each serum dilution was determined as the simple mean bin from cell counts across integer-weighted FACS bins, and subtracted by per-barcode background based on a sorted sample from yeast libraries incubated with no sera. AUC of mean bin versus serum dilution was calculated. Any barcode with less than 2 cell counts at either serum labeling concentration was eliminated. We then computed per-mutant AUC as the mean of per-barcode AUC estimates for replicate barcodes linked to the identical RBD variant. Because mutations that disrupt RBD epxression will artefactually decrease serum binding, we also censored the AUC measurement for any mutant with a previously measured impact on RBD expression of >1 log-MFI unit (RBD expression < -1), and we normalized remaining AUC values by the slope of the linear model relating serum AUC and expression globally across all library variants with expression > -1. Per-barcode AUC values are available from: https://github.com/tstarrlab/SARSr-CoV_MAP_NHP-imprinting/blob/main/results/bc_sera_binding/bc_sera_binding.csv. Final per-variant AUC averages are available from https://github.com/tstarrlab/SARSr-CoV_MAP_NHP-imprinting/blob/main/results/final_variant_scores/final_variant_scores_dms.csv and https://github.com/tstarrlab/SARSr-CoV_MAP_NHP-imprinting/blob/main/results/final_variant_scores/final_variant_scores_wts.csv for DMS and pan-sarbecovirus pools, respectively.

### Competition ELISAs

Competition ELISAs were performed as previously described^111^. In brief, EC_50_ values were determined by plating dimeric human ACE2 or dimeric *Rhinolophus alcyone* ACE2 diluted to 0.003 mg/mL onto 384-well Maxisorp plates. The plates were incubated overnight at room temperature after which the plates were slapped dry and blocked with Blocker Casein at 37°C for 1 hour. The plates were slapped dry and biotinylated SARS-CoV-2 Wu-G614, SARS-CoV-2 BA.5, SARS-CoV-1, or BtKY72 S diluted to 0.3 mg/mL and serially diluted 1:3 in TBST thereafter was added to the plates. The plates were incubated for 1 hour at 37°C, slapped dry, and washed four times with TBST. Ultra streptavidin-HRP diluted 1:2,000 in TBST was then added to each well and the plates were incubated for an additional 1 hour at 37°C, slapped dry, and washed four times with TBST. SureBlue Reserve TMB 1-Component Microwell Peroxidase Substrate was added to each well and developed for 120s. An equal volume of 1 N HCl was added to quench the reaction. The absorbance at 450 nm was measured using a BioTek Synergy *Neo2* plate reader and the resulting data were analyzed with GraphPad Prism 10 using a four parameter logistic curve to determine the EC_50_ value for each S-ACE2 pair.

384-well Maxisorp plates were coated with dimeric human ACE2 or dimeric *Rhinolophus alcyone* ACE2 and blocked with Blocker Casein as described above. Plasma samples were diluted in 384-well non-binding plates to 1:5 starting dilution followed by a 1:3 serial dilution in TBST thereafter. Next, the biotinylated SARS-CoV-2 Wu-G614, SARS-CoV-2 BA.5, SARS-CoV-1, or BtKY72 S diluted in TBST to a concentration twice the determined EC_50_ was added to each well of the non-binding plate. The plasma-S mixtures were incubated for 1 hour at 37°C. Following this incubation, the plasma-S mixture was transferred to the ACE2-coated plates and the plates were incubated for an additional 1 hour at 37°C, slapped dry, and washed four times with TBST. Ultra streptavidin-HRP diluted 1:2,000 in TBST was then added to each well and the plates were incubated for 1 hour at 37°C, slapped dry, and washed four times with TBST. SureBlue Reserve TMB 1-Component Microwell Peroxidase Substrate was added to each well and developed for 120s after which an equal volume of 1 N HCl was added to each well. The absorbance at 450 nm was immediately measured using a BioTek Synergy *Neo2* plate reader. The resulting data were first normalized using GraphPad Prism 10 with the absorbance values from wells without S defining 0% binding and the absorbance values from well with S only defining 100% binding. BD_50_ values were determined from the normalized data using the [inhibitor] vs. normalized response – variable slope model. At least two biological replicates using distinct batches of S and ACE2 proteins were performed for each sample. Two technical replicates were performed for each biological replicate.

### Flow cytometry analysis of RBD-reactive memory B cells

RBD-streptavidin tetramers were generated by incubating biotinylated SARS-CoV-2 Wu, SARS-CoV-2 BA.5, SARS-CoV-1, or BtKY72 RBDs with fluorophore-conjugated streptavidin at a 4:1 molar ratio for 30 minutes at 4°C. After this initial incubation, an excess of free biotin was added to bind any unconjugated sites on the streptavidin and the mixture was incubated for an additional 30 minutes at 4°C. The resulting RBD-streptavidin were washed once with PBS (137 mM NaCl, 2.7 mM KCl, 10 mM Na_2_HPO_4_, 1.8 mM KH_2_PO_4_, pH 7.4) and concentrated with a 100-kDa cut-off centrifugal filter (Amicon). An additional streptavidin tetramer conjugated to the MERS-CoV S was generated as above and used as a decoy during staining. The following RBD-streptavidin tetramers were generated and used for staining: SARS-CoV-2 Wu RBD-streptavidin-BV421, SARS-CoV-2 BA.5 RBD-streptavidin-PE, SARS-CoV-2 BA.5 RBD-streptavidin-Alexa 647, SARS-CoV-1 RBD-streptavidin-PE, SARS-CoV-1 RBD-streptavidin-Alexa 647, BtKY72 RBD-streptavidin-PE, BtKY72 RBD-streptavidin-Alexa 647, and MERS-CoV S-streptavidin-BV480.

Approximately 5 to 10 million peripheral blood mononuclear cells (PBMCs) per animal per time point were thawed, collected by centrifugation, and washed once with PBS. The cells were stained with Zombie UV dye (Biolegend; diluted 1:100 in PBS) for 30 minutes at room temperature. Following staining, the cells were washed twice with flow staining buffer (0.1% BSA, 0.1% NaN_3_ in PBS). The cells were next stained for 30 minutes at 4°C with the RBD-streptavidin tetramers and the following antibodies diluted in Brilliant Stain Buffer (BD) with all antibodies used at the manufacturer’s recommended volume per test: CD3-BV650 (BD), CD20-BUV737 (BD), IgD-FITC (SouthernBiotech), IgG-BUV615 (BD). After incubation, the cells were washed three times with flow staining buffer, resuspended in 500 µL of flow staining buffer, and passed through a 35-µm filter. The cells were then examined on a BD FACSymphony A3 and FACSDiva software (BD) and FlowJo 10.10.0 for analysis. Gates for identifying the BA.5-SARS-CoV-1-BtKY72 RBD-double-positive and Wu RBD-positive and - negative populations were drawn based on staining of PBMCs collected from naive African Green monkeys.

## Supplemental Figures

**Figure S1.**
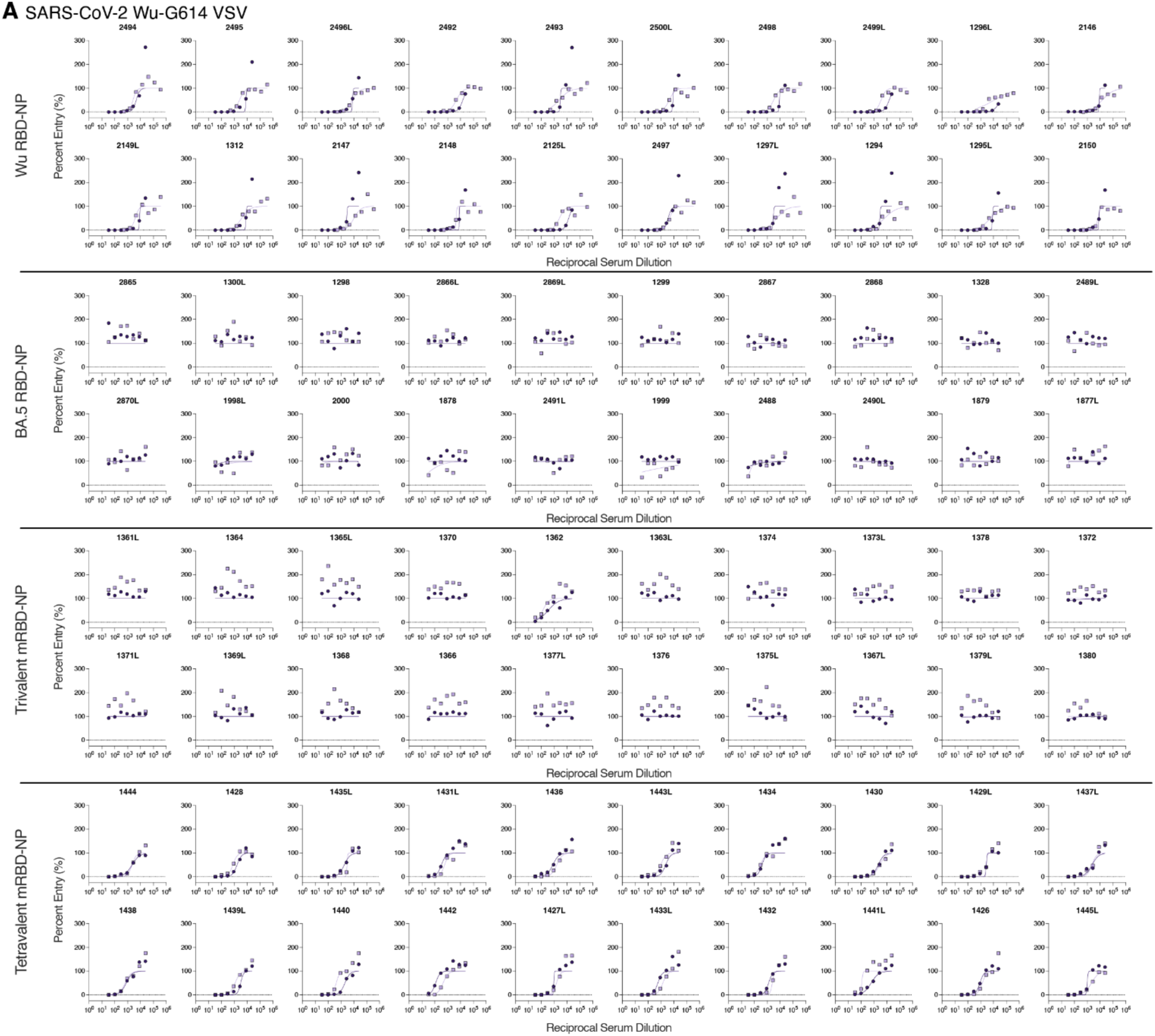
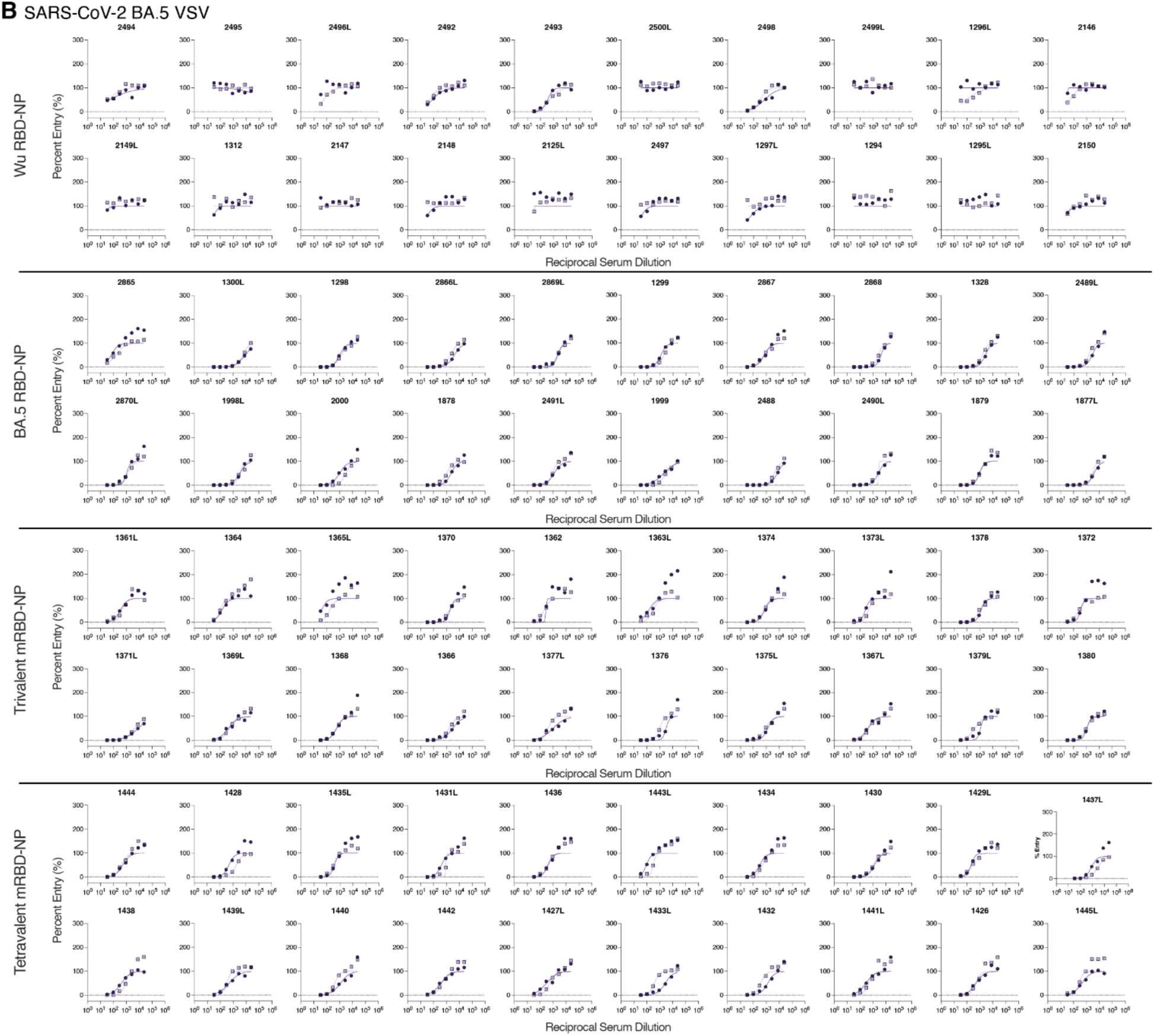
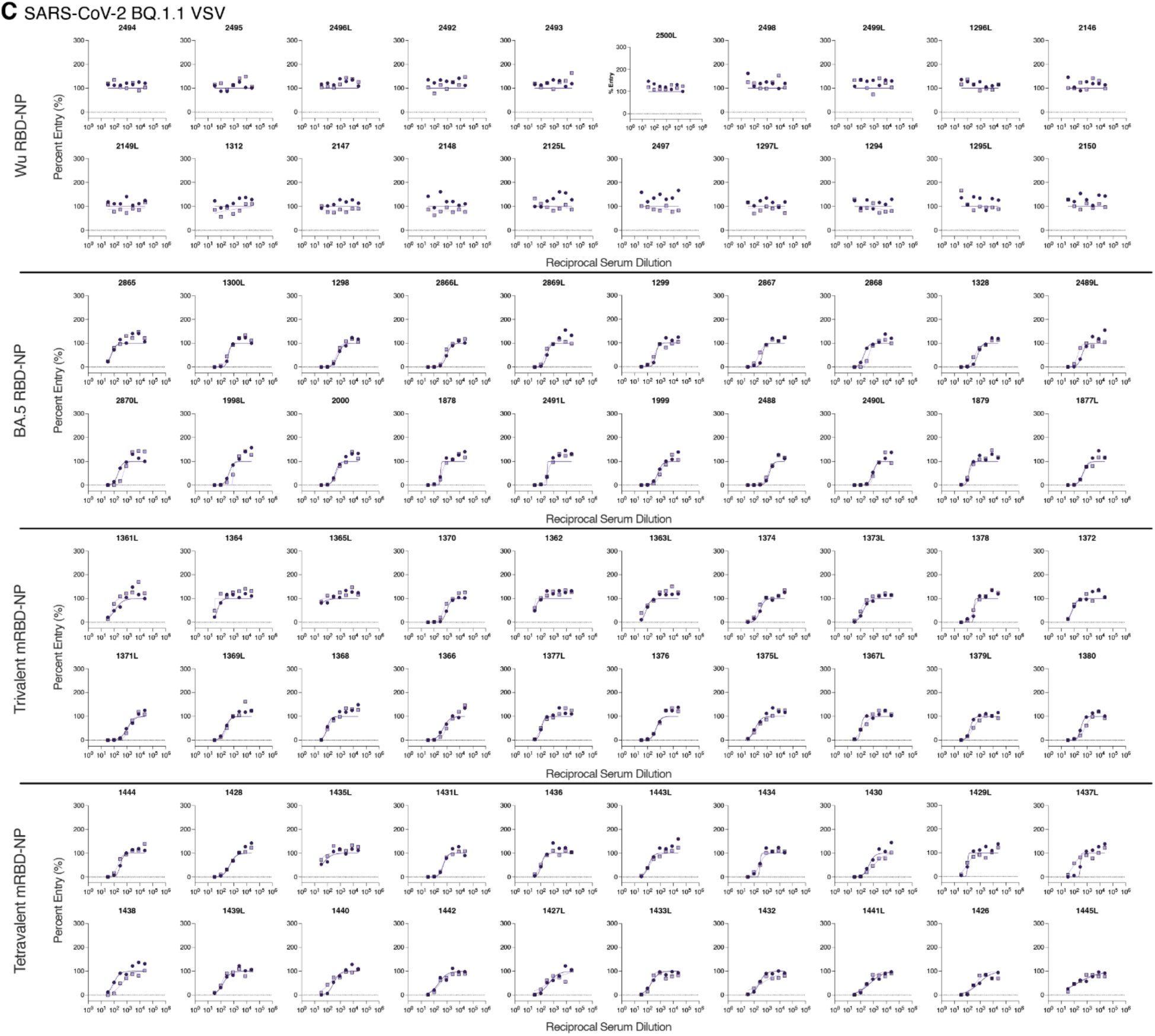
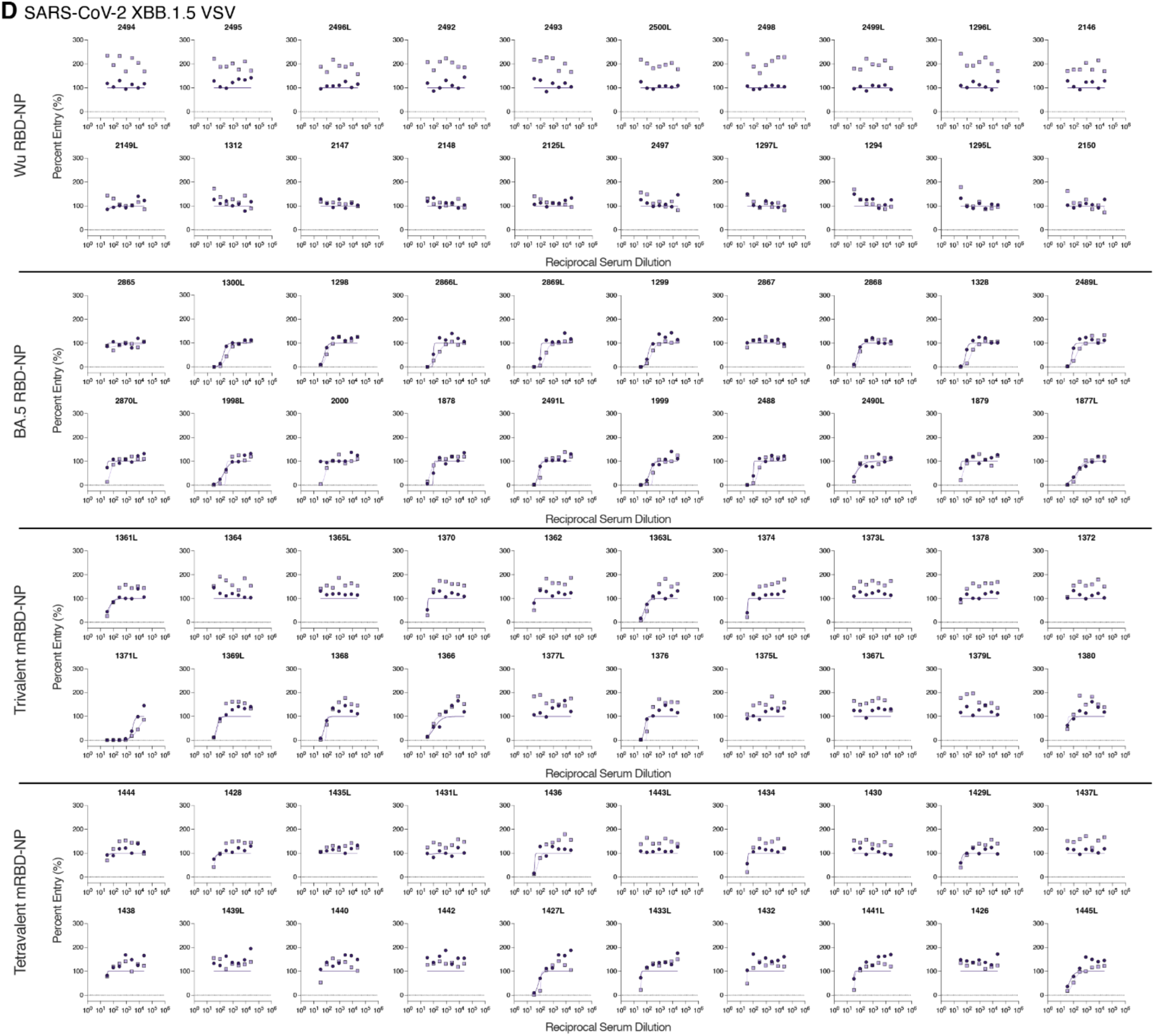
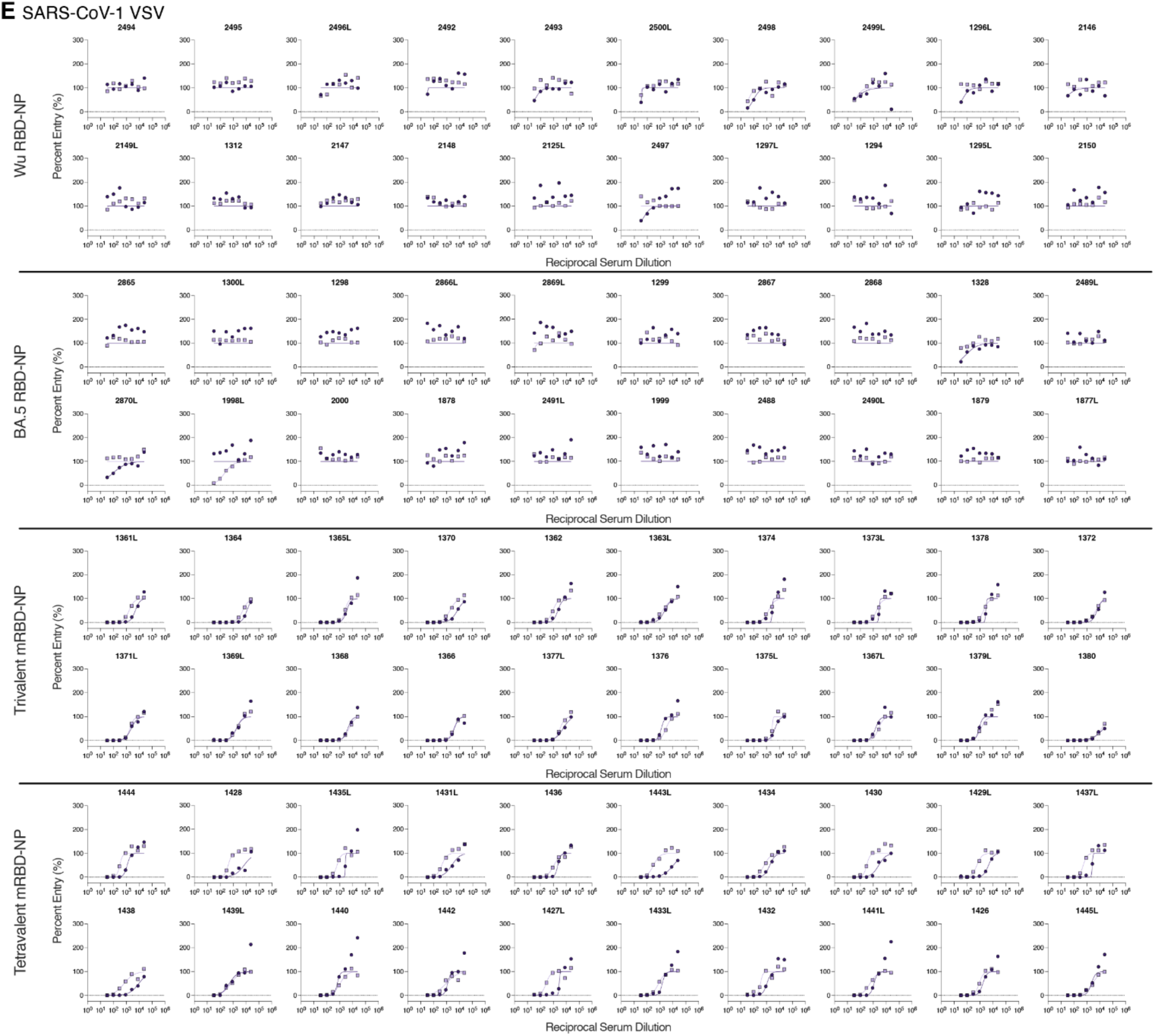
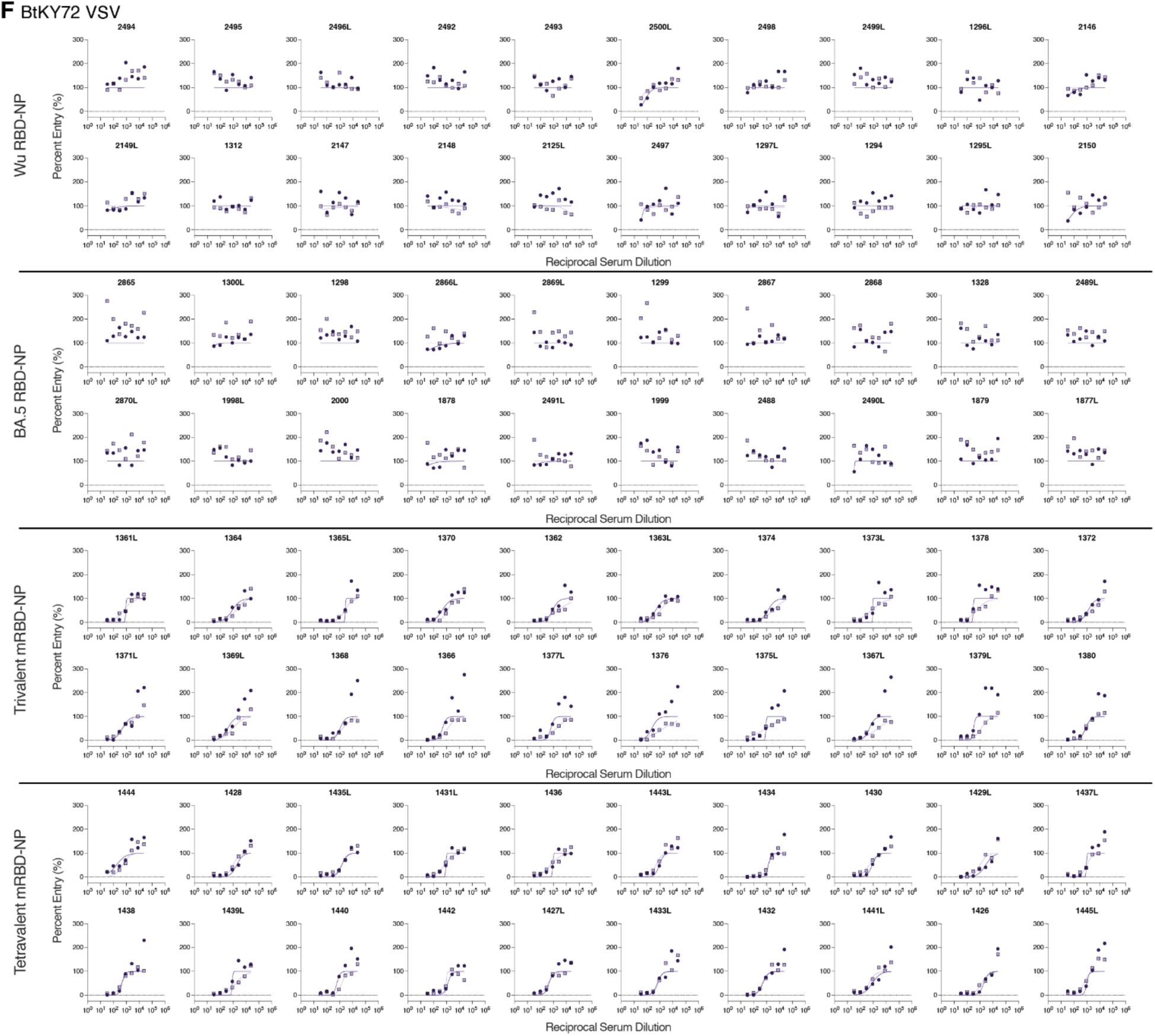
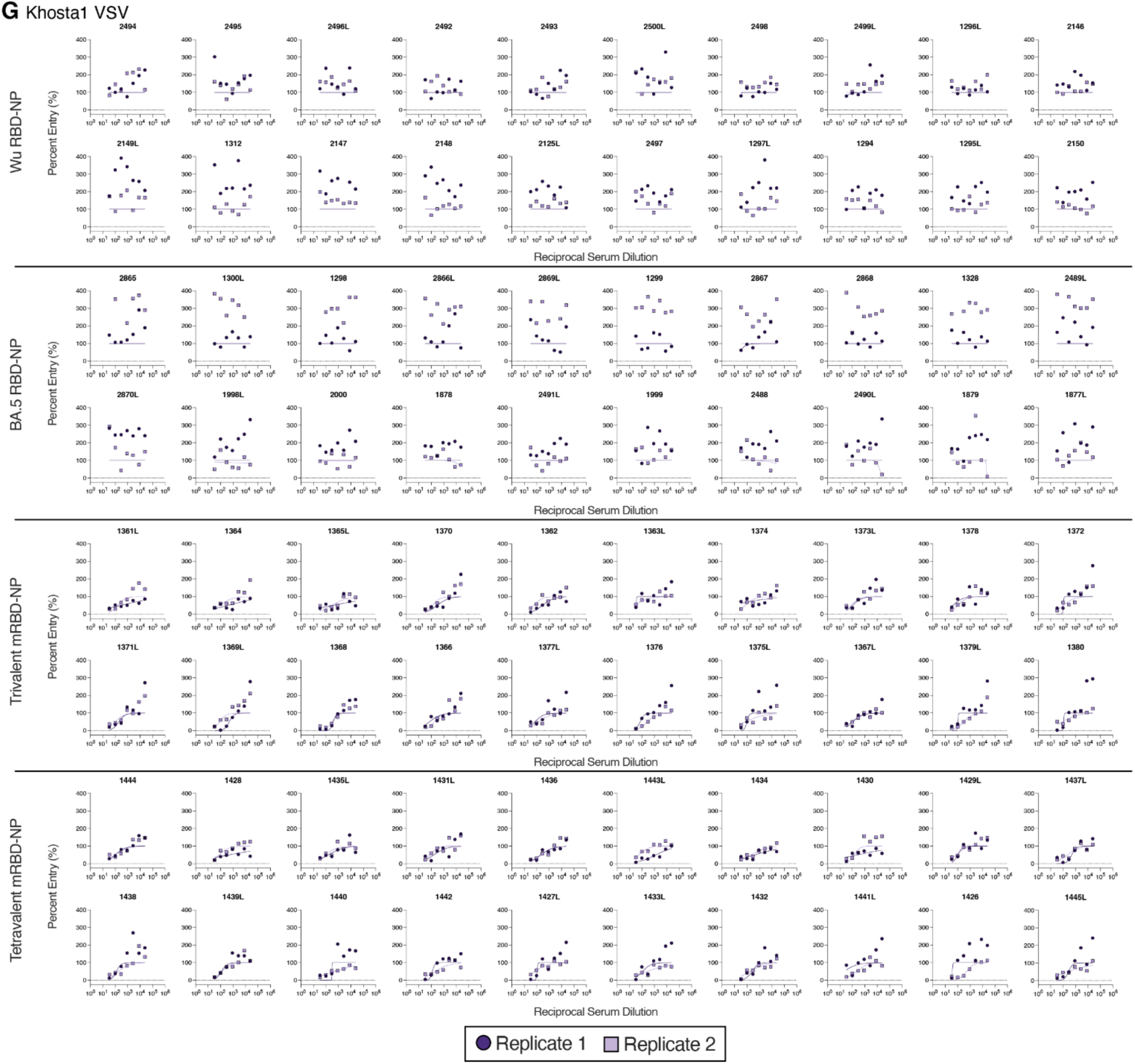
Neutralization dose-response curves for RBD-NP-immunized naive mice. Serum neutralizing antibody titers were assessed using VSV pseudotyped with the **A)** SARS-CoV-2 Wu-G614, **B)** SARS-CoV-2 BA.5, **C)** SARS-CoV-2 BQ.1.1, **D)** SARS-CoV-2 XBB.1.5, **E)** SARS-CoV-1, **F)** BtKY72, or **G)** Khosta1 S. Two biological replicates were conducted using distinct batches of pseudovirus.

**Figure S2.**
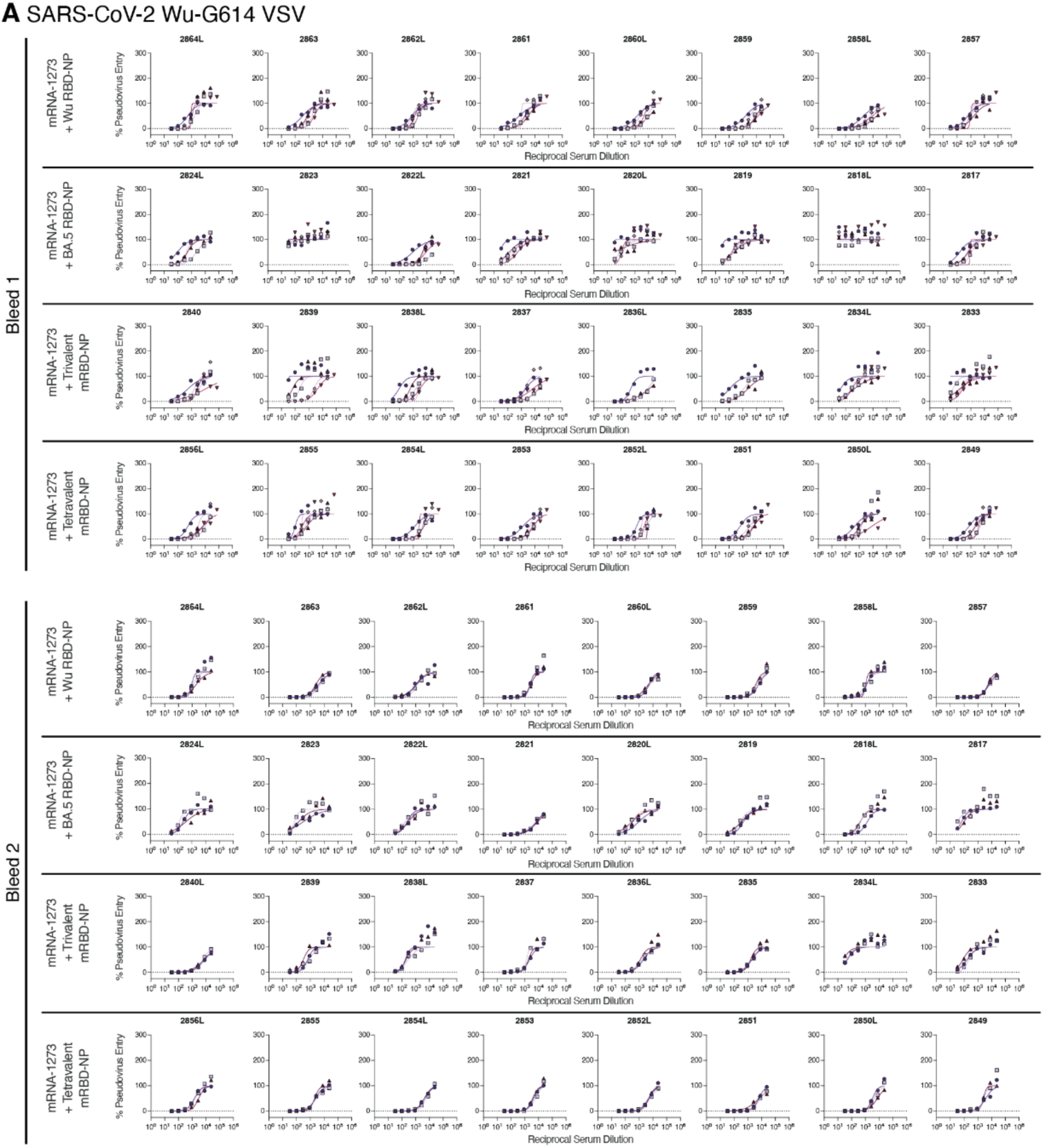
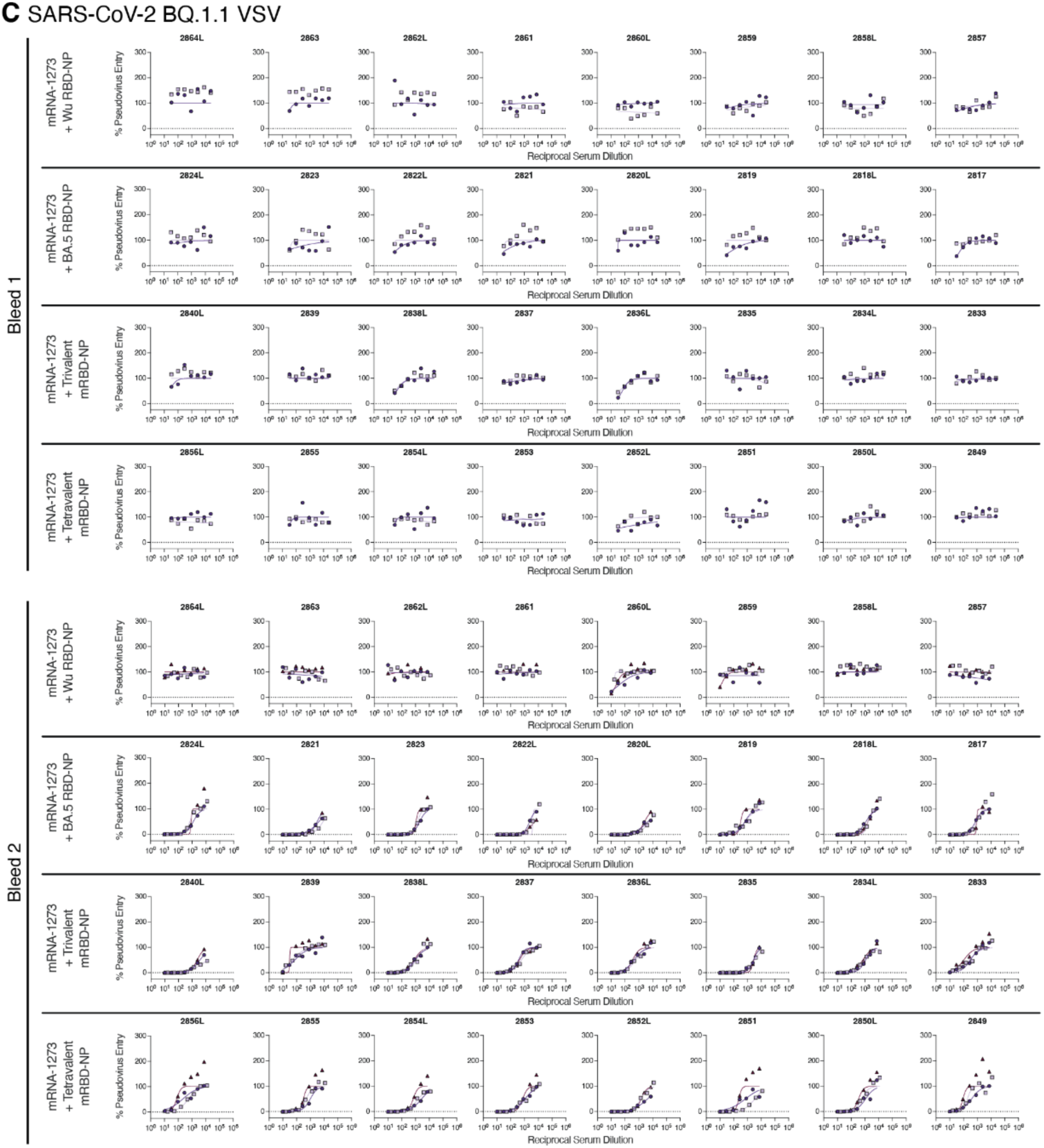
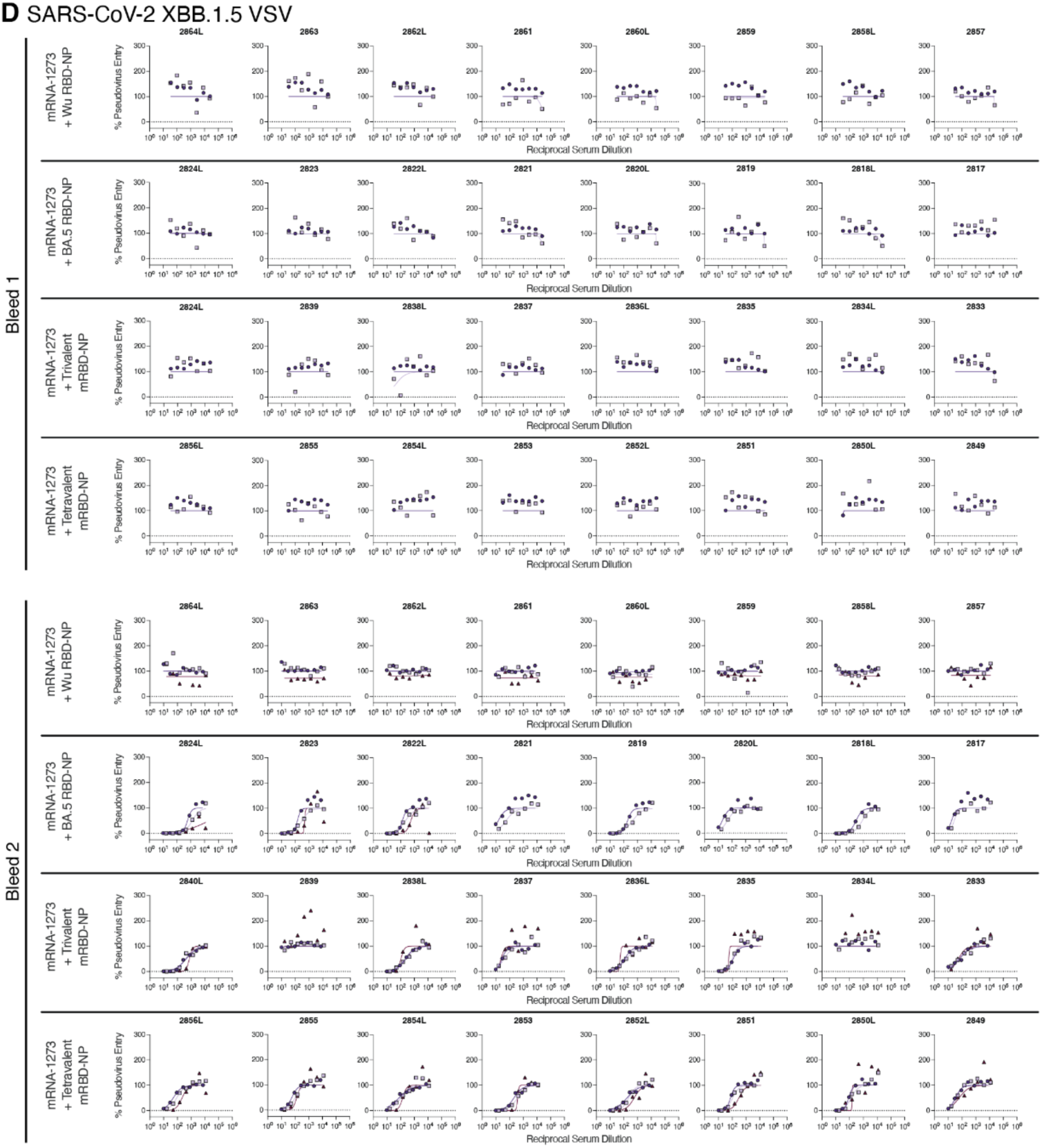
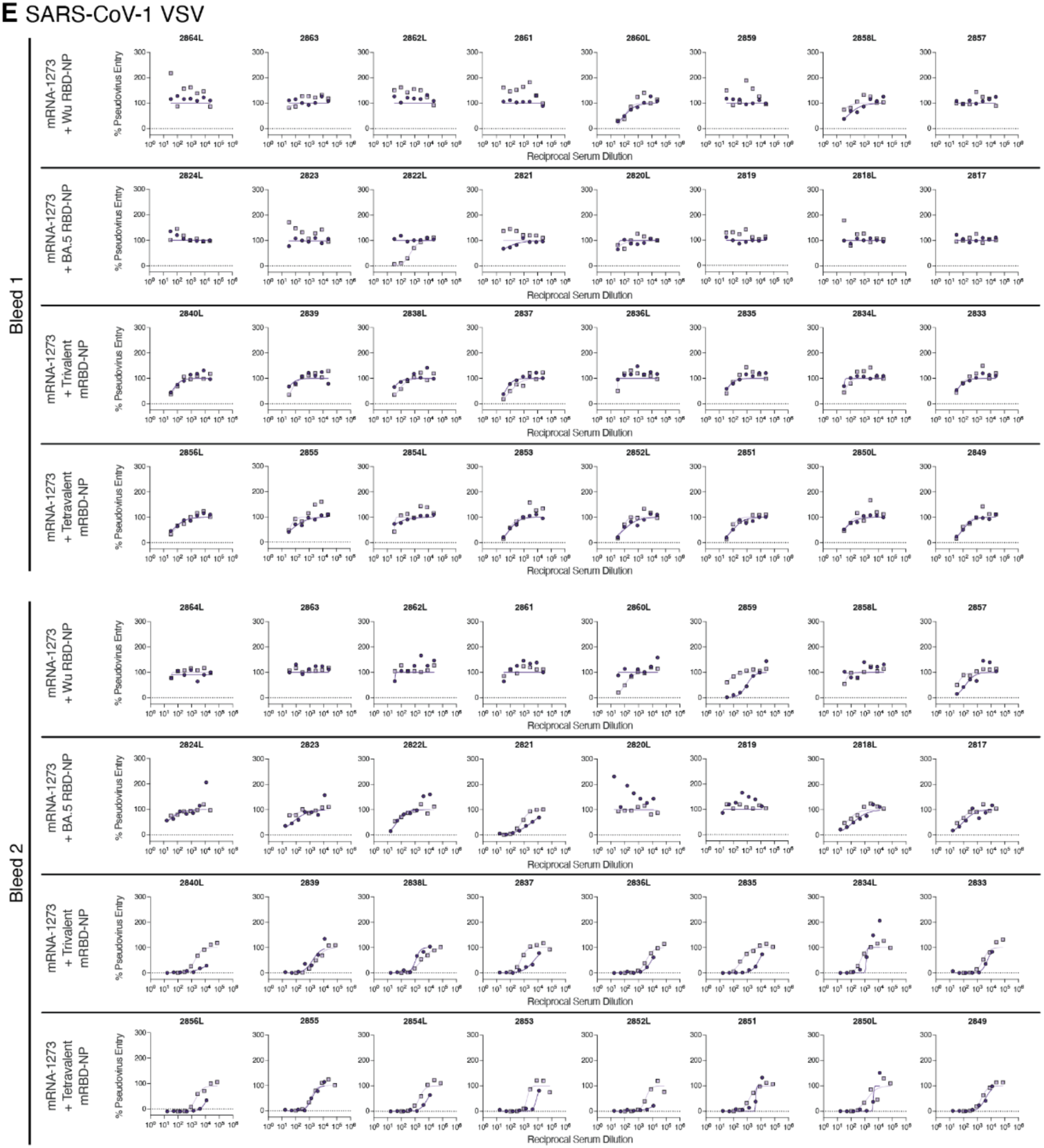
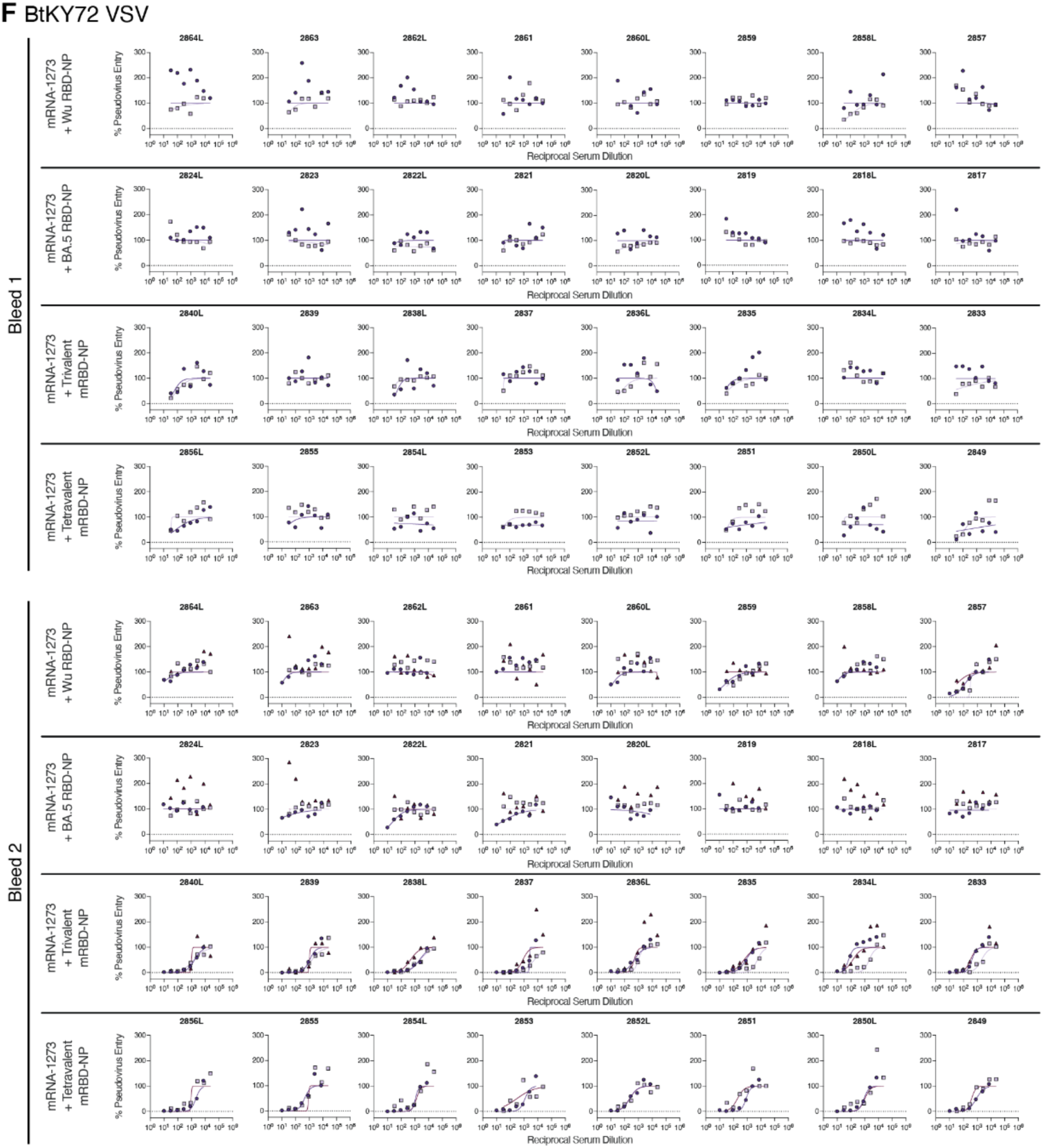
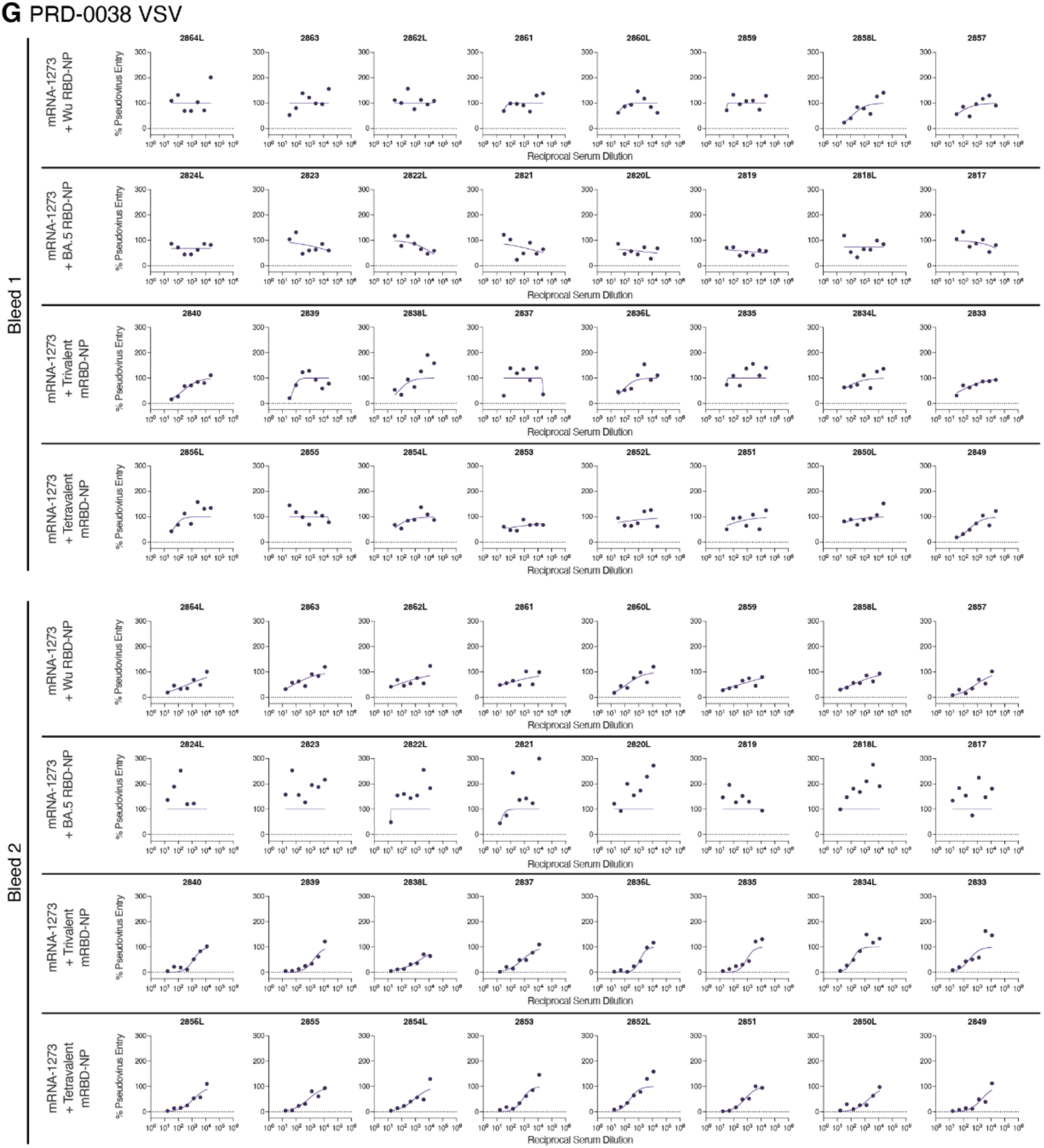
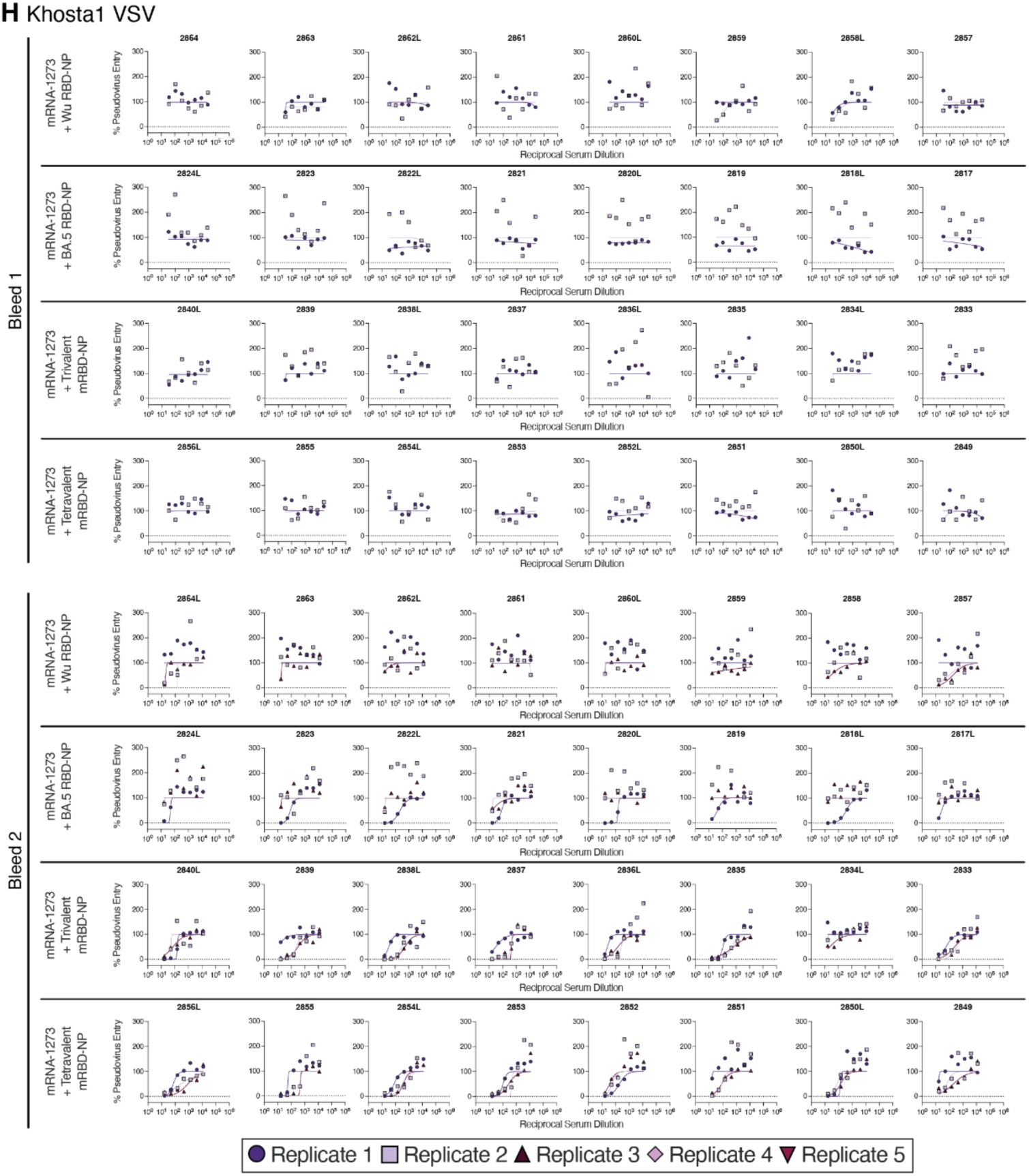
Neutralization dose-response curves for RBD-NP-immunized pre-immune mice. Serum neutralizing antibody titers were assessed using VSV pseudotyped with the **A)** SARS-CoV-2 Wu-G614, **B)** SARS-CoV-2 BA.5, **C)** SARS-CoV-2 BQ.1.1, **D)** SARS-CoV-2 XBB.1.5, **E)** SARS-CoV-1, **F)** BtKY72, **G)** PRD-0038, or **H)** Khosta1 S. One to five biological replicates were conducted using distinct batches of pseudovirus.

**Figure S3.**
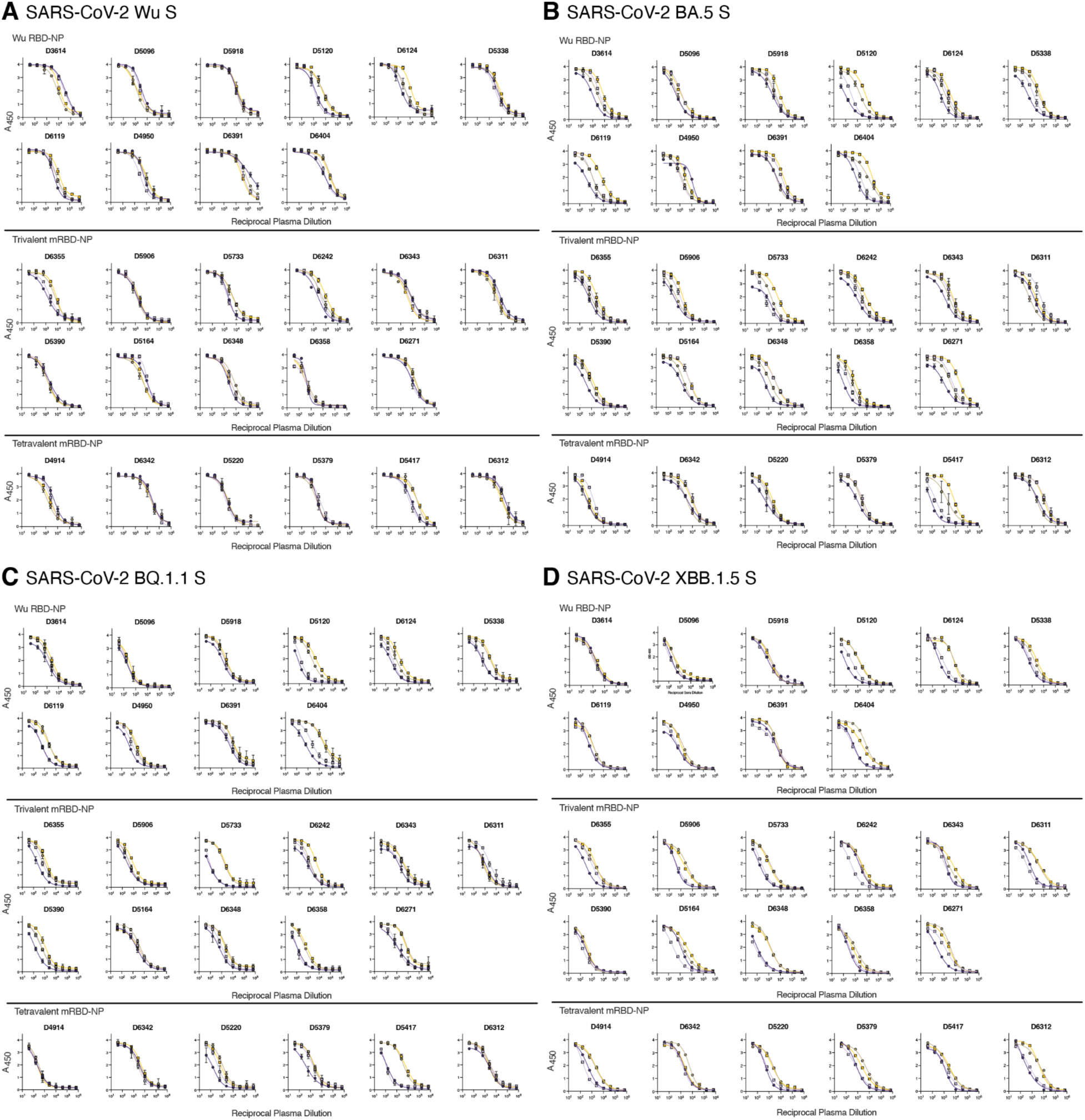
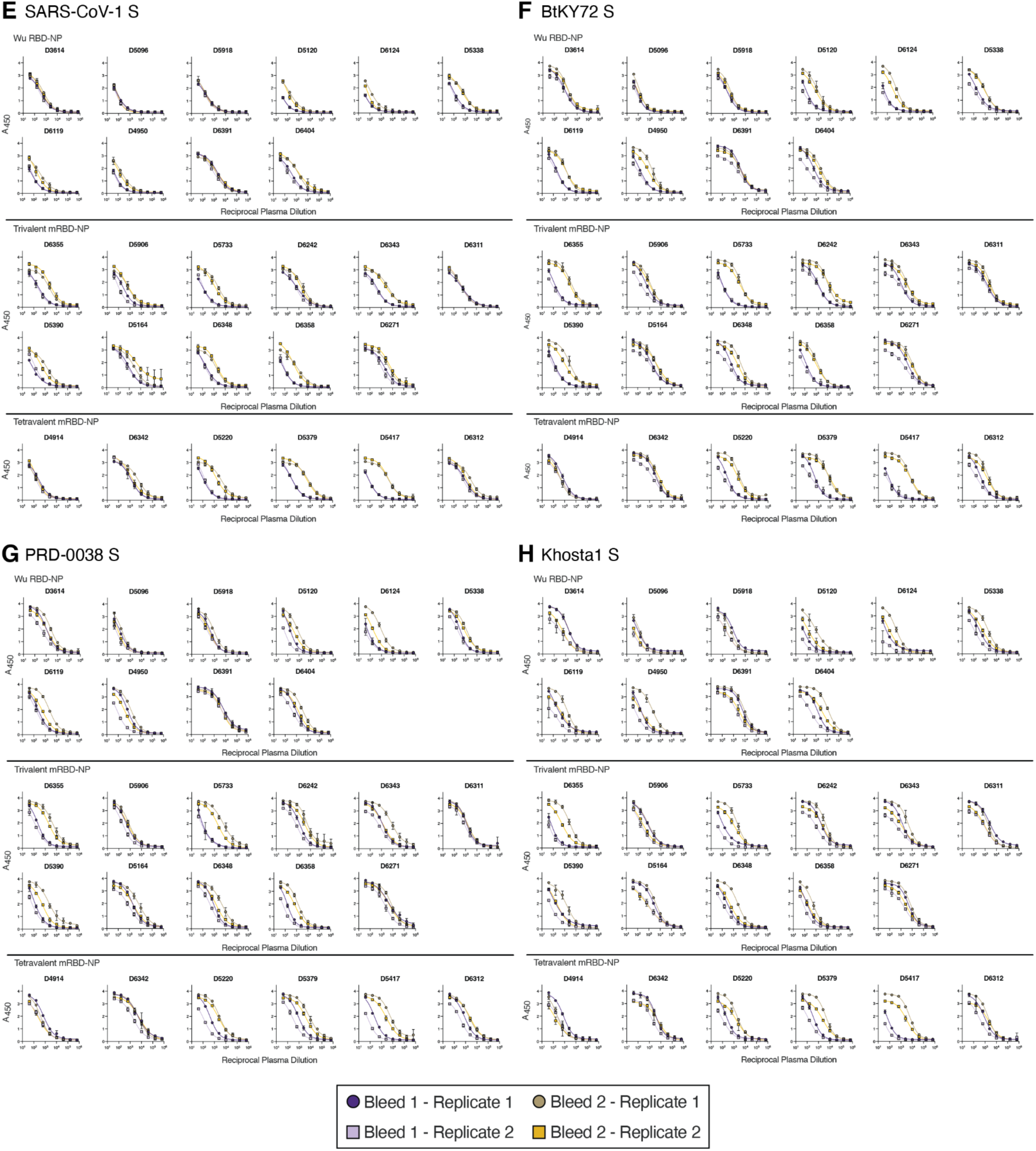
S glycoprotein ELISA dose-response curves for RBD-NP-immunized AGMs. Plasma binding antibody titers were assessed using the **A)** SARS-CoV-2 Wu, **B)** SARS-CoV-2 BA.5, **C)** SARS-CoV-2 BQ.1.1, **D)** SARS-CoV-2 XBB.1.5, **E)** SARS-CoV-1, **F)** BtKY72, **G)** PRD-0038, or **H)** Khosta1 S. Two biological replicates were each conducted in technical duplicates using distinct batches of proteins.

**Figure S4.**
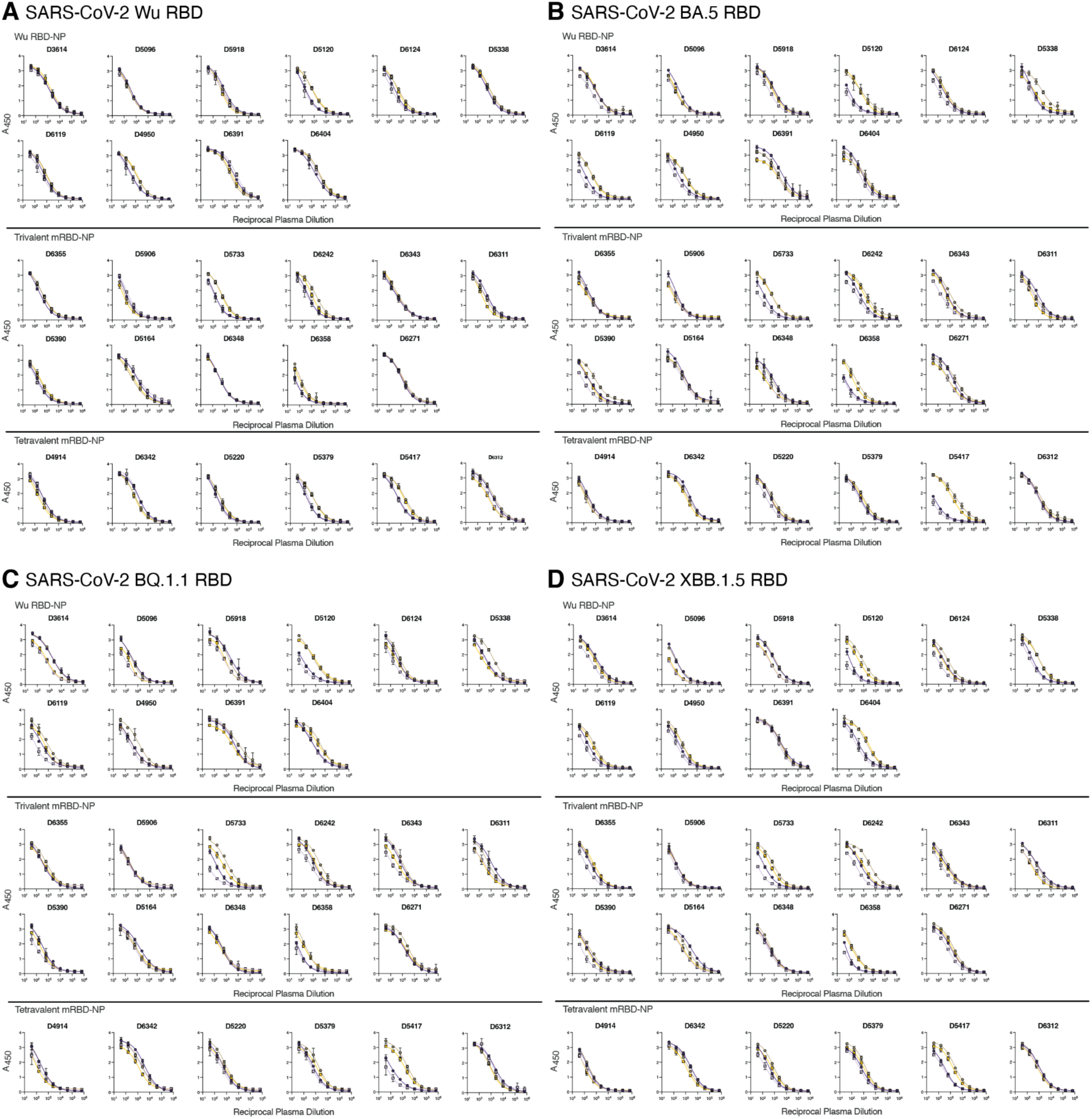
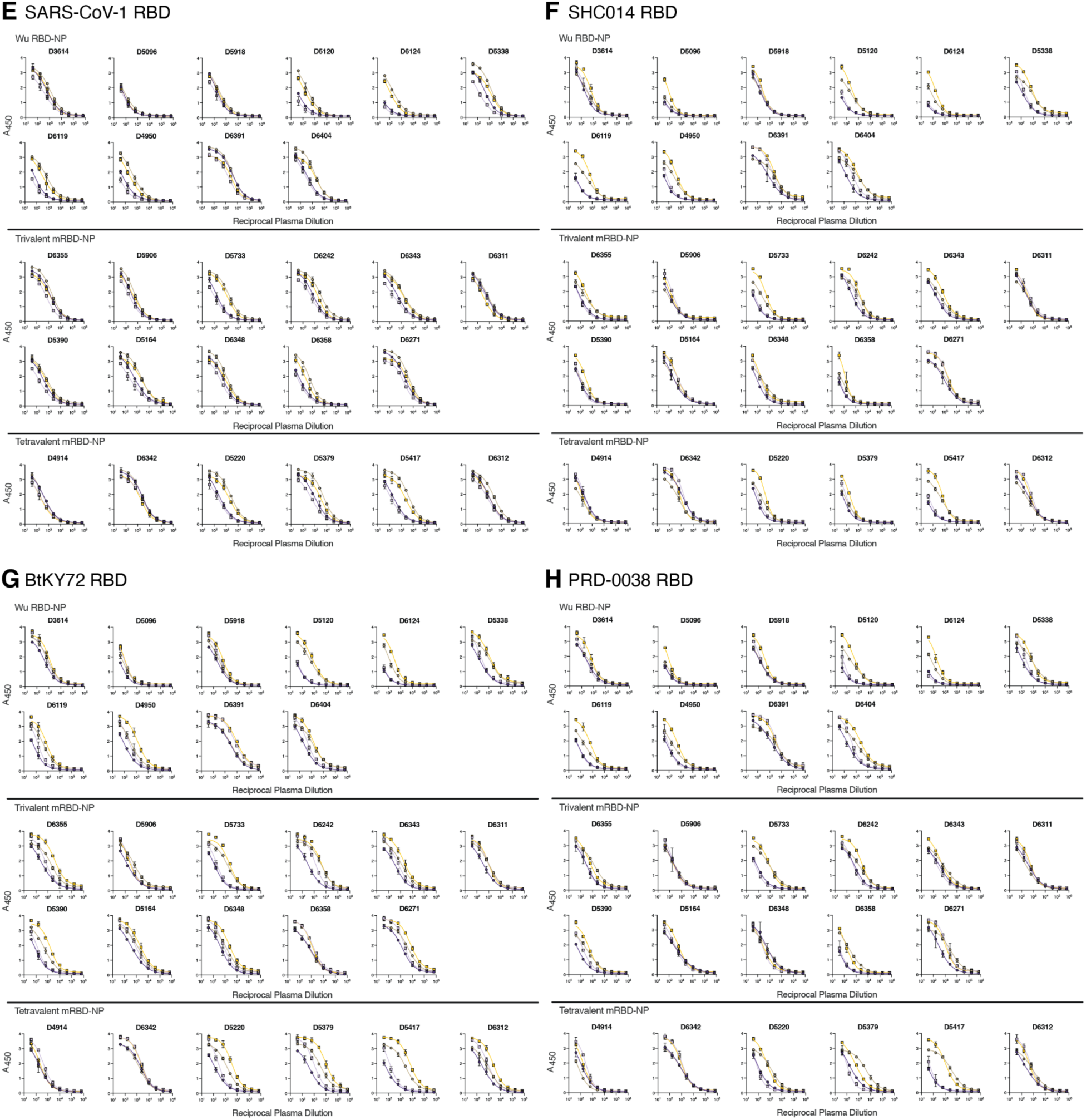
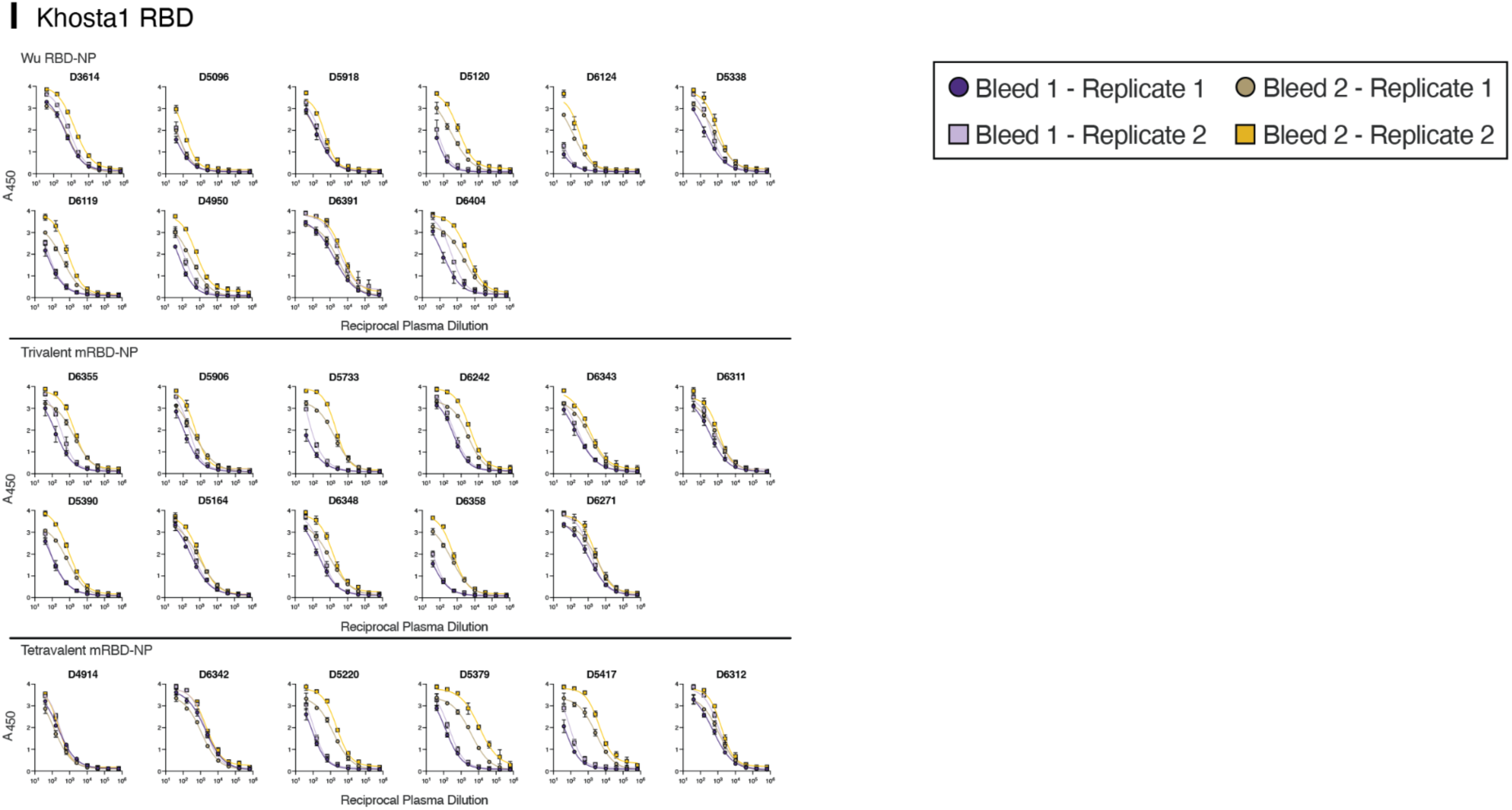
RBD ELISA dose-response curves for RBD-NP-immunized AGMs. Plasma binding antibody titers were assessed using the **A)** SARS-CoV-2 Wu, **B)** SARS-CoV-2 BA.5, **C)** SARS-CoV-2 BQ.1.1, **D)** SARS-CoV-2 XBB.1.5, **E)** SARS-CoV-1, **F)** RsSHC014, **G)** BtKY72, **H)** PRD-0038, or **I)** Khosta1 RBD. Two biological replicates were each conducted in technical duplicates using distinct batches of proteins.

**Figure S5.**
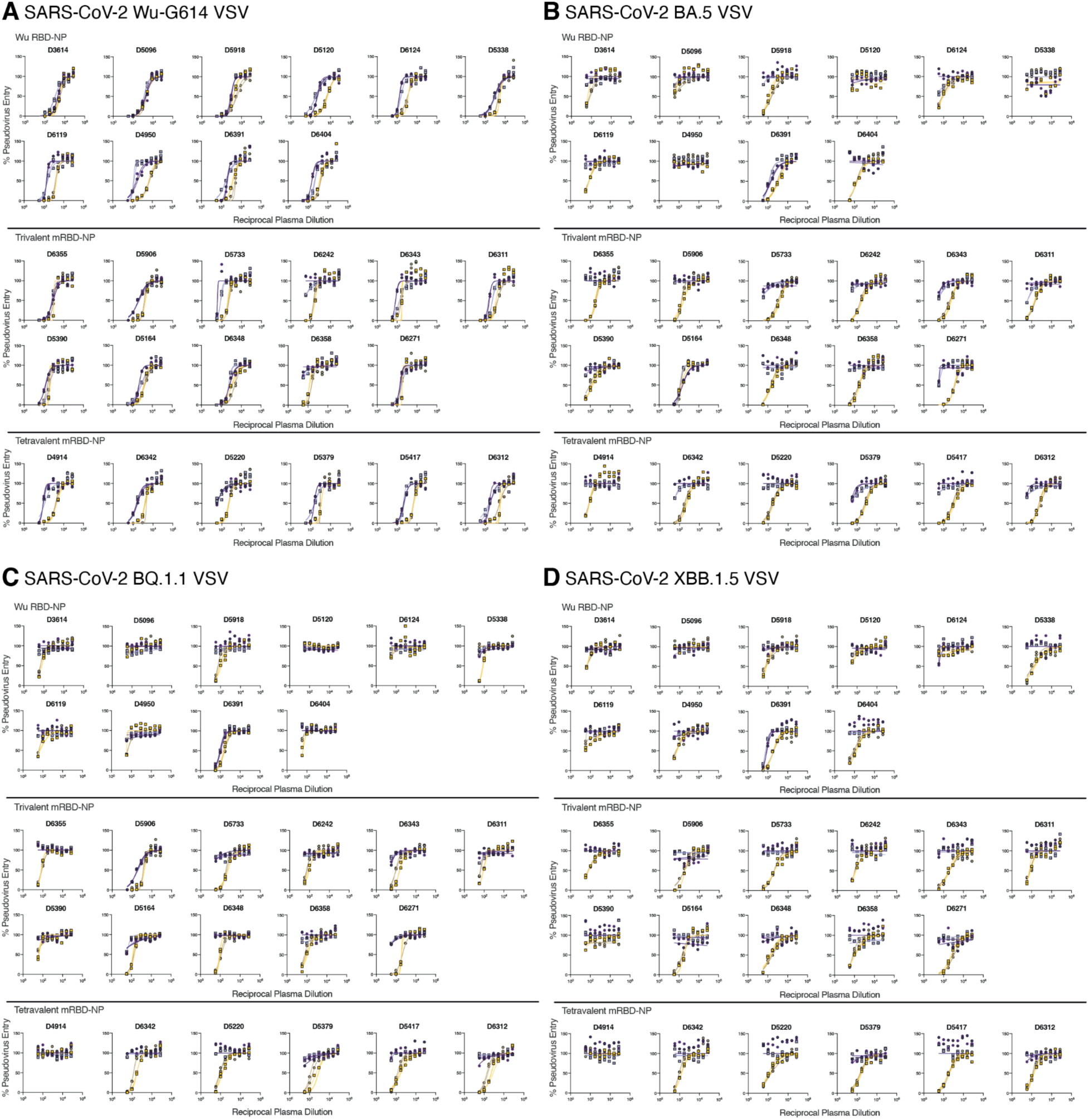
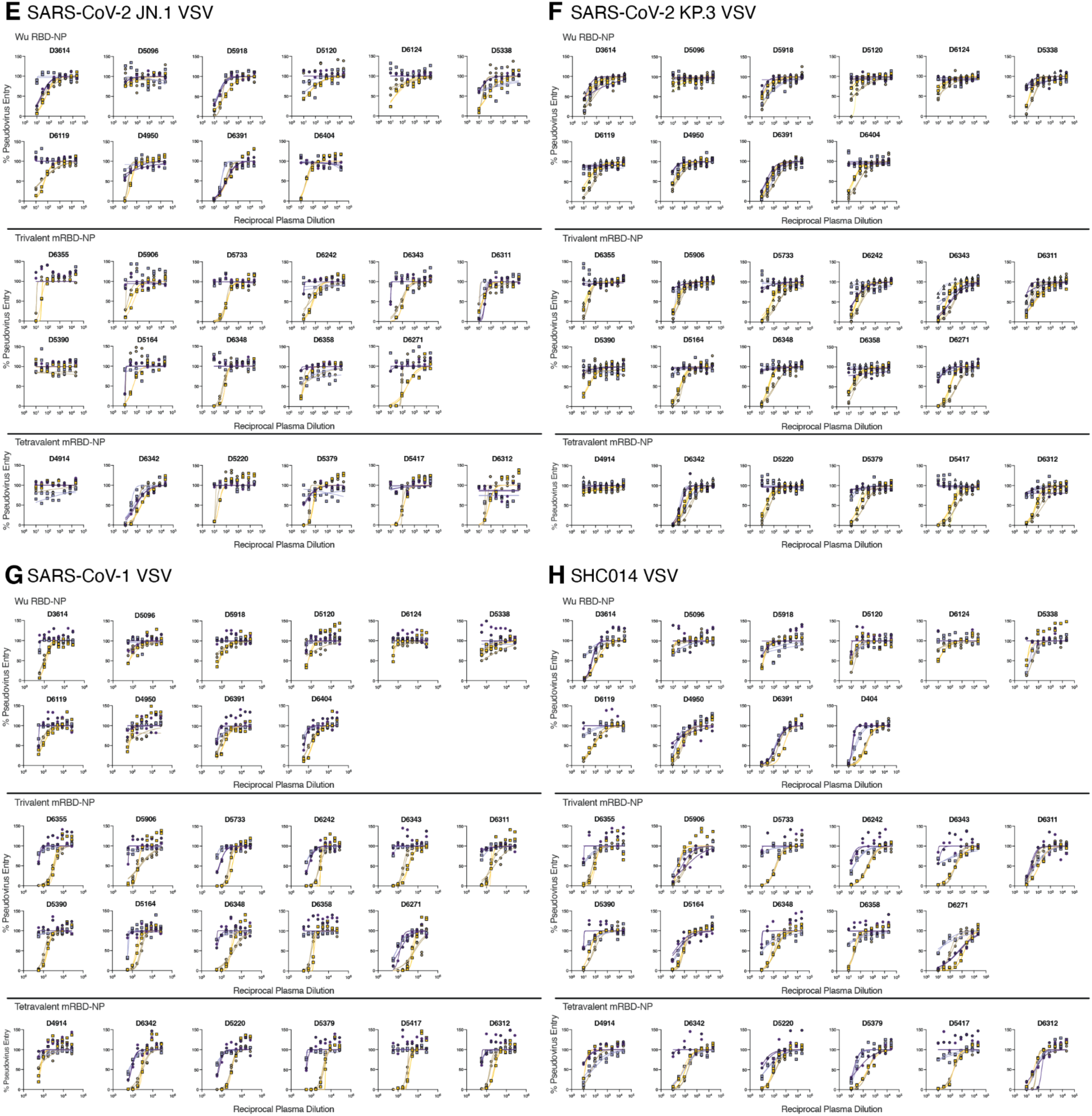
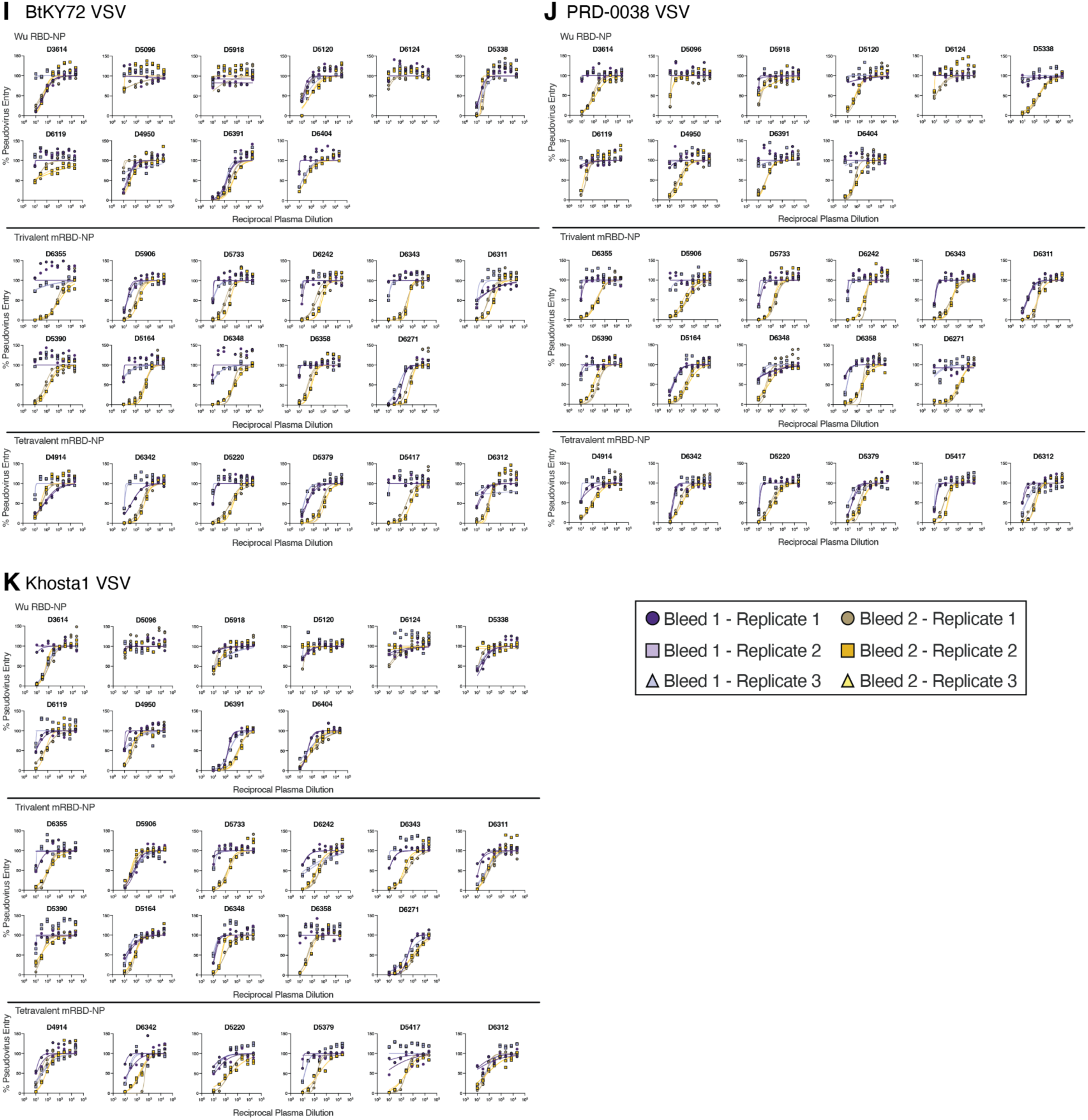
Neutralization dose-response curves for RBD-NP-immunized AGMs. Plasma neutralizing antibody titers were assessed using VSV pseudotyped with the **A)** SARS-CoV-2 Wu-G614, **B)** SARS-CoV-2 BA.5, **C)** SARS-CoV-2 BQ.1.1, **D)** SARS-CoV-2 XBB.1.5, **E)** SARS-CoV-2 JN.1, **F)** SARS-CoV-2 KP.3, **G)** SARS-CoV-1, **H)** RsSHC014, **I)** BtKY72, **J)** PRD-0038, or **K)** Khosta1 S. At least two biological replicates were conducted each in technical duplicates using distinct batches of pseudovirus.

**Figure S6.**
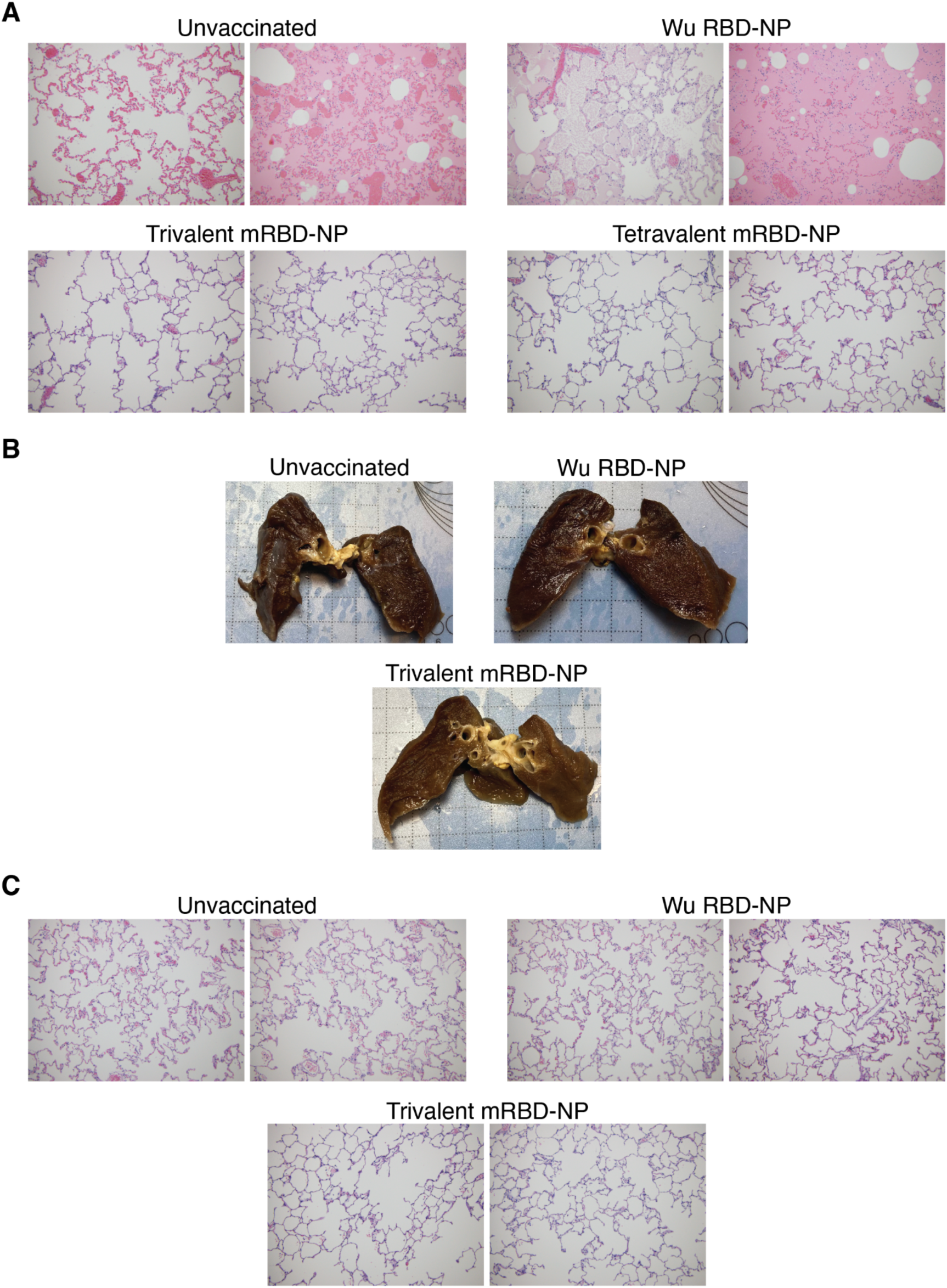
Histological analysis of lungs from SARS-CoV-2 XBB.1.5 and RsSHC014- challenged AGMs. **A)** Representative images of H&E stained lung tissue from SARS-CoV-2 XBB.1.5-challenged AGMs. **B-C)** Representative images of (B) gross lungs and (C) H&E stained lung tissue from RsSHC014-challenged AGMs.

**Figure S7.**
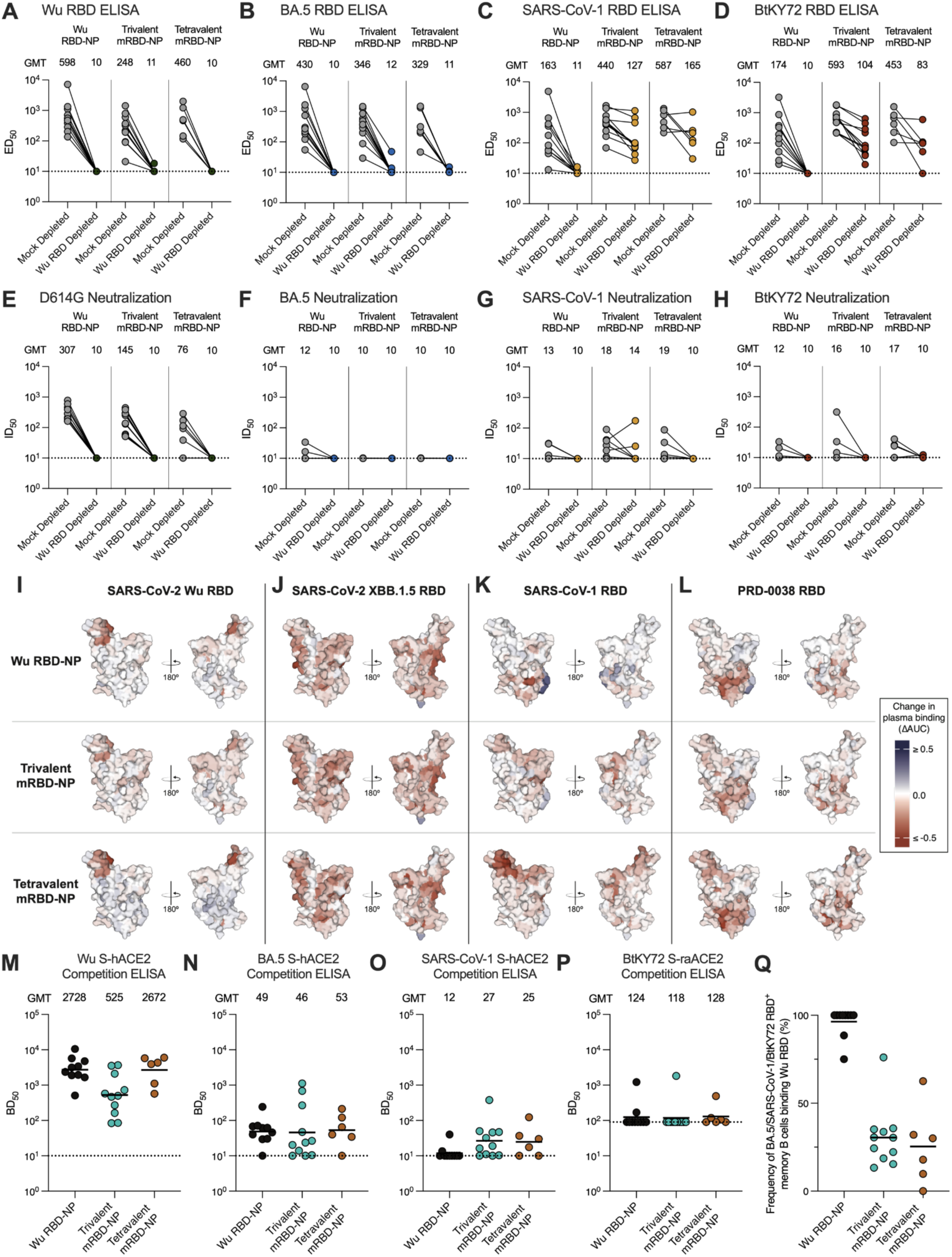
Immune imprinting following one dose of RBD NP in pre-immune AGMs. A-D) Plasma binding titers against the (A) SARS-CoV-2 Wu, (B) SARS-CoV-2 BA.5, (C) SARS-CoV-1, or (D) BtKY72 RBDs following incubation with uncoated magnetic beads (mock depletion) or Wu RBD-coated magnetic beads (Wu RBD depleted). **E-H)** Plasma neutralizing antibody titers upon mock depletion or Wu RBD depletion against VSV pseudotyped with the (E) SARS-CoV-2 Wu-G614, (F) SARS-CoV-2 BA.5, (G) SARS-CoV-1, or (H) BtKY72 S glycoproteins. Data presented reflect results obtained from one biological replicate with ELISAs and neutralization assays being conducted in technical duplicate and are representative of results obtained from two biological replicates. The geometric mean titers (GMT) are displayed above the plots. The limit of detection (ED50: 1/10 or ID50: 1/10) is represented by a dotted line. **I-L)** Escape mutations mapped for the SARS-CoV-2 Wu (I), SARS-CoV-2 Wu XBB.1.5 (J), SARS-CoV-1 (K), or PRD-0038 (L) RBDs using plasma collected after one dose of the indicated RBD-NPs. Data reflect averaged results from plasma collected from three AGMs with the highest ID50 values against XBB.1.5 in each group. Mutations at sites that increase plasma binding are represented in blue, while those that decrease plasma binding are represented in red. **M-P)** Plasma ACE2 blocking titers using the SARS-CoV-2 (M) Wu or (N) BA.5, (O) SARS-CoV-1, or (P) BtKY72 S glycoprotein. Each data point represents the BD50 for a given animal obtained by averaging two biological replicates (independently produced batches of proteins) conducted in technical duplicate. The limit of detection (BD50: 1/10 for Wu, BA.5, and SARS-CoV-1 and 1/90 for BtKY72) is represented by a dotted line. The line indicates geometric mean titers (GMTs). **Q)** Frequency of memory B cells binding to the BA.5/SARS-CoV-1/BtKY72 RBD pool that additionally recognize the Wu RBD in peripheral blood collected after one dose of NP as enumerated by flow cytometry.

**Figure S8.**
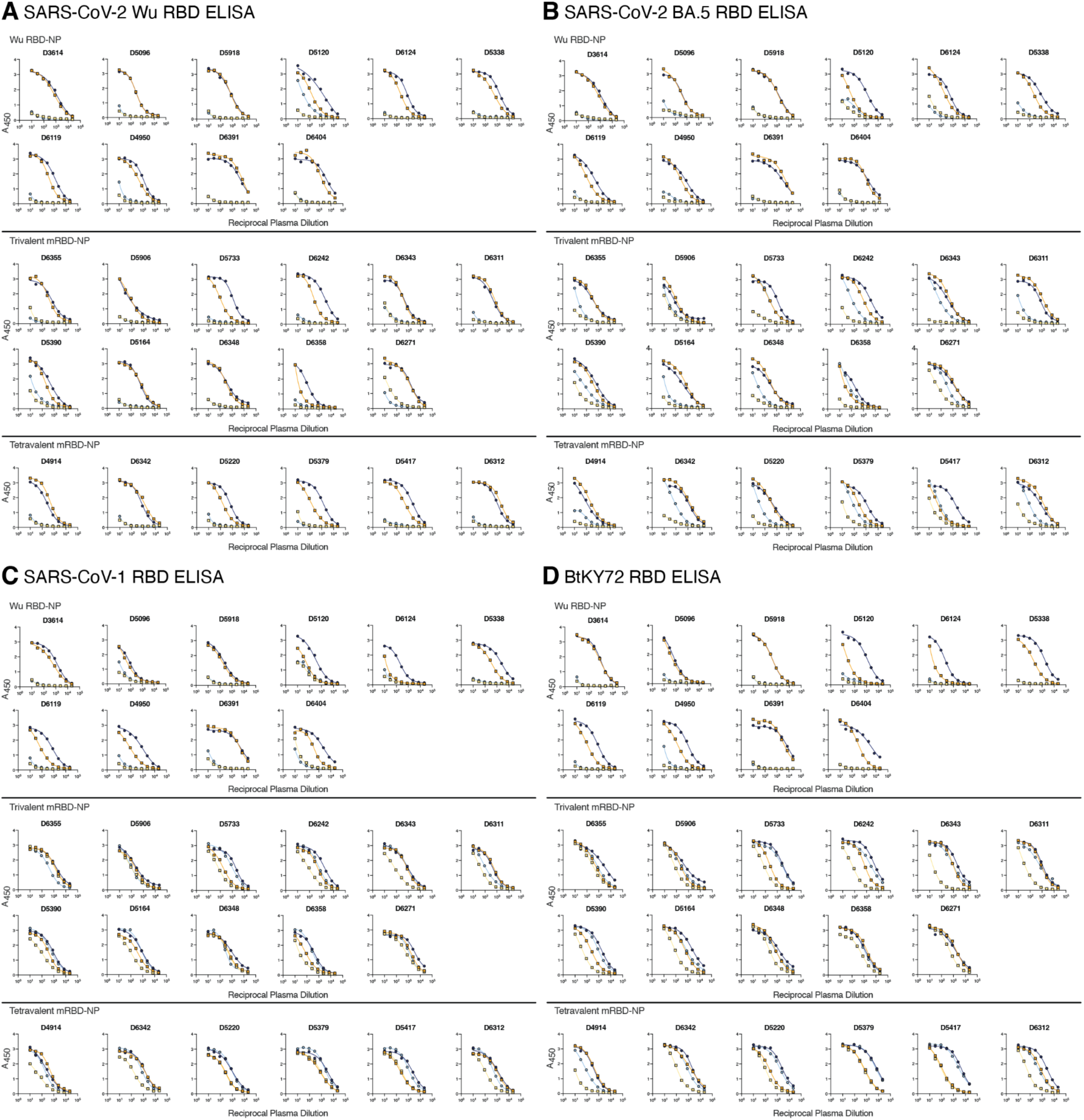
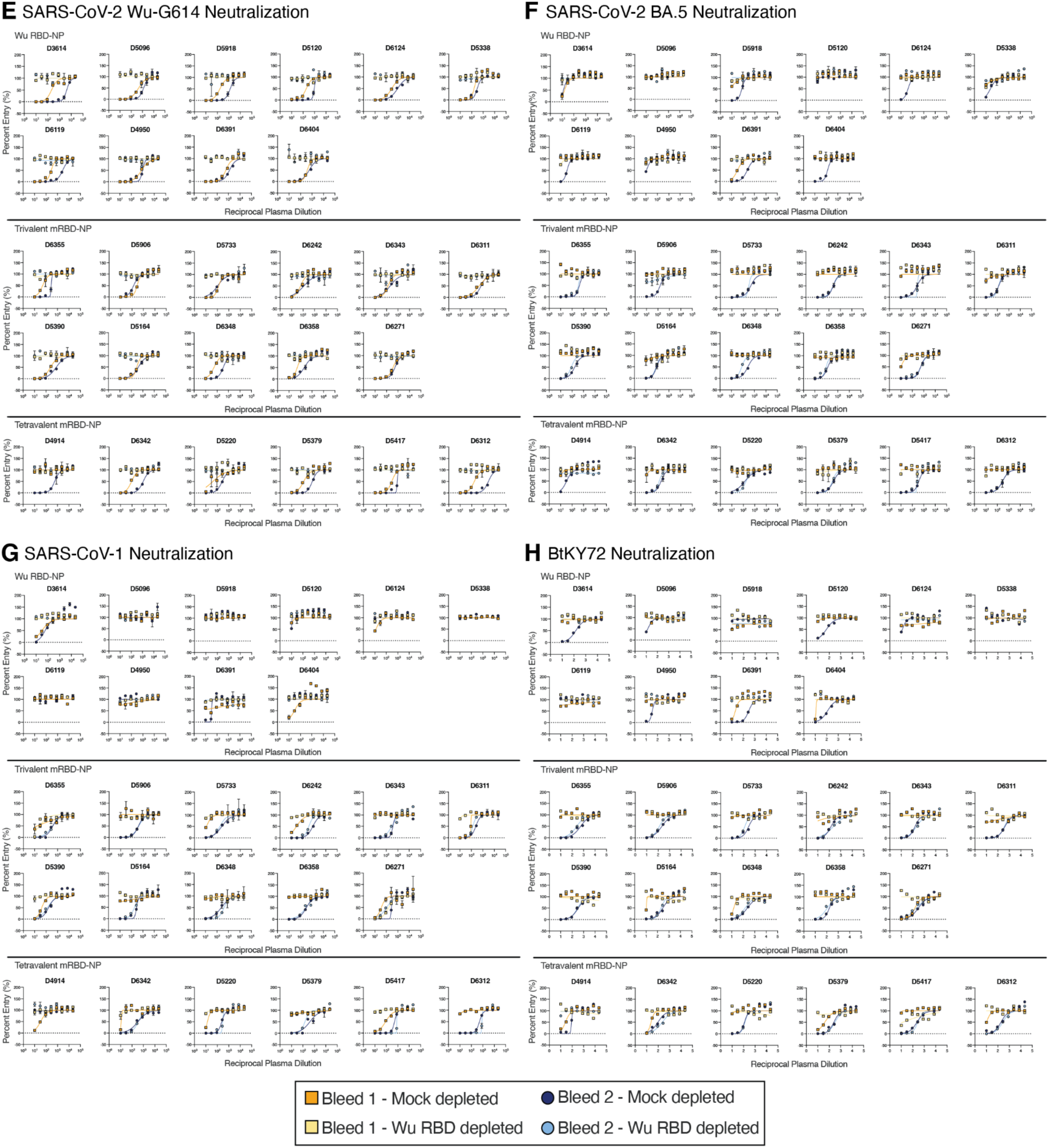
RBD ELISA AND neutralization dose-response curves for RBD-NP-immunized AGMs following mock depletion or depletion of Wu RBD-directed antibodies. Binding antibody titers for mock or Wu RBD depleted plasma were assessed using the **A)** SARS-CoV-2 Wu, **B)** SARS-CoV-2 BA.5, **C)** SARS-CoV-1, or **D)** BtKY72 RBD. Neutralizing antibody titers for mock or Wu RBD depleted plasma were assessed using VSV pseudotyped with the **E)** SARS-CoV-2 Wu, **F)** SARS-CoV-2 BA.5, **G)** SARS-CoV-1, or **H)** BtKY72 S. Data presented are from one biological replicate with both ELISAs and neutralization assays conducted in technical duplicate and representative of results from a second independent depletion experiment completed with a distinct batch of proteins and pseudoviruses.

**Figure S9.**
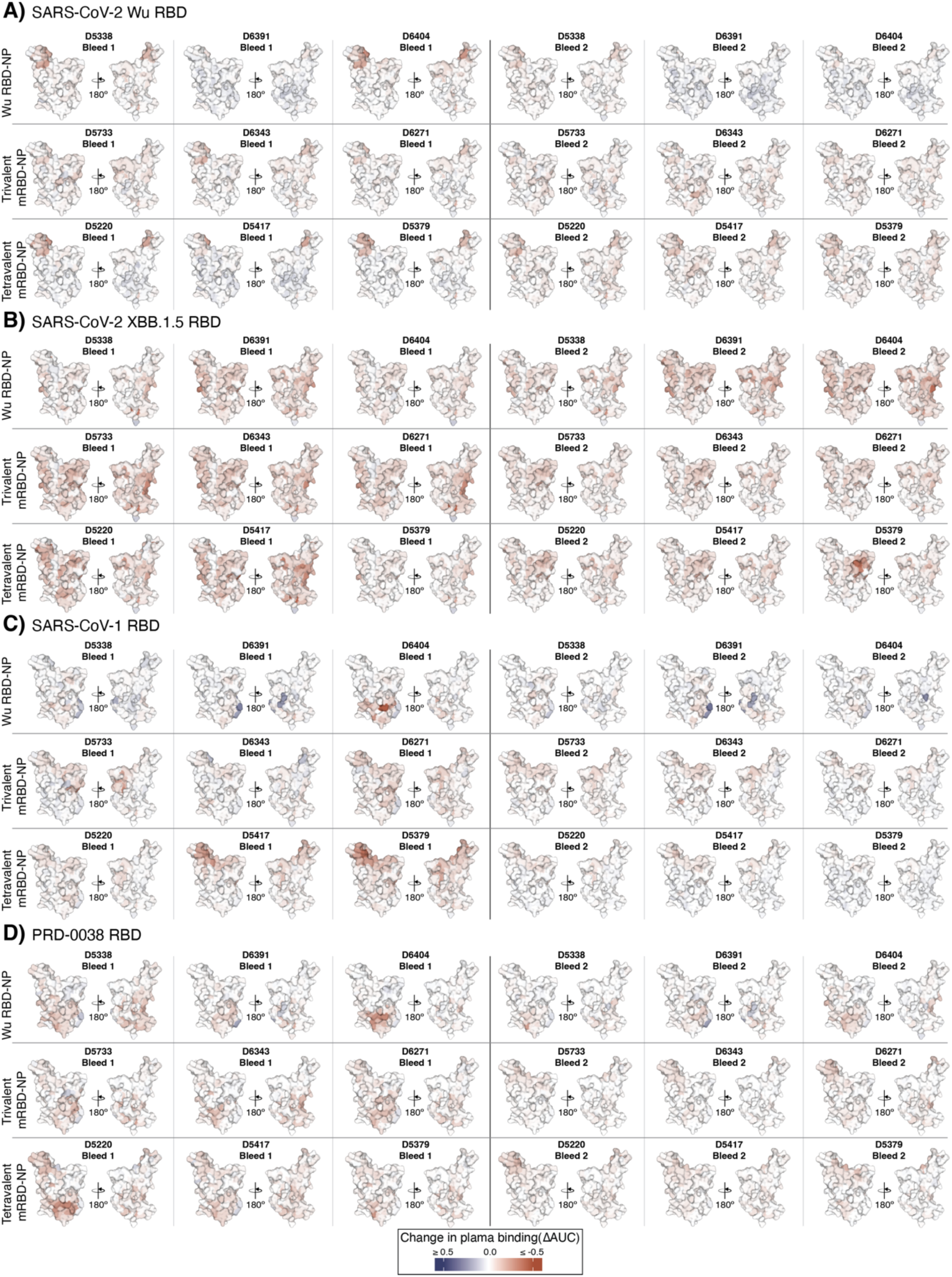
Deep mutational scanning data for RBD escape mutations for individual AGMs. Plasma escape mutations mapped for each of the 9 AGMs profiled using the **A)** SARS-CoV-2 Wu, **B)** SARS-CoV-2 XBB.1.5, **C)** SARS-CoV-1, or **D)** PRD-0038 RBD DMS libraries.

**Figure S10.**
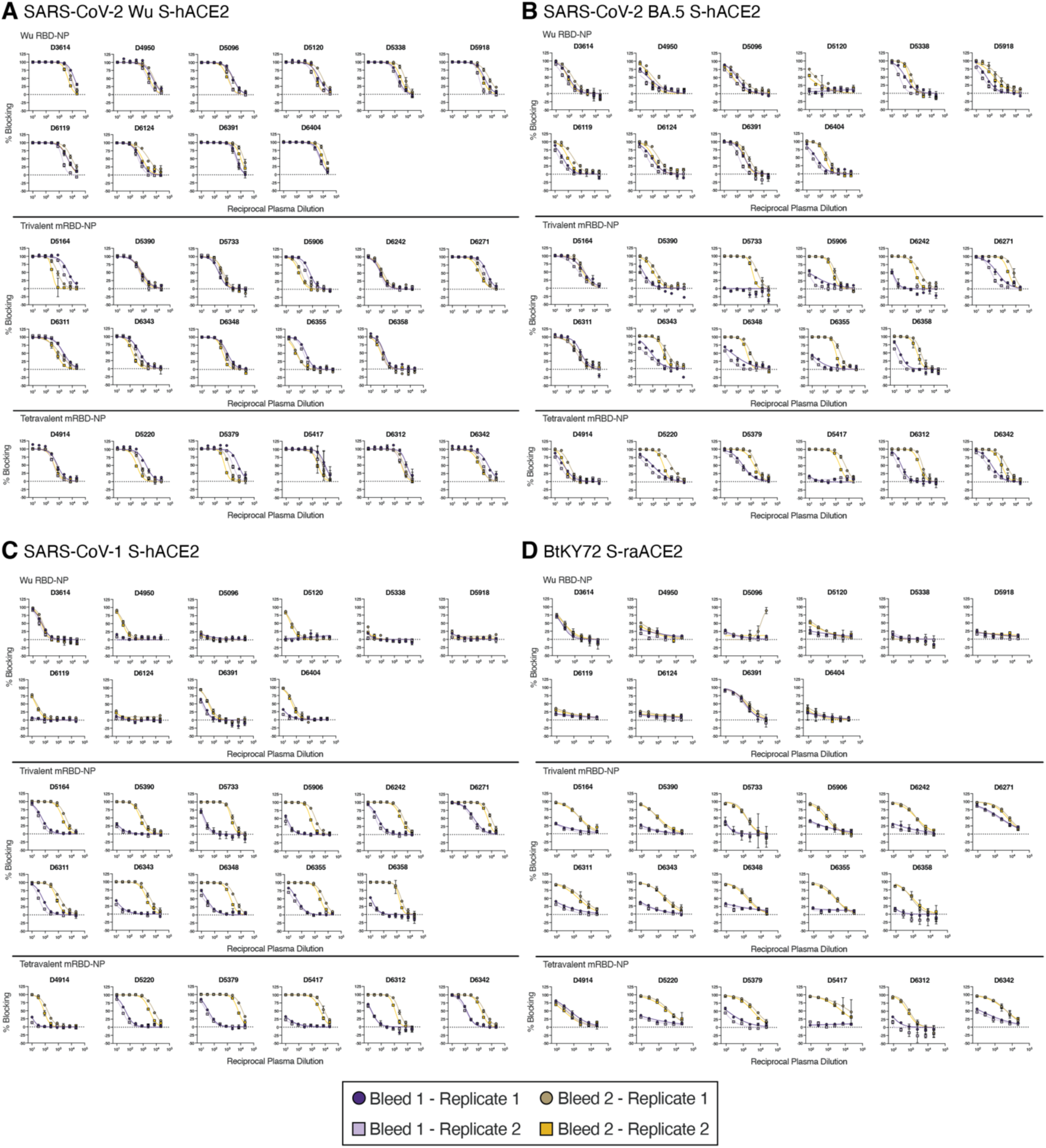
Human or *R. alcyone* ACE2 competition ELISA dose-response curves for RBD-NP-immunized AGMs. Plasma ACE2 blocking titers were assessed using the **A)** SARS-CoV-2 Wu, **B)** SARS-CoV-2 BA.5, or **C)** SARS-CoV-1 S and dimeric human ACE2 (hACE2) or **D)** BtKY72 S and dimeric *R. alcyone* ACE2 (raACE2). Two biological replicates were each conducted in technical duplicates using distinct batches of proteins.

**Figure S11.**
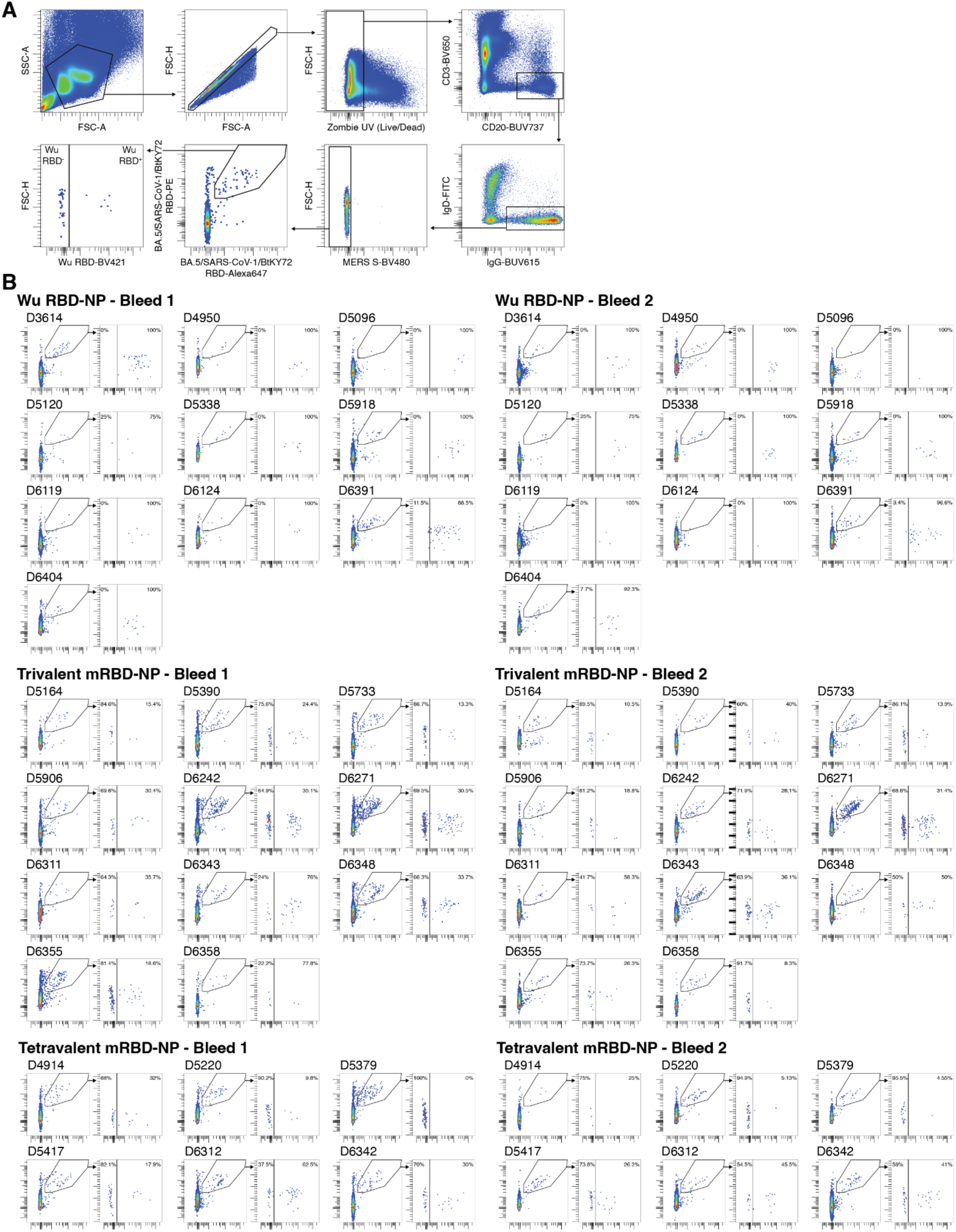
Flow cytometry analysis of memory B cells collected from RBD-NP-immunized AGMs. **A)** Gating strategy to determine binding specificity of memory B cells in peripheral blood collected from RBD-NP-immunized AGMs. **B)** Evaluation of RBD-reactivity of memory B cells for each individual AGM.

## Notes

### Competing Interest Statement

A.C.W., M.C.M., N.P.K., and D.V. are named as inventors on patent applications filed by the University of Washington related to coronavirus vaccines. The King lab has received unrelated sponsored research agreements from Pfizer and GSK.

